# Heterozygous transcriptional signatures unmask variable premature termination codon (PTC) burden alongside pathway-specific adaptations in blood outgrowth endothelial cells from patients with nonsense DNA variants causing hereditary hemorrhagic telangiectasia

**DOI:** 10.1101/2021.12.05.471269

**Authors:** Maria E Bernabeu-Herrero, Dilip Patel, Adrianna Bielowka, Patricia Chaves Guerrero, Stefan J Marciniak, Michela Noseda, Micheala A. Aldred, Claire L Shovlin

## Abstract

Frameshift and nonsense DNA variants represent the commonest causes of monogenic inherited diseases. They usually generate premature termination codon (PTC)-containing RNA transcripts that produce truncated proteins in recombinant systems, but endogenously are subject to nonsense mediated decay. To examine native consequences of these variants, we derived cells from pre-genotyped patients. Blood outgrowth endothelial cells (BOECs) were established from individuals with hereditary hemorrhagic telangiectasia (HHT) due to a heterozygous nonsense variant in *ACVRL1*, *ENG* or *SMAD4* that each encode an endothelial cell-expressed protein mediating bone morphogenetic protein (BMP)/ transforming growth factor (TGF)-β signalling. RNA sequencing alignments to PTC alleles varied from 8-23% of expected, and differed between same-donor replicates. Differential gene expression analyses were validated by single cell qRT-PCR, and identification of changes in wider, disease-specific BMP/TGF-β pathway components. However, the most differentially expressed genes clustered to disease-independent terms for post translational protein modification (isopeptide bond; ubiquitin-like conjugation). They were the only terms meeting Benjamini significance after clustering Bonferroni-ranked, differentially expressed genes from the 5,013 meeting 10% intraassay coefficients of variation, and significance was robust to normalisation methods. Optimised pulse chase experiments supported perturbed wildtype protein maturation, but no PTC-truncated protein was identified. Unexpectedly, BOEC cultures with highest PTC persistence were discriminated in unsupervised hierarchical clustering of low GINI coefficient ‘invariant’ housekeeper genes, and patterns were compatible with higher cellular stress. The findings support a model whereby PTCs are more of a burden in stressed cells, and lead us to conclude that overlooked and varying PTC burdens contribute to biological variability.

## INTRODUCTION

Mechanistic understanding, and efficacious preclinical testing of potentially new therapeutic agents require robust laboratory assays, particularly for rare diseases, and genome-personalised medicine when sample size is limiting. The current decade benefits from global ‘omics approaches that provide prior evidence in normal states,^1,2^ and avoid over-targeted experimental readouts that can miss unanticipated adaptations employed by cells in order to survive, including therapy-nullifying responses.^3^ Evaluation platforms now include patient-derived cells for bulk, single cell, and organ-on-a-CHIP analyses.

Our goals were to deliver methodologies for pre-clinical studies in patient-derived blood outgrowth endothelial cells (BOECs), incorporating understanding of the extent to which disease-causing RNA transcripts harbouring premature termination codons (PTCs) are lost in normal cellular safeguarding processes such as nonsense mediated decay (NMD).^4-6^ Currently, PubMed searches for “nonsense mediated decay” retrieves <3% of the number for “truncated protein” search terms, while “truncating variant” (not a recommended descriptor^7,8^), is still widely used to describe DNA variants in manuscripts and clinical reports that imply truncated proteins are generated in endogenous tissues, as in recombinant *in vitro* studies.

We focussed on patients with the vascular dysplasia hereditary hemorrhagic telangiectasia (HHT) where new treatments are required to further reduce morbidity from hemorrhage caused by abnormal blood vessels.^9,10^ HHT is inherited as an autosomal dominant trait, and typically caused by heterozygous, loss-of-function variants in *ACVRL1, ENG* and *SMAD4*.^11-13^ The most common variants are PTC-generating such as frameshift and nonsense (‘stop gain’), with hundreds of unique pathogenic PTC variants together responsible for 42-67% of HHT cases, according to causal gene.^14^ While a small number of HHT cells may lose the second wildtype allele,^15^ in the majority, it is retained, even in abnormal vessels where approximately half normal ENG protein expression was demonstrated in *ENG^+/-^* HHT patients.^16-18^ For the disease-causing allele, there is evidence in HHT patient-derived cells that the transcript may be lost at the RNA level (*ENG* nonsense variants in EBV-transformed lymphoblastoid cell lines^19^), or generate only short-lived, endoplasmic reticulum-retained proteins (*ENG* and *ACVRL1* multiexon deletion and missense alleles^16-18,20-23^). There has been limited attention on how these processes operate alongside pathway-specific perturbations in the heterozygous state.

Pathway-specific perturbations in HHT have been reviewed recently.^12,13^ *ACVRL1, ENG* and *SMAD4* encode endothelial-cell expressed proteins that transmit or modify signaling by the bone morphogenetic protein (BMP)/ transforming growth factor (TGF)-β superfamily, with HHT causality confirmed by null, heterozygous and endothelial-specific disease gene models generated using recombinant technology.^24-28^ Early heterozygous models were recognised as excellent simulations of the human condition^24,29^ but the same variability and often mild phenotypes emphasised the importance of a second and even third “hit” in addition to the heterozygous state.^30^ Signaling defects are usually studied in conditional homozygous null models which generate pronounced and reproducible phenotypes, for example in the context of trauma^25^ or angiogenesis.^31,32^These have provided important insights, including modified BMP/TGF-β receptor expression interpreted as an adaptive response that enabled more *ENG* deficient cells to survive.^32^

To examine native consequences of heterozygous alleles that generate PTCs, we generated blood outgrowth endothelial cells (BOECs) from pre-genotyped HHT donors and controls. Here we report marked variability in the persistence of nonsense pathogenic variants in replicate, resting state cultures from human heterozygous HHT donors; associated changes in protein maturation kinetics; and independent, unsupervised RNAseq discrimination of BOECs cultures with higher PTC burdens.

## METHODS

### Endothelial cell methods

This research was approved by national ethics committees, and all human participants provided written informed consent. Primary studies were performed in BOECs established from healthy volunteers and pre-genotyped HHT patients, using previously optimised methods.^33-35^ All analyses were performed in confluent passage 3 vials as described further in the Data Supplement.

For RNASeq library preparations, RNA was extracted from 16 separate cultures of confluent BOECs from 4 separate control and HHT donors, cultured in the presence and absence of BMP9 at 10ng/ml for 1 hr. Library preparations, alignments to *Homo sapiens* GRCh38 (STAR aligner v2.5.2b), counts of unique gene reads that fell within exon regions (Subread package v1.5.2), normalisation to total alignment counts per library, and initial variant calls^36^ were performed by Genewiz (Leipzig, Germany).

To provide numeric validations of a subset of genes, single cell (sc) qRT-PCR was performed as detailed further in the Data Supplement:^37,38^ For each culture, 40 viable (DRAQ7 negative) single cells were sorted directly into 96-well-plates containing pre-amplification mix for 48 genes selected either as endothelial gene markers as validation for endothelial differentiation, or likely relevance to the HHT cellular phenotype, or as important negative controls (Supplementary Table 1). Specific target amplification was performed using CellDirect One-Step qRT-PCR Kit (Invitrogen, Thermo Fisher, Waltham, MA) as described.^37,38^ The 48 genes examined by sc qRT-PCR were normalized to *UBC* expression, analysed, and plotted using an R script developed in house. Novel housekeeping genes were sought from genes with low Gini Coefficients (GC) within multiple cell lines.^39,40^

To enhance the technological feasibility of detecting endogenous low level, truncated proteins, metabolic labelling was performed in BOEC protein extracts using ENG mAb SN6h (Dako Denmark A/S, Denmark). Modifying methods of Pece et al,^18^ a 1 hour “pulse” with 50μCi/ml of ^35^S methionine, preceded a 0, 1, 2 or 3.5hr “chase” in media containing unlabelled methionine, immunoprecipitation and SDS-PAGE gel fractionation.

### RNA Sequencing Analyses

Percentage loss of the nonsense allele in *ACVRL1^+/-^, ENG^+/-^* and *SMAD4^+/-^*BOECs was calculated assuming that without nonsense mediated decay (NMD),^4-6^ there would have been an approximately equal number of alignments to wildtype and nonsense alleles in each HHT BOEC library.

Across all 16,807 genes, the intra-assay coefficient of variation (CV),^41^ *100 x standard deviation (SD)/mean*, was calculated for replicate pairs, adjusting the alignment per gene for total library read counts, and restricting to genes where all four untreated replicate pairs had an intra-assay coefficient of variation <10% (‘met CV10’). The 25 genes with GINI Coefficients (GCs)<0.15 in diverse cells, i.e. representing the least variable of human transcripts,^39,40^ were tested for BOEC expression and used as housekeepers for DeSeq2 normalization.^42^ This calculated the ratio of alignment counts for each selected GINI housekeeper gene in each BOEC dataset to the geometric mean of that GINI gene across all 16 BOEC datasets. The median value of GINI gene ratios in each BOEC library was then used to generate the ‘size factor’ to scale that library’s alignments.^42,43^

Gene Ontology^44,45^(GO) and pathway enrichment analysis of HHT differentially expressed genes was performed using Functional Annotation Clustering through the Database for Annotation, Visualization and Integrated Discovery (DAVID)^46^v6.8. Primary clustering analyses were on the most differentially expressed genes meeting CV10^41^ in all four untreated replicate pairs of BOECs, and where the gene’s differential expression between control and HHT BOECs met either Bonferroni p<0.05, or Bonferroni p<1.00. GO and pathway terms clustered by these genes were considered significant if the individual pathway terms met Benjamini p<0.05. To test for robustness of normalization methods and specificity, findings were compared to all gene ontology clusters generated by genes ranked without GINI Coefficient housekeeper gene normalization, and to all clusters generated by 10 sets of 1,000 genes randomly derived from the complete dataset of gene alignments in the BOECs.

### Data Analysis

STATA IC v 15.0 (Statacorp, College Station, TX) was used for data handling including calculation of summary statistics, comparison of groups using Kruskal Wallis when Dunn’s test was used to derive pairwise comparisons; linear regression, unsupervised heirachical clustering, and principal component analyses (PCA). Data were visualised using GraphPad Prism 9 (GraphPad Software, San Diego, CA), and STATA IC v 15.0 (StataCorp., College Station, TX).

## RESULTS

### Establishing BOEC platforms

To examine native consequences of PTC-generating variants, we cultured primary blood outgrowth endothelial cells (BOECs) from pre-genotyped individuals with known disease-causal nonsense variants (HHT patients) and healthy volunteers. BOEC establishment rates were independent of the number of peripheral blood mononuclear cells plated; time to first colony appearance; HHT versus control status; and *ACVRL1^+/^* ^PTC^ versus *ENG^+/^*^PTC^ donor (all p-values >0.22, Supplementary Fig. 1). BOECs were confirmed as CD31^+^, CD44^+^, CD105^+^, CD90^-^ by bulk flow cytometric evaluations (Supplementary Fig. 2A) and displayed consistently high expression of a panel of endothelial marker genes evaluated by single cell (sc) qRT-PCR (Supplementary Fig. 2B, Supplementary Table 1).

BOECs were used to derive libraries for RNA sequencing (RNASeq), yielding reads of consistently high quality (Supplementary Fig. 3). RNASeq methodology was validated by same-donor control BOECs of identical passage but defrosted and cultured separately for single cell (sc) qRT-PCR experiments: For 48 genes examined using both methodologies, mean expression across 20 single control BOECs mirrored mean bulk RNASeq alignments in replicate control BOEC cultures (Supplementary Fig. 2C). Overall, 59% of the variability in the mean control BOEC RNASeq alignments normalized to total library count was accounted for by sc qRT-PCR ranking expression (p<0.0001).

### BOEC identity supported by pathway-specific gene analyses

Initial global unsupervised analytics were performed by Genewiz blinded to RNA origin, and based on alignments normalized to total library reads. Euclidean distances (Fig. 1A) and principal component analyses (PCA, Fig 1B) distinguished clusters of BOECs, and on unblinding these were seen to be distinguished by genotype. In keeping with the expected loss due to nonsense mediated decay,^4-6^ HHT BOECs demonstrated reduced overall alignments to the gene harbouring the mutant PTC allele (Fig. 1C). There were also trends for *ACVRL1* alignments to be low in *ENG*^+/PTC^ BOECs, and for *ENG* alignments to be low in *ACVRL1*^+/PTC^ BOECs, although these did not reach statistical significance.

**Fig 1.**
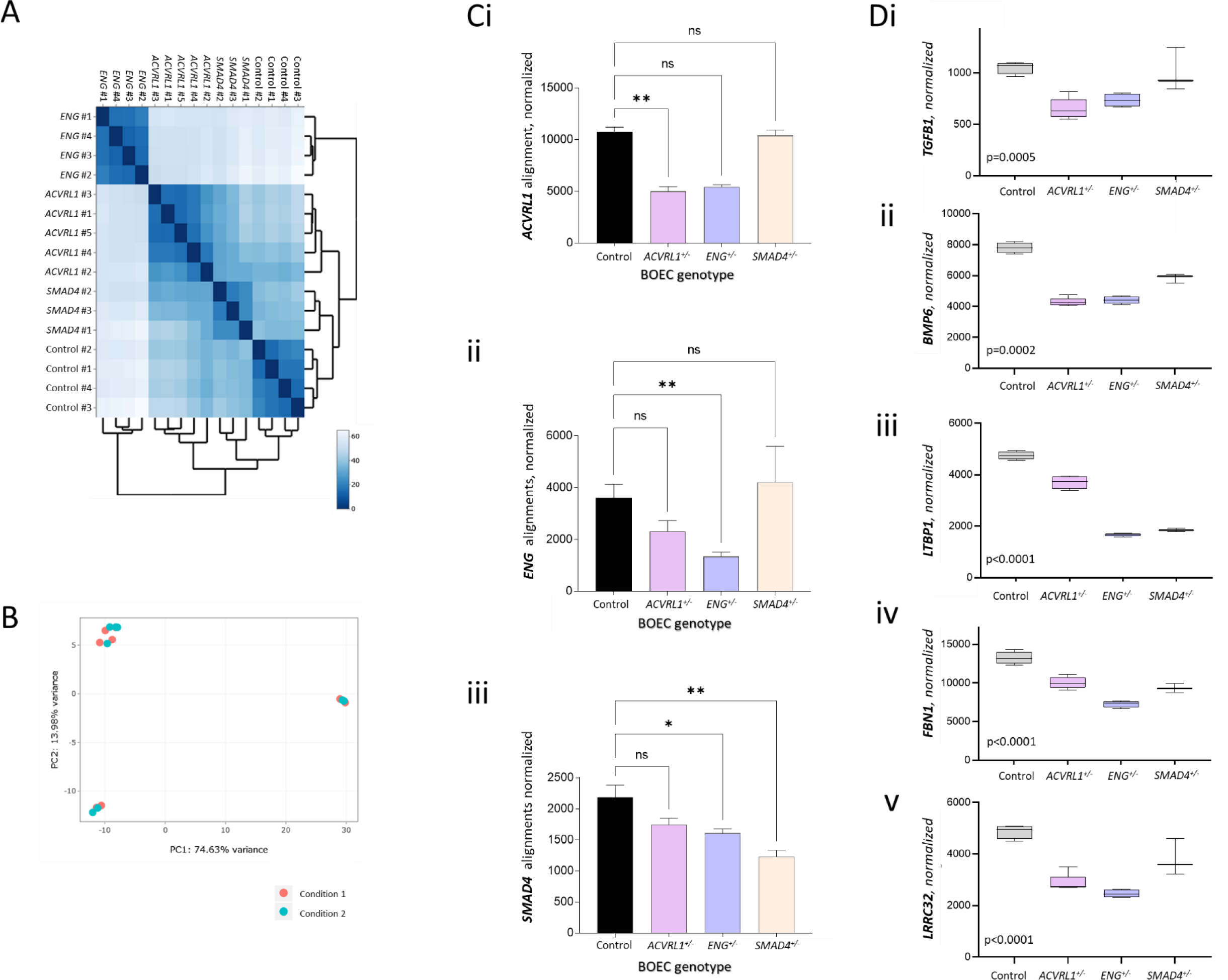
Global and disease pathway RNASeq findings in BOECs. **A)** Unsupervised Euclidean distance analyses for the 16 BOEC RNASeq datasets. When unblinded, donor replicates had clustered together. Note the cluster of 4 BOECs distinguished by greatest Euclidian distances were from genotype *ENG* ^+/PTC^. **B)** Unsupervised principal component analysis of the 16 BOEC RNASeq datasets based on the distance matrix where samples were projected to a 2D plane spanned by their first two principal components. An experimental covariate (10ng/ml BMP9 for 1 hour) was discernible only in control BOECs (top left). **C)** Normalized alignments to HHT gene transcripts ENG (ENSG00000106991), ACVRL1 (ENSG00000139567), and SMAD4 (ENSG00000141646) in control and HHT BOECs. Data from all samples per donor, error bars indicate mean and standard deviation,**p<0.005 by Dunn’s test post Kruskal Wallis. **D)** Expression of genes encoding TGF-β/BMP ligands [**i)** *TGF-*β1, **ii)** *BMP6*], and extracellular matrix BMP/TGF-β sequestration and activation proteins [**iii)** *LTBP1,* **iv)** *FBN1*, **iv)** *POSTN* and **v)** *LRRC32*]. Expression was DeSeq2-normalized^42^ using low GINI coefficient genes^39,40^ that displayed least variability in BOECs (Supplementary Table 2, Supplementary Figure 4). Box plots show interquartile range, error bars minimum to maximum. P-values were calculated by Kruskal Wallis.

Individual genes encoding other BMP and TGF-β signalling pathway components ranked highly in the genes differentially expressed between HHT and control BOECs, whether normalized to total library reads, or using more stringent analyses that restricted to 5,013 genes meeting an intra-assay coefficient of variation of 10% (CV10),^41^ and employing DeSeq2 normalization^42^ using low Gini Coefficient (GC<0.15^39,40^) housekeeper genes. Examples of differentially expressed genes are provided in Fig. 1D, including those encoding superfamily ligands (*TGFB1* Fig. 1Di, *BMP6* Fig. 1Dii), and extracellular matrix proteins involved in TGF-β/BMP ligand activation (latent transforming growth factor beta binding protein 1 (*LTBP1*, Fig. 1Diii), fibrillin 1 (*FBN1* Fig. 1Div), and leucine rich repeat containing 32 (*LRRC32,* Fig. 1Dv) which were all lower in the HHT BOECs than controls.

Taken together, the findings supported the HHT genotypic BOEC origin, with evidence suggesting wider changes in transcripts for the BMP/TGF-β signaling pathways following heterozygous loss of ACVRL1, ENG or SMAD4 function.

### Gene ontology and pathway term clustering identifies differences in protein processing pathways in HHT BOECs

To explore differences beyond the specific pathways impacted by loss-of-function variants, unbiased gene ontology clustering^44-46^ was performed on differentially expressed genes in the 5,013 meeting CV10.^41^ To examine at the highest stringency (‘Stringency Level 1’), we restricted analyses to the 30 genes (0.6% of the 5,013 CV10 genes) that met Bonferroni p<0.05 for differences between untreated control and HHT BOECs (Supplementary Table 3A). Only one cluster contained terms meeting predesignated significance. These terms were Uniprot Keywords^47^ (KW)-1017 isopeptide-bond (9 genes, Benjamini p=0.034), and KW-0832 Ubl conjugation (10 genes involved in ubiquitin-like (Ubl) conjugation, Benjamini p=0.034, Fig. 2Ai). Ten of the 30 (33%) most differentially expressed genes meeting Bonferroni p<0.05 mapped to these terms, reflecting the overlapping biology (ubiquitin and ubiquitin-like (Ubl) peptides bind to protein substrates via isopeptide bonds to target for lysosomal degradation). For Stringency Level 2, the 161 genes that met Bonferroni p<1.00 were clustered (Supplementary Table 3B). These generated 2 clusters meeting predesignated significance, with terms again including KW-1017∼isopeptide-bond (29 genes, Benjamini p=0.015), and KW-0832∼Ubl conjugation (36 genes, Benjamini p=0.026, Fig. 2Aii). With no apparent difference between the BOEC HHT datasets (Fig. 2B), we concluded that the data were consistent first, with differences in post translational protein processes between the PTC (HHT) and non-PTC (control) datasets, and second, with recent experimental data demonstrating a ubiquitin pathway is used for nascent protein degradation from PTC-containing mRNAs.^48^

**Fig 2.**
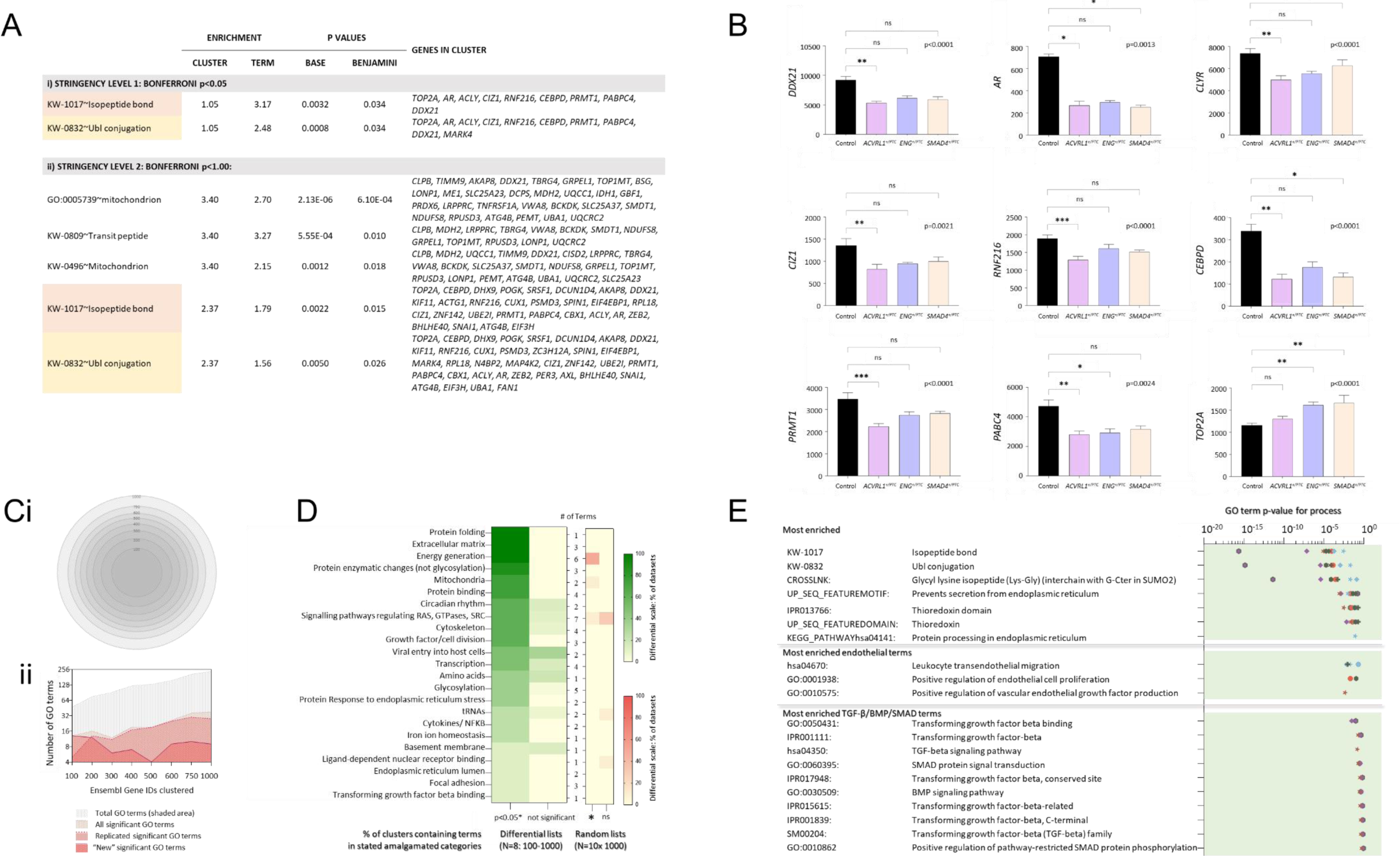
Gene Ontology clustering of genes differentially expressed between HHT and control BOECs. **A)** All significant terms enriched by clustering genes differentially expressed to **i)** Bonferroni p<0.05 (N=30); **ii)** Bonferroni p <1.00 (N=161). **B)** Intergenotype comparisons of expression for the 9 ‘Bonferroni p<0.05’ genes that clustered to both isopeptide-bond and Ubl conjugation. Mean, standard devation and Kruskal Wallis p-values displayed; post-test pairwise p values by Dunn’s test: *p<0.05, **p<0.01, ***p<0.005, ns not significant, p≥0.05. **C)** Broader, lower stringency clustering from the top ranked 100-1,000 genes by HHT differential expression (all in the 5,013 Ensembl gene IDs meeting an intraassay CV10.^41^). **i)** Overview–each of the top-sliced 100-1,000 genes (listed in Supplementary Table 4) are contained within the larger datasets. **ii)** Comparison of gene ontology (GO) term numbers^44^ clustered significantly by the top 100–1,000 genes ranked by differential expression between HHT and control BOECs (details in Supplementary Table 5). The y axis indicates total GO-Direct terms; GO-Direct terms meeting significance; significant terms replicated in ≥1 other gene set, and ‘new’ clusters meeting significance that had not been identified in preceding, smaller datasets. **D)** Heatmaps of amalgamated GO-term processes across the top 100-1,000 ranked HHT differential genes from the 5,013 Ensembl gene IDs meeting CV10 (green), and 10 random datasets of 1,000 genes (red, Supplementary Table 6). Left column: % of datasets containing GO terms meeting significance (p<0.05); right column: proportion of datasets where only a GO term not meeting significance had been clustered. **E)** Specific terms enriched by clustering differentially expressed genes in each dataset. The lowest p-value for each cluster is displayed. Note consistency with the most enriched terms displayed in **C.**

For a broader overview, lower stringency analyses were performed examining HHT-differentially expressed gene normalized to total read counts per library (Supplementary Methods; Fig. 2Ci). As shown in Fig. 2Cii and Supplementary Table 4, in step-wise increases from the top 100 (∼2%) to top 1,000 (∼20%) genes, each increment increased the total number of clusters and GO terms meeting significance, with significant terms emerging repeatedly. There was a marked contrast between the number of significant terms emerging from the clustering of genes differentially expressed in HHT versus control BOECs, compared to 10 sets of randomly selected 1,000 genes from the same BOEC datasets (Fig. 2D). Notably, the most significant terms were not specifically related to endothelial function or BMP/TGF-β signaling predicted by the loss-of-function alleles, but again Uniprot key words KW-1017 (isopeptide-bond) and KW-0832 (Ubiquitin-like conjugation) (Fig. 2E).

### PTC-containing transcript retention in BOECs

The differential expression of genes clustering to protein terms would be most easily explained if HHT BOECs were generating mutant protein from the PTC allele. Initial unsupervised VarScan2^36^ variant call format (vcf) files did not detect the nonsense variants using standard parameters (minor allele frequency >25%). To test whether PTC-containing RNAs were persisting at lower levels in HHT BOECs, supervised examination of RNASeq binary sequence alignment map (bam) files was performed. In 12 of the 16 HHT and control BOECs, alignments to GRCh38/hg38^49^ demonstrated significant mismatches at chr12:51,916,158, chr9:127,829,769 or chr18:51,065,563 corresponding to the HHT donors’ genotypes *ACVRL1* c.1171G>T, p.(Glu391X), *ENG* c.277C>T, p.(Arg93X), and *SMAD4* c.1096C>T, p.(Gln368X) (Fig. 3A, Supplementary Fig. 5). In all cases, HHT nonsense allele alignments were within the expected BOECs, and aligned to spliced exons (almost all alignments sharply defined the exon-intron boundaries, Supplementary Fig. 6). All ratios were skewed from the 1:1 expected for germline heterozygous alleles with a mean wildtype: nonsense allele ratio of 9:1 (range 5:1-13:1, Fig. 3B).

**Fig 3.**
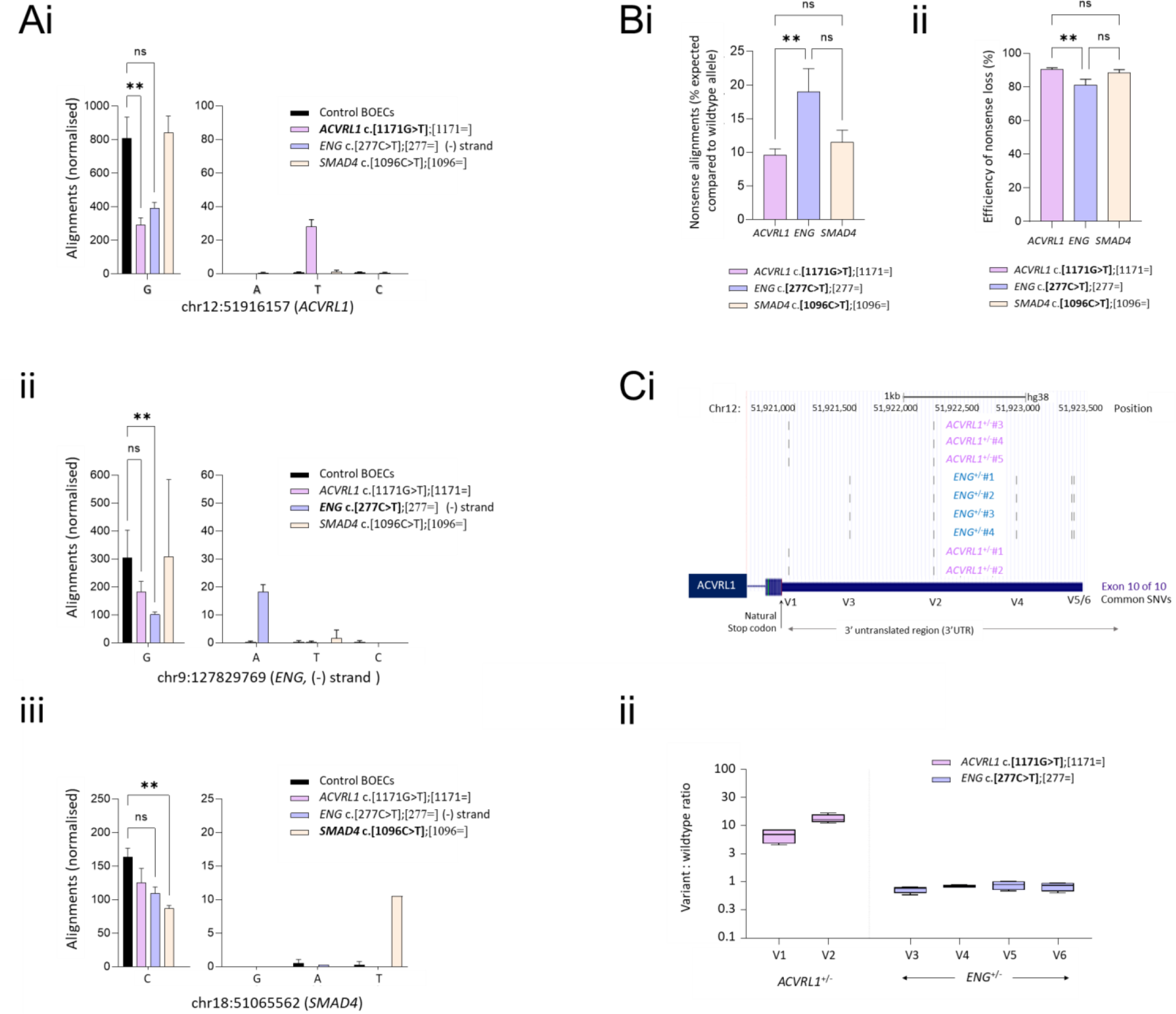
Evidence for loss of nonsense alleles in human blood outgrowth endothelial cells (BOECs). **A)** Quantitative metrics for BOEC RNASeq alignments to nonsense alleles in *ACVRL1*, *ENG* and *SMAD4* genomic loci in GRCh38/hg38^49^ corresponding to known heterozygous pathogenic variants in source BOECs. Left: wildtype allele at **i)** chr12:51,916,158, **ii)** chr9:127,829,769, **iii)** chr18:51065563. Right: alternate alleles at genomic position at 10X scale. Colour-code key describes both alleles for each donor with the relevant nonsense donor allele indicated in bold. Mean and standard deviation displayed. Individual pairwise p-values (** p<0.005) calculated by Dunn’s test post Kruskal Wallis (overall p-values p<0.0001 [*ACVRL1*], p=0.0002 [*ENG*], p=0.003 [*SMAD4*]). **B)** Alignments to PTC allele in the HHT BOECs by genotype, colour coded as in **A)**. **i)** Percentage of total expected if equal to wildtype allele alignment expected for heterozygotes. **ii)** Percentage “loss” of total expected allele alignments. **C)** Alignments to common single nucleotide variants (SNVs) in *ACVRL1* 3’ UTR quantified by VarScan2.^36^ **i)** Genomic (GRCh38/hg38^49^) positions of six common single nucleotide variants (SNVs V1-V6) in BOECs from donors heterozygous for *ACVRL1* c.1171G>T, (p.Glu391X), or *ENG* c.277C>G, (p.Arg93X). Positions are illustrated by custom tracks uploaded to the University of California Santa Cruz (UCSC) Genome Browser.^50,51^ **ii)** Variant: wildtype ratios in *ACVRL1*^+/PTC^ and wildtype *ACVRL1*^+/+^; *ENG*^+/PTC^ BOECs. The higher ratios for V1 and V2 in *ACVRL1*^+/PTC^ BOECs can be attributed to presence *in* cis with wildtype c.1171G, loss of the c.1171T inframe transcripts generating the PTC, and presence of the SNV in all (V1), or only longer (V2) *ACVRL1* 3’UTR transcripts. (See also Supplementary Fig. 7).

Retention of PTC-containing transcripts at levels below those detected by standard variant call file metrics would enable common heterozygous variants *in cis* with the wildtype allele to emerge as over-represented rather than the sole sequence, and this was the case: By chance, both *ENG* ^+/PTC^ and *ACVRL1*^+/PTC^ donors were heterozygous for common variants in the *ACVRL1* 3’untranslated region (UTR) which was examined in the UCSC Genome Browser^50,51^ (Fig 3Ci). VarScan2 indicated the expected 1:1 ratio for four different *ACVRL1* 3’UTR variants in *ENG* ^+/PTC^ BOECs (i.e. *ACVRL1*^+/+^), but skewed allele ratios for the two *ACVRL1* 3’UTR variants in *ACVRL1* ^+/PTC^ BOECs (Fig. 3C). The findings were consistent with persistence of a proportion of *ACVRL1* PTC-containing transcripts to the 3’UTR, which would potentially permit in-frame protein translation.

### Pulse chase examination of ENG protein in HHT and control BOECs

To test aberrant production of mutant protein, we focused on endoglin which had the highest retention of PTC-containing transcripts in BOECs (Fig. 3B), and well-established monoclonal antibodies for pulse chase experiments (mAbs PD31, P4A4 and SN6h^52,53^). These were optimised for use in BOECs as detailed in the Supplementary Methods. Although most HHT-causal *ENG* nonsense variants are too proximal (5’) for predicted proteins to contain both of the ENG orphan (OR) domains that span the mAb epitopes (Fig. 4A), we were able to establish BOECs from an HHT patient heterozygous for a rare nonsense substitution (*ENG* c.1306C>T) generating a PTC at Q436, at the end of the zona pellucida (ZP)-N domain. This predicted that the complete OR1/2 domains and wildtype tertiary structure of all SN6h epitope residues (A119-G230)^53^ would be present in a protein truncated at the PTC (Fig. 4A). Even so, Western blot and pulse chase analyses displayed no size differences in endoglin SN6h-reacting protein extracts from *ENG* Q436X BOECs compared to control BOECs (Fig. 4B).

**Fig. 4:**
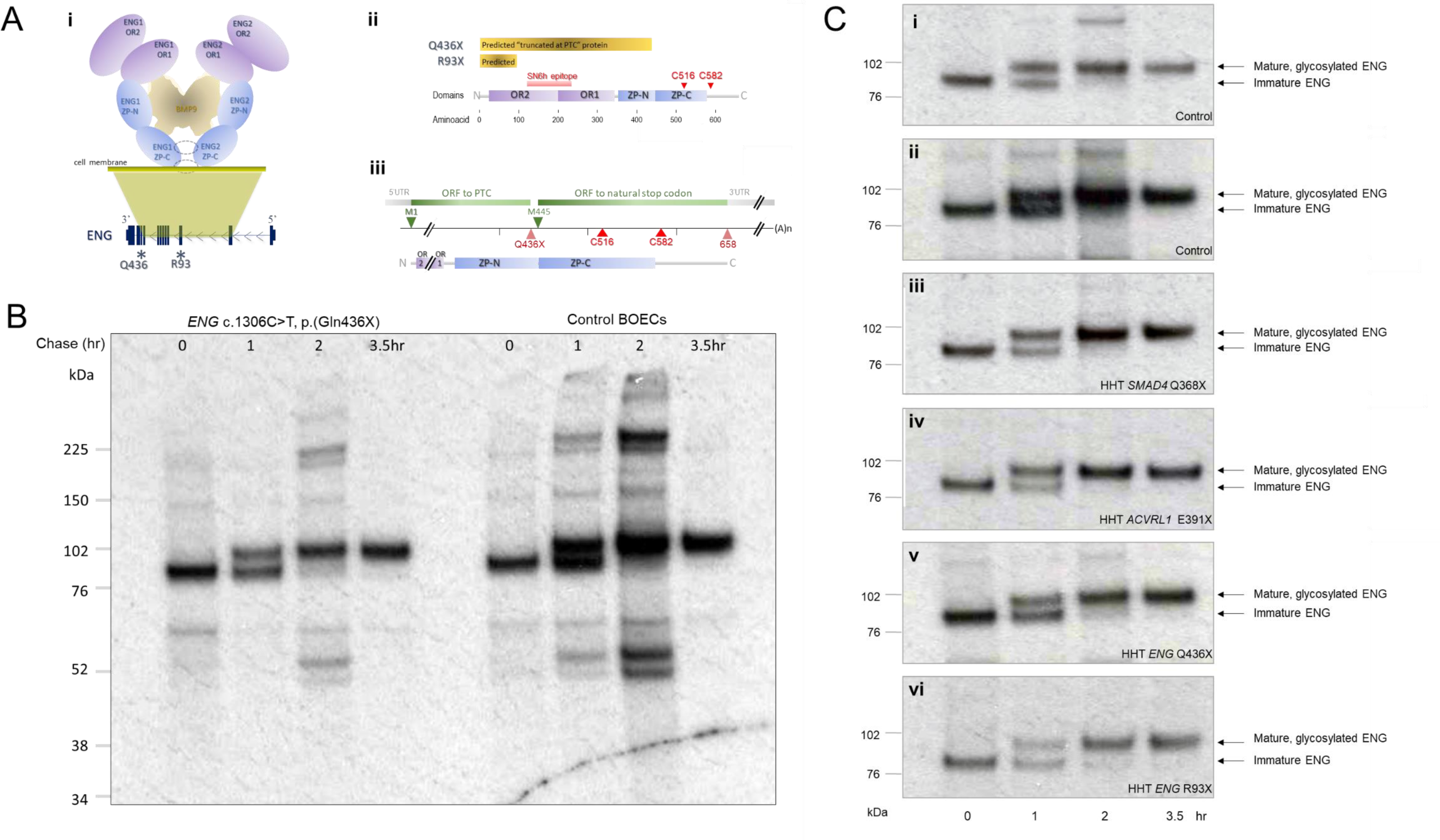
Testing truncated ENG protein expression in normal and HHT endothelial cells. **A)** ENG cartoons: **i)** Extracellular ENG protein domains in BMP-binding heterodimeric conformation by genomic DNA origin, highlighting Q436 distal to orphan domains OR1 and OR2, but proximal to intermolecular dimerization cysteines C516 and C582 that are sited in and beyond the zona pellucida (ZP) domains.^53^ **ii)** “Predicted” truncated ENG Q436X and R93X proteins by wildtype ENG domains and SN6h epitope^52,53^). **iii) ENG** Q436X open reading frames from methionine (M)1 to PTC, and M445 to natural stop codon. **B)** Pulse chase experiments demonstrating generation of endoglin protein in control and *ENG* c.1306C>T, (p.Gln436X) BOEC lysates that were size fractionated on a reducing gel. Note no difference in patterns of sub full length proteins in HHT or control BOECs, and no evidence for a truncated protein product at an expected size based on truncation at the PTC. **C)** Comparison of maturation patterns of full length endoglin protein following pulse chase in BOECs of **i-iv)** ENG^+/+^ (wildtype) ‘controls’ and **v-vi)** *ENG* ^+/PTC^) BOECs: **i)** Control A; **ii)** Control B (same blot as Fig. 5B); **iii)** *SMAD4* c.1096C>T, (Q368X); **iv)** *ACVRL1* c.1171G>T, (E391X), and **v**) Exon 10 *ENG* c.1306C>T, (Q436X, same blot as Fig. 5B); **vi)** Exon 3 *ENG* c.277C>G, (R93X).

The maturation of wildtype ENG protein was examined across further HHT and control BOECs. Commencing the chase when only newly-synthesised, immature endoglin protein was present (Fig. 4C), mature protein became the more pronounced species within 1hr in control BOECs, and was the only species visible by 2hr (Fig. 4Ci, 4Cii). For *SMAD4^+/PTC^* (Fig. 4Ciii), and *ACVRL1^+/PTC^* BOECs (Fig. 4Civ), a similar pattern was observed. However, as seen in Fig. 4B, in BOECs heterozygous for *ENG* c.1306C>T (Q436X) or the more proximal *ENG* c.277C>G (R93X), immature *ENG* protein remained the more pronounced species at 1hr, with traces still present at 2hr (Fig. 4Cv,vi).

### Potential origins of aberrant protein

Two alternate mechanisms generate aberrant proteins that could be indistinguishable by pulse chase from the wildtype protein – PTC read-through (generating full length protein with a missense substitution), and ribosomal re-initiation (generating shorter, amino-truncated proteins that would also escape detection in BOECs using available methods).^5,6^ As detailed in Supplementary Table 7, no PTC had features to suggest strong PTC read-through enhancement, though all PTC contexts were in support of ribosomal re-initiation. Further, since PTC readthrough is a therapeutic strategy^33,54^ and no HHT pathogenic missense substitutions are described at these variants (Supplementary Table 7), if high level natural read-through were the mechanism, the *ENG^+/PTC^* BOECs with highest PTC persistence would be predicted to resemble control BOECs more than other HHT BOECs. Instead, as shown by the global analyses in Fig. 1A, *ENG^+/PTC^*BOECs resembled control BOECs less than other HHT BOECs (Fig. 1A). Thus the BOECS with the least efficient degradation of PTC transcripts were the most distinct (Fig. 5A).

**Fig 5.**
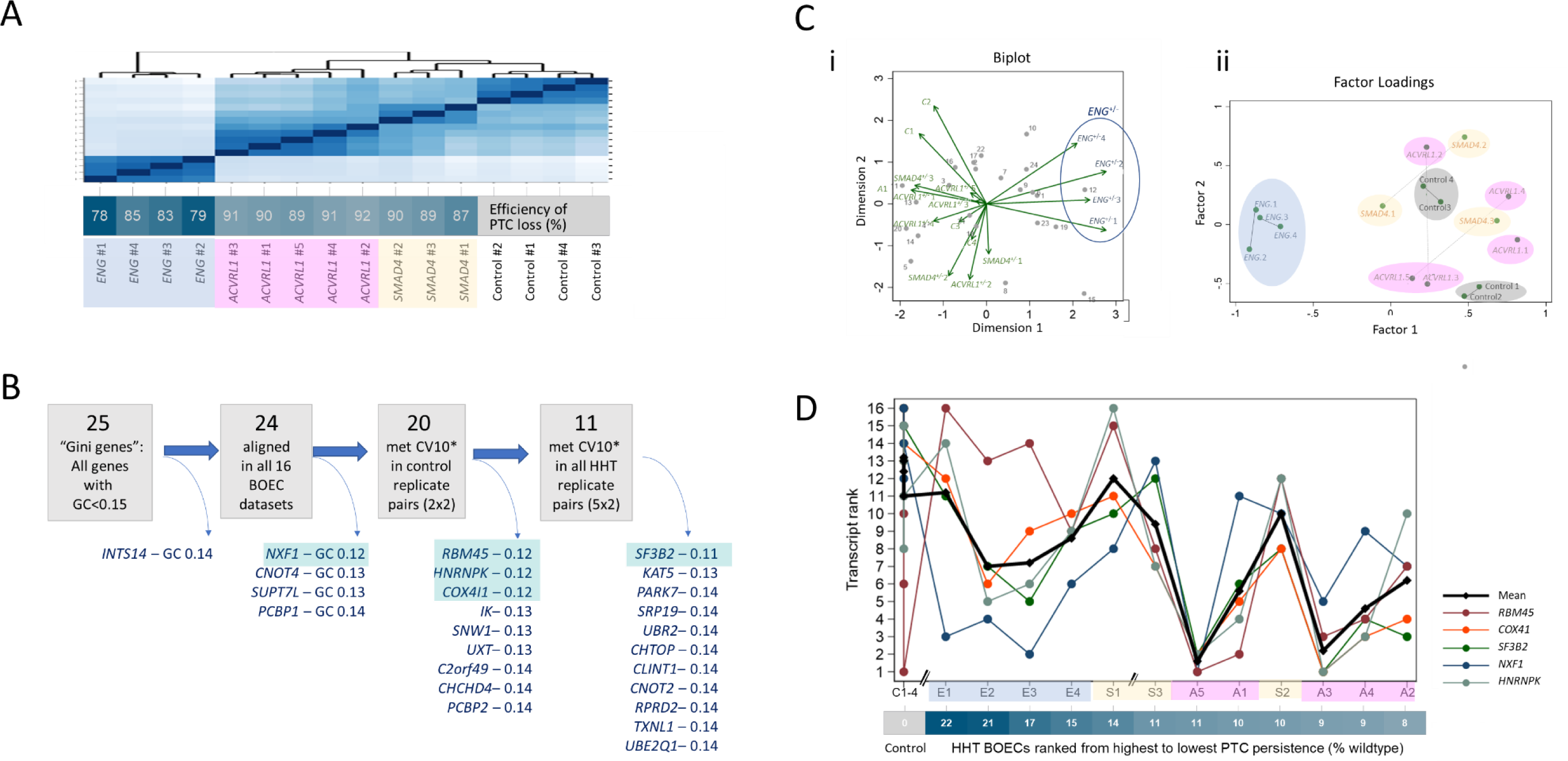
PTC sequence and endothelial contexts. **A)** Unsupervised Euclidean distance analyses for the 16 BOEC RNASeq datasets: Note efficiency of PTC loss was not associated with the differences identified by the initial global data. **B)** BOEC expression of the 25 lowest Gini Coefficient (GC <0.15) genes^39,40^ with GC<0.13 genes highlighted in blue (details in Supplementary Table 2). **C)** Unsupervised analyses of the 24 BOEC-expressed Gini genes ranked in 16 BOEC cultures. **i)** Multivariate analysis of genes (observations, grey dots) across the BOECs (variables, green arrows). Note separation of the 4 *ENG^+/PTC^* BOEC cultures**. ii)** Principal factor analysis on 16 BOEC datasets using ranks of the 24 GINI genes. The 15 retained factors and 120 evaluated parameters again distinguished the *ENG*^+/PTC^ BOEC cultures (blue) from *ACVRL1*^+/PTC^ (pink); *SMAD4*^+/PTC^ (yellow) and control BOECs (grey). **D)** Respective ranks of the 5 lowest GC genes (highlighted blue in **B**) across BOEC cultures ranked by % PTC persistence and genotype (C: control, E: *ENG*^+/PTC^, S: *SMAD4*^+/PTC^, A: *ACVRL1*^+/PTC^. Note switch between 14 and 11% persistence for *NXF1* (blue) and *RBM45* (brown).

### GINI genes also distinguish ENG^+/PTC^ BOECS

Our final analyses were intended simply to explore further evidence for differences between *ENG^+/PTC^* and other HHT BOECs. We have previously shown that for human monocyte rRNA-depleted RNASeq libraries, housekeeper normalisation using the 25 genes with GINI coefficients (GC) <0.15^39,40^ reduced control variability and that the relative ranking of these genes was similar across 24 donor-treatment combinations,^55^ in keeping with data from cell lines indicating these are the least variant of all human transcripts.^39,40^ In contrast, our preliminary studies using the least variant genes in the BOECs had identified apparent discrepancies.^56^

We therefore performed unsupervised hierarchical clustering for all 25 genes with GC<0.15 (Fig. 5B). Unsupervised multivariate analyses of these ‘housekeeper’ genes ranked by relative expression across the 16 BOEC cultures again demarcated the 4 cultures of *ENG^+/-^* BOECs (Fig. 5C). The unsupervised analyses were performed in the opposite direction, and expected to confirm the observations observed in HHT-derived monocytes without PTCs.^55^ Instead, across the HHT BOEC cultures harboring PTCs, there was marked variability in GINI gene respective rankings (Fig. 5D).

### PTC associations with cellular stress phenotypes

Within BOECs, for the least variant (GC 0.11-0.12) genes, the respective rankings of *RBM45* and *NXF1* were seen to switch between the BOECs with 14% and 11% PTC persistence (Fig. 5D). *RBM45* encodes RNA binding protein 45 (RBM45) that binds to m^6^A-modified mRNAs^57^ and forms reversible nuclear inclusions during stress,^58^ while *NXF1* encodes the nuclear RNA export receptor for bulk (homeostatic) export of mRNAs, dissociated from mRNAs under stress.^59,60^ Thus, the RNASeq findings for these previously considered near-invariant genes would be consistent with the highest PTC-expressing BOEC cultures exhibiting a stressed phenotype (Fig. 6).

**Fig 6.**
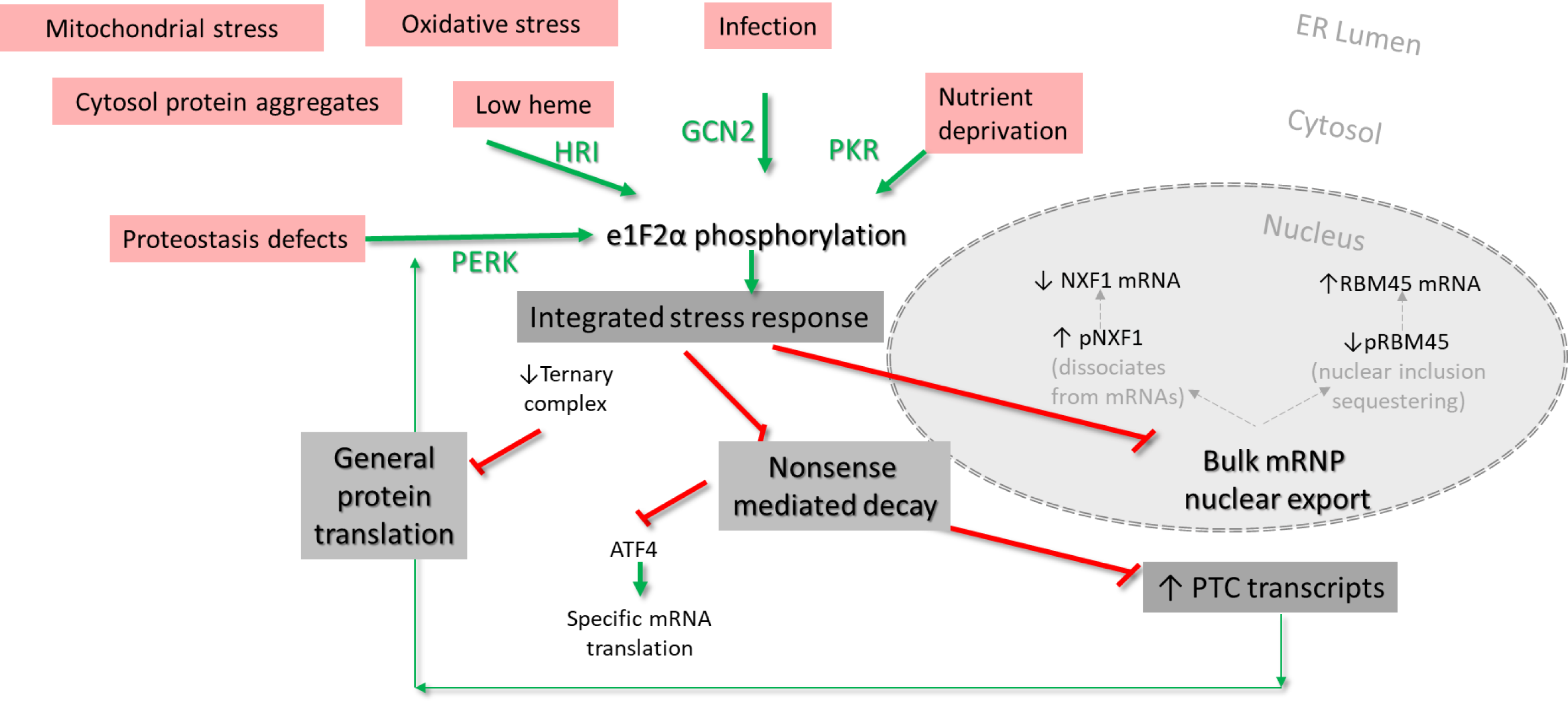
Extended model of stress and premature termination codon interaction. As described,^61-64^ in the integrated stress response (ISR), diverse cytosolic stresses are sensed by four kinases, protein kinase RNA-like ER kinase (PERK), heme-regulated eIF2α (HRI) kinase, general control nonderepressible 2 (GCN2) kinase and protein kinase R (PKR). PERK, HRI, GCN2 and PKR can each phosphorylate the α subunit of eukaryotic initiation factor 2 (eIF2α) which inhibits assembly of the charged methionyl-initiator tRNA eIF2•GTP•methionyl-intiator tRNA ternary complex (TC) required for new protein synthesis. This inhibits translation-dependent nonsense mediating decay (NMD) which allows PTC-harbouring transcripts to persist. Genes regulated by upstream open reading frames use NMD suppression to increase full length protein translation by ribosomal reinitiation. Examples include *ATF4* encoding activating transcription factor 4 that acts as a master transcription factor during the ISR. Stress conditions also modify export factors preventing bulk mRNA export via nuclear RNA export factor 1 protein (pNXF1), and dissociating other RNA binding proteins such as the RNA binding motif protein 45 (pRBM45)^57,58^ that aggregates in nuclear foci. Although GCN2 is classically considered to sense viral infection, a variety of bacterial and other microbial infections are now implicated.^63^ Where the ISR is not transient and stress sustained, apoptosis results.61-64 Note the only new data in the cartoon incorporate low GINI coefficient genes *NXF1*^59,60^ and *RBM45*^57,58^ where theoretical considerations^34,65^ suggest mRNAs are likely to be higher to replace ‘sequestered’ RBM45 protein, and lower where free protein concentrations of NXF1 are increased.

## DISCUSSION

Frameshift and nonsense mutations are common causes of genetic disease, and their PTC-containing transcripts are often thought either to be fully degraded by NMD, or to persist to generate proteins truncated at the PTC. Using patient-derived blood outgrowth endothelial cells, we showed neither assumption to be correct. Specifically, we demonstrated that a fraction of PTC-containing transcripts are retained in spliced RNA transcripts at differing proportions between replicate cultures from the same donor. While genes differentially expressed between patient and control endothelial cells were consistent with pathway-specific changes due to these loss-of-function variants, RNASeq profiles also indicated a second, pathway-independent set of differentially expressed genes related to ubiquitin protein degradation pathways. BOECs with greatest PTC persistence were discriminated in unsupervised hierarchical clustering, and in a state displaying a stress-compatible switch in normally invariant low GINI coefficient genes. In other words, PTC-generating mutations were associated with a second phenotype beyond the conventional pathway-specific perturbations that have been the focus of models for human genetic disease.

Study strengths include the integration of recent developments in RNA biology with molecular medicine, and study design elements that focused on variant examination in endogenous cells from carefully selected donors. Confidence in discovery analyses was enhanced by single cell qRT-PCR validations; restricting analyses to genes with low intra-replicate variability; focusing on findings robust to different analytic methods; and using exacting pulse chase characterization of primary, non-transformed endothelial cells in as close to the natural state as possible. The main weakness is the relatively small number of BOEC datasets examined, although this has been a more rigorous evaluation across all 3 major HHT genotypes in a way that has not been previously performed due to the rarity of the *SMAD4*^+/PTC^ genotype in HHT.^11,66^ Another study limitation is that it does not distinguish processes generically associated with a PTC, from processes that are specific to HHT cells. Although unlikely given the nature of the RNA, protein and pathway findings, we cannot exclude the changes in some way being augmented in HHT BOECs.

Detailed mechanistic dissection needs to be the subject of future work, incorporating the latest experimental data where a reporter system identified a ubiquitin pathway for nascent protein degradation from PTC containing mRNAs.^48^ That said, the current data already support testable models that may solve and facilitate mitigation of previously unexplained phenotypic differences:

- First, while all HHT BOECs displayed differential gene expression profiles supporting pathway-specific changes associated with disease causation^11-28^ and isopeptide bond/ubiquitin-like conjugation relevant to post translational protein degradation, it was *ENG*^+/PTC^ BOECs with the highest PTC burdens that displayed the most distinctive transcriptomes.
- Second, there was marked variability in persistence of PTCs between replicate cultures from the same donor. The cellular state may not have been examined previously as shown by the distinction using normally stably expressed genes, and points to a potential cut off at between 11-14% PTC persistence when two of the most invariant (GC<0.13) housekeeper genes switched expression to a pattern more in-keeping with stressed cells.
- The data point to the *ENG^+/PTC^* variants being more detrimental in BOECs. It is not possible to say whether *ENG^+/PTC^* mutant monomers or heterodimers are more challenging for protein degradation pathways than other HHT mutant proteins (it has long been known that many HHT mutant, non-functional proteins are retained in the endoplasmic reticulum for degradation.^18-23,67^) Homeostatic mechanisms may simply be saturated by excessive production of mutant proteins, irrespective of their nature. Consistent with this, the reversal of *RBM45* and *NXF1* expression patterns was observed with PTC persistence of ≥14% in one of *SMAD4*^+/PTC^ BOEC cultures in addition to all 4 *ENG*^+/PTC^ BOEC cultures.

While it is plausible that varying PTC retention rates of unknown origin were the cause of divergent cellular phenotypes in the BOECs, we favor an alternate, testable interpretation. It is recognized that through the integrated stress response and phosphorylation of eIF2α, a variety of cellular stresses suppress protein translation and NMD and for transient stress, that this suppression is reversible.^61-64^ Thus, stress inhibition of NMD would increase the number of PTC-harboring transcripts, and cell burden^48^ consequences when protein translation resumed.

In summary, we provide evidence that in blood outgrowth endothelial cells (BOECs), operating alongside pathway-specific consequences due to their loss-of-function nature, PTC-generating variants provide a parallel and novel genotypic cause of cellular variability. Precise mechanisms, contributions from environmental stressors, and relevance to other PTCs will need to be the subject of future work.

## Acknowledgments

The project received specific funding from The US Department of Defense Discovery Award W81XWH-16-1-0607 (PI Aldred, CoI Shovlin, *Stopping the Stops: A Novel Therapeutic Approach for Hemorrhage from Vascular Malformations*); The National Institute for Health Research Imperial BRC Institute for Translational Medicine and Therapeutics (PI Shovlin, CoI Aldred; *Nonsense read-through topical therapies for patients with inherited genetic diseases: Proof of concept studies in hereditary haemorrhagic telangiectasia*); and Imperial College Healthcare NHS Trust (PI Shovlin, CoI Noseda; *Towards NHS Laboratory functional assays of genomic variants of uncertain significance –defining shared monocyte/target endothelial signatures in inherited vasculopathies*). MAA was supported by the National Institutes of Health (grant R35HL140019). The Genotype-Tissue Expression (GTEx) Project was supported by the Common Fund of the Office of the Director of the National Institutes of Health, and by NCI, NHGRI, NHLBI, NIDA, NIMH, and NINDS. The data used for the analyses described in this manuscript were from the V8, Aug 2019 release accessed in October and November 2021. We thank the study participants for their willing cooperation in these studies. The views expressed are those of the authors, and not necessarily those of the NHS, the NIHR or the Department of Health and Social Care

## Author contributions

MBH, MAA and CLS devised the BOEC strategy. MBH, DP and MAA performed BOEC cultures. DP prepared RNA, co-designed pulse chase experiments, performed pulse chase experiments, and provided blots for Fig. 4B/C. AB performed early RNASeq analyses, devised the CV10 approach and contributed to sc qRT-PCR design. PCG and MN designed and performed sc qRT-PCR, bulk flow cytometric analyses, and provided data for Supplementary Table 1 and Supplementary Fig. 2A/B. SJM advised on the integrated stress response. CLS reviewed and phenotyped patients, recruited patients, contributed to sc qRT-PCR and pulse chase experimental design, performed data analyses generating all other Figures/Tables, and wrote the manuscript. All authors have reviewed and approved the final manuscript.

## Data availability

Non-sensitive data underlying this article are available at <u>10.5281/zenodo.5201823 and</u> can be used under the Creative Commons Attribution licence. Primary sequence data used in this research was collected subject to the participants’ informed consent. Access to these data will only be granted in line with that consent, subject to approval by the project ethics board and under a Data Sharing Agreement. Blood outgrowth endothelial cells used in this research were collected subject to the informed consent of the participants. Access will only be granted in line with that consent, subject to approval by the project

