## Supplementary material for "Heterozygous transcriptional signatures unmask variable premature termination codon (PTC) burden alongside pathway-specific adaptations in blood outgrowth endothelial cells from patients with nonsense DNA variants causing hereditary hemorrhagic telangiectasia": Data Supplement

#### SUPPLEMENTARY DATA

**SUPPLEMENTARY METHODS:**
Endothelial cell culture:
Single cell qRT-PCR
RNA Sequencing
RNA sequencing and alignments
RNA sequencing analyses
Intra replicate variability
Defining housekeeper genes for BOECs
Gene Ontology analyses
Metabolic labelling, pulse-chase analysis
**SUPPLEMENTARY TABLES**
Supplementary Table 1: Single cell (sc) qRT-PCR genes and assays
Supplementary Table 2: Low GINI-coefficient genes
Supplementary Table 3: Bonferroni-differentially expressed CV10 genes (DeSeq2 normalized)
Supplementary Table 4: Top 1000 differentially expressed genes (normalized to total counts per library)
Supplementary Table 5: Gene Ontology terms enriched by clustering top 100-1,000 Gene IDs meeting CV10
Supplementary Table 6: Gene Ontology terms enriched clustering 10 sets of 1000 random genes
Supplementary Table 7: PTC sequence contexts
**SUPPLEMENTARY FIGURES**
Supplementary Fig. 1: Blood outgrowth endothelial cell (BOEC) derivation indices
Supplementary Fig. 2: Flow cytometry and RNASeq validations of blood outgrowth endothelial cells
Supplementary Fig. 3: Quality scores generated across the 150bp reads for the 4 pairs of untreated BOECs
Supplementary Fig. 4: Low Gini-coefficient reference genes for RNASeq normalizations
Supplementary Fig. 5: Examples of raw sequence reads for nonsense variant detection
Supplementary Fig. 6: Alignments of reads to spliced transcripts
Supplementary Fig. 7: Detailed examination of common single nucleotide variants in *ACVRL1* 3' UTR
**SUPPLEMENTARY REFERENCES:**
**APPENDIX:**
Supplementary Table 4: Full list of 1000 differentially expressed genes (normalized total counts per library)

### SUPPLEMENTARY METHODS

#### Endothelial cell culture:

This research was approved by the East of Scotland Research Ethics Service (EoSRES, 16/ES/0095). All participants provided written informed consent. Blood outgrowth endothelial cells (BOECs) were established from healthy volunteers, and HHT patients heterozygous for pathogenic DNA variants in *ACVRL1*, *ENG*, or *SMAD4* as described.[1] First BOEC colonies were identified 8-20 days after plating peripheral blood mononuclear cells (PBMCs). Once colonies emerged, 22/25 proliferated steadily with the median time to third passage of 16 (range 6-28) days. This rate approximated to a normal distribution (Supplementary Fig. 1). For the current project, BOECs were established from volunteers and 24 HHT patients heterozygous for 10 different nonsense ("stop gain") pathogenic DNA variants in HHT genotypes: *ENG* c.277C>T, (p.Arg93X); *ENG* c.744C>A, (p.Tyr258X), *ENG* c.1306C>T, (p.Gln436X), *ACVRL1* c.172G>T, (p.Glu58X), *ACVRL1* c.247G>T, (p.Glu83X), *ACVRL1* c.475G>T, (p.Glu159X), *ACVRL1* c.1171G>T, (p.Glu391X); and *SMAD4* c.1096C>T, (p.Gln366X). Each nonsense variant was established from one to five different donors (median 2). Current studies *ACVRL1* c.[1171G>T];[c.1171=]; *ENG* c.[277C>T];[277=]; *ENG* c.[1306C>T]; [1306=], and *SMAD4* c.[1096C>T];[1096=].

#### Single cell qRT-PCR

Quantitative (q)RT-PCR was performed in single HHT and control BOECs. $SMAD4^{+/PTC}$ was selected as the primary test HHT genotype because the SMAD4 protein is downstream of *ACVRL1* and *ENG* gene products in the canonical signalling pathways perturbed in HHT.[2,3] BOECs were cultured in the presence and absence of BMP9 at 10ng/ml for 1 hr. Viable (DRAQ7 negative) BOECs were single-cell sorted prior to qRT-PCR quantifications. For each group, 44 single-cells were sorted directly into 96-well-plates containing pre-amplification mix, and specific target amplification was performed using CellDirect One-Step qRT-PCR Kit (Invitrogen, Thermo Fisher, Waltham, MA) as previously described.[4,5] 48 genes were selected as either likely relevant to the HHT cellular phenotype based on literature reviews; endothelial gene markers as validation for endothelial differentiation; or important negative controls. qRT-PCR was performed using 48.48 Dynamic Array chips in BioMark HD system (Fluidigm, San Francisco, CA) and TaqMan primer/probe sets (ABI, Thermo Fisher, Waltham, MA) as listed in Supplementary Table 1. Ct values were obtained using Fluidigm Real-Time PCR Analysis software (v4, Fluidigm, San Francisco, CA). Samples were normalized to *UBC* expression. All data was analysed and plotted using an R script developed in house. Flow cytometry and single cell sorting were performed using a FACS Aria II (Becton Dickinson, New Jersey) equipped with 355 nm ultraviolet, 405 nm violet, 488 nm blue, 561 nm yellow–green and 640 nm red lasers and analysed using FlowJo™ v10.[6] Directly conjugated, monoclonal antibodies from Biolegend (San Diego, CA) were used. Genes were also ranked as the product of number of expressing single BOECs and relative expression, in order to perform two-way comparisons with RNASeq alignments. Here, the 48 genes examined by sc qRT-PCR were categorized by tertiles, or ranked individually for linear regression using the normally distributed log of the mean untreated RNASeq alignments as the dependent variable STATA IC v 15.0 (Statacorp, College Station, TX).

#### RNA Sequencing

##### RNA sequencing and alignments

RNA was extracted from 16 separate cultures of confluent BOECs from 4 donors, cultured in the presence and absence of BMP9 at 10ng/ml for 1 hr, using methods described previously.[7,8] For RNA sequencing, following ribosomal (r)-RNA depletion, RNA was fragmented and random primed for first and second strand cDNA synthesis, end repair, 5' phosphorylation, adaptor ligation, PCR enrichment and Illumina HiSeq sequencing using paired-end 150bp reads (Genewiz, Leipzig, Germany). Sequenced reads were trimmed and aligned to *Homo sapiens* GRCh38[9] using STAR aligner v2.5.2b, and counting of unique gene reads that fell within exon regions using Subread package v1.5.2 (Genewiz, Leipzig, Germany). Alignments were normalized to the total alignments per library. In blinded analyses, Genewiz analytics, generated variant call format (vcf) files using VarScan2.[10]

Binary sequence alignment (bam) and vcf files were analysed in Galaxy Version 2.4.1[11] and uploaded to the UCSC Genome Browser,[12,13] and Integrated Genome Browser (IGB) 9.1.8[14] for visualisation. Percentage loss of the nonsense allele in $ACVRL1^{+/-}$, $ENG^{+/-}$ and $SMAD4^{+/-}$ BOECs was calculated assuming that without nonsense mediated decay (NMD),[15-18] in these heterozygous cells there would have been an approximately equal number of alignments to wildtype and nonsense alleles. For individual BOEC cultures/RNASeq libraries, the % persistence of the nonsense pathogenic variant was compared to ratios of common single nucleotide variant (SNV) alignments that VarScan2 identified in the untranslated regions of *ACVRL1*, *ENG* and *SMAD4*, and to expression of low GINI coefficient housekeeper genes as described below.

#### RNA sequencing analyses

##### Intra replicate variability

Experimental design had included 7 replicate pairs, and across all replicate pairs the median ratio of normalized read counts for 16,807 genes between replicates was 0.98 (interquartile range 0.95-1.08). Early analyses attempting to use all 16,807 genes are described elsewhere.[19] It was recognised that a proportion of genes demonstrated marked variability between HHT replicate cultures, with no apparent pattern. Therefore, we adopted measures used for immunoassays: only genes with an intra-assay coefficient of variation <10% (meeting CV10)[20] in all untreated replicates were analysed for the current manuscript: CV was calculated for each gene in each replicate pair as:

$$CV = 100x\ standard\ deviation\ (SD)/mean)$$

'CV10 genes' were those where all untreated replicates met CV10.

##### Differential gene expression

DeSeq2 normalizations[21,22] were performed initially aiming to use the 25 genes with GINI coefficients (GC) <0.15 within multiple cell lines,[23,24] i.e. the least variable of all known human transcripts, as for primary human monocytes.[25] These are presented in Supplementary Fig. 4 and the main manuscript. The most stringent discovery analyses focussed on the subgroup of 5,013 genes that met CV10[20] ranked by DeSeq2[21,22] normalized expression using the 8 least variable of the GC<0.15 GINI housekeeper genes in BOECs (*HNRNPK, COX4I1, IK, UBR2, UBE2Q1, SNW1, TXNL1*, and *KAT5*, as shown in Supplementary Fig. 4). Briefly, two tailed, two sample unequal variance t tests were performed comparing DeSeq2 normalized untreated control BOECs and untreated HHT (*ACVRL, ENG, SMAD4*) BOECs, then multiplied by 5,013 for Bonferroni-adjusted p values. Differential expression analyses were performed on genes where DeSeq2 normalized expression differed between untreated HHT and control BOECs to Bonferroni p<0.05 (30 genes, Supplementary Table 2A), and all meeting Bonferroni p<1.00 (159 genes, Supplementary Table 2B). For a complementary approach, read counts normalized to total library counts[21,22] (as supplied by Genewiz), were ranked according to the ratio of counts in all untreated HHT (*ACVRL, ENG, SMAD4*) BOECs over untreated control BOECs counts. Genes with greater intra-replicate variability were weighted down by including a confidence index[26] from the standard deviation of untreated control BOECs. To enable unidirectional ranking, the square root of the square of the calculation below was used to rank:

$$Ln\left(\frac{Mean\ ENG,\ ACVRL\ and\ SMAD4\ untreated\ BOEC\ alignment}{Mean\ untreated\ control\ BOEC\ alignment}\right)*\left(-\ln\left(\frac{1}{Standard\ deviation}\right)\right.$$

##### Gene Ontology analyses

Gene Ontology (GO) enrichment analysis was performed by Functional Annotation Clustering using the Database for Annotation, Visualization and Integrated Discovery (DAVID) v6.8.[27-29] For the stringent Bonferroni-based lists, all terms were included. When cluster analyses were performed for the top 100, 200, 300, 400, 500, 600, 750, and 1,000 genes differentially expressed by total library counts (Supplementary Table 3), clusters were restricted to GO Direct categories directly annotated by the source database for molecular function [MF], cellular component [CC] and biological processes [BP] (Supplementary Table 4). As previously,[38,30] we also examined random sets of genes derived from the BOEC gene dataset (Supplementary Table 5). Clusters meeting enrichment of ≥1.2 fold and GO term p<0.05 were compared between experimentally-ranked and randomly-generated datasets. Data were visualised using GraphPad Prism 9 (GraphPad Software, San Diego, CA), and STATA IC v 15.0 (Statacorp, College Station, TX).

#### Metabolic labelling, and pulse-chase analysis

For metabolic labelling, pulse and chase periods were tested in a series of modifications to the method of Pece et al.[30,31] Briefly, HUVEC and BOECs were grown in parallel to 90% confluence in 6 well plates and incubated at 37°C in methionine-free Dulbecco's modified Eagle's medium (DMEM, low glucose) supplemented with 50µCi/ml of $^{35}S$ Methionine, 10% fetal calf serum (FCS), and penicillin/streptomycin for test "pulses" of 20, 40 and 60 minutes, before washes and incubation in DMEM containing 200µM methionine. Washed BOECs were solubilised in 1% Triton lysis buffer and frozen overnight. SN6h was complexed for 1 hour with Dynabeads

Sepharose A beads (Invitrogen, Thermo Fisher, Waltham, MA), which were washed with a protease inhibitor-free lysis buffer, RIPA buffer (0.05M Tris-HCL pH 7.4, 0.01M NaCl, 1mM EDTA, 0.1% SDS, 0.5% Triton X-100, 1% deoxycholate and TNTE buffer (0.05 M Tris-HCl, pH 7.4, 0.01M NaCl, 0.1% Triton X-100, 1mM EDTA). Following protein estimation and pre-clearance, saturating quantities of the antigen-containing lysate was added and incubated for 1 hr at 4$^{o}$C on a rotisserie. Samples were eluted by heating at 98$^{o}$C for 3 mins in non-reducing gel samples buffer (60mM Tris-buffer, pH 6.8, 2% SDS, 10% glycerol, 0.05% bromophenol blue) and reduced with β2 mercaptoethanol. Immune complexes fractionated by SDS-PAGE on 4-12% gradient gels indicated a 60 min $^{35}$S-methionine "pulse" was optimal to maximise protein intensity (*Fig. SA1*) before maturation of the ENG protein by glycosylation that became evident during a subsequent "chase".

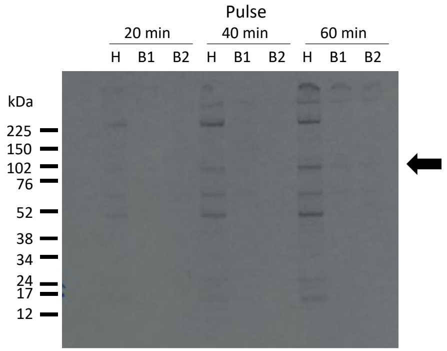

*Fig. SA1:* Optimization of $^{35}$S-methionine pulse period in HUVEC with BOEC comparators. Post fractionation of immune complexes by SDS-PAGE on 4-12% gradient gels. Data are from a 4 day exposure, without signal amplification. Note higher ENG protein expression (monomer arrowed) in HUVEC (H) compared to BOECs (B1 [control] and B2 [*ENG* c.1306C>T, p.Gln436X]). HHT donor]).

The lower expression of ENG in the control BOECs was initially surprising, but subsequently supported by greater similarity in control BOEC alignments to *ENG*, *ACVRL1* and *SMAD4* (main Fig. 2C) than data from normal arteries and non-BOEC endothelial cells in the Genotype-Tissue Expression (GTEx) Project.[32] Data were extracted via the GTEx Portal on 19/04/2022 and are graphed in *Fig. SA2*.

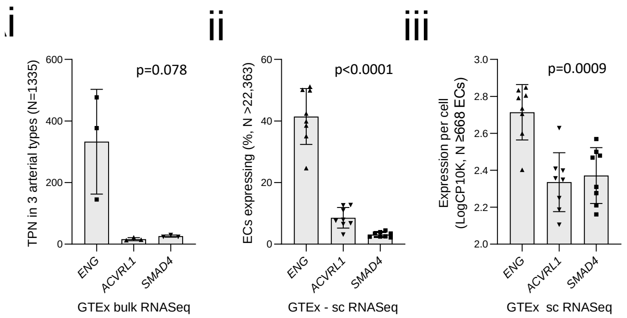

*Fig. SA2:* Expression of *ACVRL1*, *ENG* and *SMAD4* by bulk and single cell RNASeq alignments in the Genotype Tissue Expression Project (GTEx).[32] **(i)** Median transcript alignments per million base pairs (TPM) in 432 aortas, 240 coronary arteries, and 663 tibial arteries. **(ii)** Percentage of single ECs where transcript alignments detected (762 ECs from breast tissue, ≥3111 ECs from esophageal mucosa, 3244 ECs from esophageal muscularis, 7070 left ventricle ECs, 2412 lung ECs, 1124 ECs from skeletal muscle, 2575 prostatic ECs, and 730 ECs from sun-exposed skin on the lower leg). **(iii)** Logarithmic scale expression (by CP10K) in the 688 ECs where transcript alignments were detected as described in **ii**. Note that a CP10K of 2.5 means that 2.5 out of 100,000 UMI counts belong to that cell, equivalent to a fraction of <0.001 (<0.1%). For all graphs, error bars indicate mean and standard deviation, and p values were calculated by Friedman.

For experimental BOEC pulse chase evaluation, analyses were restricted to BOECs from HHT patients with nonsense variants in *ENG* exons, and as $ENG^{+/+}$ controls, control, $ACVRL1^{+/PTC}$ and $SMAD4^{+/PTC}$ donors. Equivalent BOEC numbers were incubated at 37$^{o}$C in methionine-free, low glucose Dulbecco modified eagle medium (DMEM), supplemented with 50µCi/ml of $^{35}$S methionine, 10% fetal calf serum and penicillin/streptomycin for 1 hr ("pulse"), before washes and incubation in DMEM containing 200µM methionine for a further 0, 1, 2 or 3.5hr "chase". At the designated timepoint, washed BOECs were solubilised in 1% Triton lysis buffer and frozen overnight before immunoprecipitation.

#### Supplementary Table 1: Single cell (sc) qRT-PCR genes and assays

| Gene | assay ID |
|---|---|
| *Kdr* | Hs00911700_m1 |
| *Angpt1* | Hs00375822_m1 |
| *Angpt4* | Hs00907078_m1 |
| *Angptl7* | Hs00221727_m1 |
| *TGB1 (CD29)* | Hs00559595_m1 |
| *Pi16* | Hs00542137_m1 |
| *Cnn1* | Hs00154543_m1 |
| *Decorin* | Hs00754870_s1 |
| *Nkx2.5* | Hs00231763_m1 |
| *Gata6* | Hs00232018_m1 |
| *Ereg* | Hs00914313_m1 |
| *Cdh5* | Hs00901463_m1 |
| *THY1 (CD90)* | Hs00174816_m1 |
| *Wt1* | Hs01103751_m1 |
| *Cxcl5* | Hs01099660_g1 |
| *Pecam1* | Hs00169777_m1 |
| *Wnt5a* | Hs00998537_m1 |
| *Twist* | Hs02379973_s1 |
| *Klf4* | Hs00358836_m1 |
| *Pdgfrbeta* | Hs01019589_m1 |
| *Csf1* | Hs00174164_m1 |
| *Cd146 (MCAM)* | Hs00174838_m1 |
| *NT5E (CD73)* | Hs00159686_m1 |
| *Tbx20* | Hs00396596_m1 |
| *TMSB4X* | Hs03407480_gH |
| *Pouf51* | Hs00999632_g1 |
| *Vimentin* | Hs00958111_m1 |
| *Snai1* | Hs00195591_m1 |
| *Snai2* | Hs00161904_m1 |
| *Kit* | Hs00174029_m1 |
| *Nanog* | Hs04260366_g1 |
| *Ptch2* | Hs00184804_m1 |
| *Ccl2* | Hs00234140_m1 |
| *Ang* | Hs04195574_sH |
| *Angptl2* | Hs00765775_m1 |
| *Angptl4* | Hs01101127_m1 |
| *PDGFra* | Hs00998018_m1 |
| *CD44* | Hs01075861_m1 |
| *Coro1b* | Hs01127579_m1 |
| *VWF* | Hs01109446_m1 |
| *Bmper* | Hs01564922_m1 |
| *CD105* | Hs00923996_m1 |
| *Nestin* | Hs04187831_g1 |
| *Baf60b (SMARCD2)* | Hs01015702_m1 |
| *alpha-sma (ACTA2)* | Hs00426835_g1 |
| *Baf60c (SMARCD3)* | Hs00162003_m1 |
| *UBC* | Hs01871556_s1 |

#### Supplementary Table 2: Low GINI-coefficient genes

| Gene | *GC | Encodes | Function | Source | max CV (%) control pairs | max CV (%) in HHT pairs |
|---|---|---|---|---|---|---|
| *SF3B2* | 0.11 | Splicing Factor 3b Subunit 2 | Splicing: component of U2 small nuclear (sn)RNP (subunit 2 of the splicing factor 3b protein complex) | HPA | 3.9 | 8.1 |
| *COX4I1* | 0.12 | Cytochrome C Oxidase Subunit 4I1 | Mitochondria: terminal enzyme of the mitochondrial respiratory chain | GTEx | 3.1 | 13.7 |
| *HNRNPK* | 0.12 | Heterogeneous Nuclear Ribonucleoprotein K | RNA binding protein, complexes with heterogeneous nuclear (hn)RNA | KLIJn | 3.7 | 10.9 |
| *NXF1* | 0.12 | Nuclear RNA Export Factor 1 | Nuclear polyadenylated RNA export factor | HPA | 10.3 | 10.7 |
| *RBM45* | 0.12 | RNA Binding Motif Protein 45 | RNA binding- preferential binds to poly(C) RNA (N6-methyladenosine ($m^6A$)) | HPA | 3.5 | 13.7 |
| *CNOT4* | 0.13 | CCR4-NOT Transcription Complex Subunit 4 | E3 Ubiquitin-Protein Ligase CNOT4- role in deadenylation-dependent mRNA decay. | HPA | 12.2 | 13.7 |
| *IK* | 0.13 | IK cytokine | Nucleus pre-mRNA splicing as a component of the spliceosome | HPA | 1.4 | 10.6 |
| *KAT5* | 0.13 | Lysine Acetyltransferase 5 | Chromatin remodelling | HPA | 3.8 | 8.7 |
| *SNW1* | 0.13 | SNW Domain Containing 1 | Gene transcription regulation by interacting with poly(A)-binding protein 2 | HPA | 1.9 | 12.0 |
| *SUPT7L* | 0.13 | SPT7 Like, STAGA Complex Subunit Gamma | Chromatin-modifying multiprotein complex | HPA | 11.2 | 7.9 |
| *UXT* | 0.13 | Ubiquitously Expressed Prefoldin Like Chaperone | Gene transcription regulation | HPA | 7.0 | 23.5 |
| *C2orf49* | 0.14 | C2orf49 | Component of tRNA-splicing ligase | HPA | 6.8 | 14.3 |
| *CHCHD4* | 0.14 | Coiled-Coil-Helix-Coiled-Coil-Helix Domain Containing 4 | Component of a redox-sensitive mitochondrial intermembrane space import machinery | GTEx | 4.2 | 22.4 |
| *CHTOP* | 0.14 | Chromatin Target Of PRMT1 | Nuclear mRNP formation | HPA | 3.1 | 5.2 |
| *CLINT1* | 0.14 | Clathrin Interactor 1 | Protein transport- clathrin-coated vesicles from the trans-Golgi network | HPA | 3.8 | 6.8 |
| *CNOT2* | 0.14 | CCR4-NOT Transcription Complex Subunit 2 | mRNA synthesis, degradation, transport | HPA | 2.8 | 6.6 |
| *INTS14* | 0.14 | Integrator Complex Subunit 14, also known as Von Willebrand Factor A Domain-Containing Protein 9 | Complex involved in snRNA U1 and U2 transcription and processing. | HPA | no alignments | no alignments |
| *PARK7* | 0.14 | Parkinsonism Associated Deglycase | Peptidase C56, redox-sensitive chaperone, protects neurons against oxidative stress/cell death | HPA | 0.4 | 9.8 |
| *PCBP1* | 0.14 | Poly(RC) Binding Protein 1 | Poly(RC) Binding Protein 1, processing of capped intron-containing pre mRNAs | HPA | 17.2 | 17.0 |
| *PCBP2* | 0.14 | Poly(RC) Binding Protein 2 | Poly(RC) Binding Protein 2, processing of capped intron-containing pre mRNAs | HPA | 7.2 | 13.8 |
| *RPRD2* | 0.14 | Regulation Of Nuclear Pre-MRNA Domain Containing 2 | Part of RNA polymerase II holoenzyme, involved in mRNA 3'-end processing. | HPA | 3.5 | 7.5 |
| *SRP19* | 0.14 | Signal Recognition Particle 19 | SRP-dependent cotranslational protein targeting to endoplasmic reticulum membrane | HPA | 7.6 | 9.8 |
| *TXNL1* | 0.14 | Thioredoxin Like 1 | Enables disulfide oxidoreductase activity. Located in cytosol. | HPA | 1.9 | 9.7 |
| *UBE2Q1* | 0.14 | Ubiquitin Conjugating Enzyme E2 Q1 | Catalyzes the covalent attachment of ubiquitin to other proteins | HPA | 4.7 | 5.7 |
| *UBR2* | 0.14 | Ubiquitin Protein Ligase E3 Component N-Recognin 2 | An E3 ubiquitin ligase component | HPA | 6.0 | 8.2 |

The lowest Gini Coefficient (GC)[23,24] proposed 'housekeeper' genes by expression and coefficient of variation (CV) in control and HHT BOECs. Genes are ordered by GC, and columns detail gene name, function, GC source, and CV categories where CV10 is the intraassay coefficient of variation,[20] and CV10 denotes a CV <10% between replicate pairs. There were alignments to 24 of the 25 GC<0.15 genes in the BOECs (no alignments were detected to *INTS14*). Twenty of the 24 (83%) genes met CV10 in both control replicate pairs (untreated, and BMP9 treated). However, only 11 (46%) also met CV10 in all HHT replicate pairs. In other words, unlike in human monocytes derived from HHT patients with non PTC disease causal variants,[25] the majority of GC genes did not pass the criteria set for examining HHT/control differential expression. Details of products and functions of lowest 25 GINI coefficient genes,[23,24] were sourced through Uniprot[33] and GeneCards[34] databases. RNP: ribonucleoprotein. GC: GINI coefficient. HPA: RNA-seq-based dataset from the Human Protein Atlas group-data sets from 19,628 protein coding genes in 56 cell lines (HPA_C) and 19,613 protein coding genes in 59 tissues (HPA_T).[23,24] GTEx: RNASeq data of 46,711 genes in 53 human tissue samples from the Genotype-Tissue Expression (GTEx) project.[23,24,32] KLIJn: RNA-seq-based analysis of 57,711 genes in 622 human cancer cell lines from EBI Expression Atlas E-MTAB-2706). [23,24]

#### Supplementary Table 3: Bonferroni-differentially expressed CV10 genes (DeSeq2 normalized)

| Ensembl Gene ID | Official gene symbol | P value t-test 2 tailed, 2 sample unequal variance | Bonferroni p-value | Ratio (mean HHT/mean Control) |
|---|---|---|---|---|
| **A) Bonferroni p<0.05** | | | | |
| ENSG00000169083 | *AR* | 2.17428E-07 | 0.001 | 1.359 |
| ENSG00000148337 | *CIZ1* | 2.80332E-07 | 0.001 | 1.612 |
| ENSG00000146701 | *MDH2* | 4.61435E-07 | 0.002 | 1.973 |
| ENSG00000221869 | *CEBPD* | 4.65552E-07 | 0.002 | 0.789 |
| ENSG00000007047 | *MARK4* | 4.93737E-07 | 0.002 | 0.504 |
| ENSG00000087269 | *NOP14* | 5.8029E-07 | 0.003 | 1.354 |
| ENSG00000122862 | *SRGN* | 6.85128E-07 | 0.003 | 0.831 |
| ENSG00000133110 | *POSTN* | 7.7799E-07 | 0.004 | 0.818 |
| ENSG00000067182 | *TNFRSF1A* | 1.0464E-06 | 0.005 | 1.154 |
| ENSG00000232956 | *SNHG15* | 1.26773E-06 | 0.006 | 0.978 |
| ENSG00000183172 | *SMDT1* | 1.34219E-06 | 0.007 | 1.623 |
| ENSG00000134684 | *YARS* | 1.73883E-06 | 0.009 | 1.648 |
| ENSG00000167173 | *C15orf39* | 2.64171E-06 | 0.013 | 0.760 |
| ENSG00000110455 | *ACCS* | 2.81363E-06 | 0.014 | 1.015 |
| ENSG00000089159 | *PXN* | 3.31396E-06 | 0.017 | 0.908 |
| ENSG00000162618 | *ADGRL4* | 3.46757E-06 | 0.017 | 0.964 |
| ENSG00000131473 | *ACLY* | 3.61675E-06 | 0.018 | 1.004 |
| ENSG00000081760 | *AACS* | 3.64992E-06 | 0.018 | 1.229 |
| ENSG00000164124 | *TMEM144* | 3.86772E-06 | 0.019 | 0.803 |
| ENSG00000126457 | *PRMT1* | 4.1425E-06 | 0.021 | 0.704 |
| ENSG00000166794 | *PPIB* | 5.57233E-06 | 0.028 | 1.095 |
| ENSG00000165732 | *DDX21* | 5.65458E-06 | 0.028 | 0.823 |
| ENSG00000090621 | *PABPC4* | 7.2285E-06 | 0.036 | 25.705 |
| ENSG00000131747 | *TOP2A* | 7.31375E-06 | 0.037 | 0.867 |
| ENSG00000011275 | *RNF216* | 8.27766E-06 | 0.041 | 2.148 |
| ENSG00000109107 | *ALDOC* | 8.34378E-06 | 0.042 | 1.271 |
| ENSG00000135069 | *PSAT1* | 8.57446E-06 | 0.043 | 1.046 |
| ENSG00000212719 | *C17orf51* | 9.71802E-06 | 0.049 | 1.241 |
| ENSG00000167487 | *KLHL26* | 9.7862E-06 | 0.049 | 0.688 |
| **B) Bonferroni p<1.00** | | | | |
| ENSG00000162244 | *RPL29* | 9.9457E-06 | 0.050 | 0.884 |
| ENSG00000072736 | *NFATC3* | 1.05045E-05 | 0.053 | 0.889 |
| ENSG00000135763 | *URB2* | 1.11592E-05 | 0.056 | 0.908 |
| ENSG00000155660 | *PDIA4* | 1.11645E-05 | 0.056 | 1.002 |
| ENSG00000089327 | *FXYD5* | 1.12214E-05 | 0.056 | 1.216 |
| ENSG00000176946 | *THAP4* | 1.19594E-05 | 0.060 | 0.938 |
| ENSG00000147065 | *MSN* | 1.23077E-05 | 0.062 | 1.290 |
| ENSG00000150551 | *LYPD1* | 1.28748E-05 | 0.065 | 1.020 |
| ENSG00000196365 | *LONP1* | 1.42373E-05 | 0.071 | 1.182 |
| ENSG00000109519 | *GRPEL1* | 1.46206E-05 | 0.073 | 0.925 |
| ENSG00000105771 | *SMG9* | 1.46648E-05 | 0.074 | 0.949 |
| ENSG00000179912 | *R3HDM2* | 1.49295E-05 | 0.075 | 0.981 |
| ENSG00000181789 | *COPG1* | 1.52107E-05 | 0.076 | 2.324 |
| ENSG00000177156 | *TALDO1* | 1.53405E-05 | 0.077 | 1.019 |
| ENSG00000174233 | *ADCY6* | 1.81383E-05 | 0.091 | 1.093 |
| ENSG00000197415 | *VEPH1* | 1.87122E-05 | 0.094 | 1.098 |
| ENSG00000173334 | *TRIB1* | 1.99623E-05 | 0.100 | 0.930 |
| ENSG00000161638 | *ITGA5* | 2.12307E-05 | 0.106 | 0.955 |
| ENSG00000273749 | *CYFIP1* | 2.32855E-05 | 0.117 | 0.750 |
| ENSG00000186866 | *POFUT2* | 2.45016E-05 | 0.123 | 0.871 |
| ENSG00000167601 | *AXL* | 2.64951E-05 | 0.133 | 1.002 |
| ENSG00000106723 | *SPIN1* | 2.6763E-05 | 0.134 | 0.911 |
| ENSG00000138160 | *KIF11* | 2.78428E-05 | 0.140 | 1.117 |
| ENSG00000156110 | *ADK* | 2.78866E-05 | 0.140 | 0.936 |
| ENSG00000100241 | *SBF1* | 2.81599E-05 | 0.141 | 1.021 |
| ENSG00000063177 | *RPL18* | 2.89518E-05 | 0.145 | 0.981 |
| ENSG00000065833 | *ME1* | 3.09363E-05 | 0.155 | 3.843 |

| Ensembl Gene ID | Official gene symbol | P value t-test 2 tailed, 2 sample unequal variance | Bonferroni p-value | Ratio (mean HHT/mean Control) |
|---|---|---|---|---|
| ENSG00000213551 | *DNAJC9* | 3.19441E-05 | 0.160 | 1.156 |
| ENSG00000163933 | *RFT1* | 3.32653E-05 | 0.167 | 0.791 |
| ENSG00000125648 | *SLC25A23* | 3.39196E-05 | 0.170 | 1.052 |
| ENSG00000130725 | *UBE2M* | 3.50599E-05 | 0.176 | 0.875 |
| ENSG00000145354 | *CISD2* | 3.54802E-05 | 0.178 | 0.907 |
| ENSG00000196396 | *PTPN1* | 3.66647E-05 | 0.184 | 1.381 |
| ENSG00000108468 | *CBX1* | 3.70687E-05 | 0.186 | 0.981 |
| ENSG00000102763 | *VWA8* | 3.96164E-05 | 0.199 | 1.102 |
| ENSG00000257923 | *CUX1* | 4.03479E-05 | 0.202 | 0.876 |
| ENSG00000106635 | *BCL7B* | 4.16653E-05 | 0.209 | 1.014 |
| ENSG00000138413 | *IDH1* | 4.21199E-05 | 0.211 | 0.733 |
| ENSG00000110104 | *CCDC86* | 4.23359E-05 | 0.212 | 1.189 |
| ENSG00000105127 | *AKAP8* | 4.26181E-05 | 0.214 | 0.925 |
| ENSG00000152953 | *STK32B* | 4.39293E-05 | 0.220 | 1.039 |
| ENSG00000100575 | *TIMM9* | 4.43954E-05 | 0.223 | 1.396 |
| ENSG00000134882 | *UBAC2* | 4.44308E-05 | 0.223 | 0.869 |
| ENSG00000109184 | *DCUN1D4* | 4.47585E-05 | 0.224 | 0.778 |
| ENSG00000106397 | *PLOD3* | 4.53711E-05 | 0.227 | 1.165 |
| ENSG00000134452 | *FBXO18* | 4.54658E-05 | 0.228 | 1.158 |
| ENSG00000168397 | *ATG4B* | 4.55588E-05 | 0.228 | 1.686 |
| ENSG00000010278 | *CD9* | 4.57309E-05 | 0.229 | 0.805 |
| ENSG00000160075 | *SSU72* | 4.8919E-05 | 0.245 | 0.896 |
| ENSG00000136270 | *TBRG4* | 5.05129E-05 | 0.253 | 0.769 |
| ENSG00000184428 | *TOP1MT* | 5.25926E-05 | 0.264 | 1.092 |
| ENSG00000132471 | *WBP2* | 5.32038E-05 | 0.267 | 1.040 |
| ENSG00000173207 | *CKS1B* | 5.40049E-05 | 0.271 | 1.473 |
| ENSG00000168067 | *MAP4K2* | 5.41645E-05 | 0.272 | 3.153 |
| ENSG00000107862 | *GBF1* | 5.51424E-05 | 0.276 | 0.819 |
| ENSG00000154359 | *LONRF1* | 5.77497E-05 | 0.289 | 0.971 |
| ENSG00000279495 | | 5.85132E-05 | 0.293 | 0.870 |
| ENSG00000112837 | *TBX18* | 6.30065E-05 | 0.316 | 0.797 |
| ENSG00000130175 | *PRKCSH* | 6.42955E-05 | 0.322 | 1.469 |
| ENSG00000110717 | *NDUFS8* | 6.59658E-05 | 0.331 | 1.085 |
| ENSG00000169902 | *TPST1* | 6.71196E-05 | 0.336 | 1.006 |
| ENSG00000127804 | *METTL16* | 6.78066E-05 | 0.340 | 0.761 |
| ENSG00000065054 | *SLC9A3R2* | 6.79157E-05 | 0.340 | 0.936 |
| ENSG00000107130 | *NCS1* | 6.7971E-05 | 0.341 | 0.185 |
| ENSG00000124216 | *SNAI1* | 6.87027E-05 | 0.344 | 0.803 |
| ENSG00000103091 | *WDR59* | 6.99565E-05 | 0.351 | 1.085 |
| ENSG00000204469 | *PRRC2A* | 7.0127E-05 | 0.352 | 0.887 |
| ENSG00000072571 | *HMMR* | 7.05599E-05 | 0.354 | 1.143 |
| ENSG00000135976 | *ANKRD36* | 7.1758E-05 | 0.360 | 1.181 |
| ENSG00000162129 | *CLPB* | 7.25711E-05 | 0.364 | 1.020 |
| ENSG00000136450 | *SRSF1* | 7.27345E-05 | 0.365 | 0.796 |
| ENSG00000140400 | *MAN2C1* | 7.32183E-05 | 0.367 | 1.376 |
| ENSG00000204713 | *TRIM27* | 7.32339E-05 | 0.367 | 0.643 |
| ENSG00000198690 | *FAN1* | 7.62791E-05 | 0.382 | 1.223 |
| ENSG00000100600 | *LGMN* | 7.6541E-05 | 0.384 | 0.745 |
| ENSG00000103507 | *BCKDK* | 7.79359E-05 | 0.391 | 0.963 |
| ENSG00000101019 | *UQCC1* | 7.82885E-05 | 0.392 | 0.964 |
| ENSG00000108469 | *RECQL5* | 7.96618E-05 | 0.399 | 1.023 |
| ENSG00000185033 | *SEMA4B* | 8.08788E-05 | 0.405 | 1.383 |
| ENSG00000172270 | *BSG* | 8.17753E-05 | 0.410 | 0.765 |
| ENSG00000140740 | *UQCRC2* | 8.25239E-05 | 0.414 | 0.719 |
| ENSG00000117592 | *PRDX6* | 8.75153E-05 | 0.439 | 0.975 |
| ENSG00000175376 | *EIF1AD* | 8.88598E-05 | 0.445 | 1.501 |
| ENSG00000100162 | *CENPM* | 8.93274E-05 | 0.448 | 1.162 |
| ENSG00000186594 | *MIR22HG* | 9.01266E-05 | 0.452 | 1.270 |
| ENSG00000136240 | *KDELR2* | 9.22001E-05 | 0.462 | 0.829 |

| Ensembl Gene ID | Official gene symbol | P value t-test 2 tailed, 2 sample unequal variance | Bonferroni p-value for 5,013 genes | Ratio (mean HHT/mean Control) |
|---|---|---|---|---|
| ENSG00000198885 | *ITPRIPL1* | 9.86812E-05 | 0.495 | 3.105 |
| ENSG00000187840 | *EIF4EBP1* | 0.000100507 | 0.504 | 1.810 |
| ENSG00000214756 | *METTL12* | 0.00010193 | 0.511 | 1.269 |
| ENSG00000110063 | *DCPS* | 0.000105002 | 0.526 | 4.329 |
| ENSG00000065000 | *AP3D1* | 0.000105414 | 0.528 | 0.219 |
| ENSG00000103275 | *UBE2I* | 0.000107785 | 0.540 | 1.152 |
| ENSG00000095319 | *NUP188* | 0.000109405 | 0.548 | 0.664 |
| ENSG00000130985 | *UBA1* | 0.000110682 | 0.555 | 1.239 |
| ENSG00000012983 | *MAP4K5* | 0.000111763 | 0.560 | 0.914 |
| ENSG00000205707 | *LYRM5* | 0.000114379 | 0.573 | 0.934 |
| ENSG00000205339 | *IPO7* | 0.000114776 | 0.575 | 0.486 |
| ENSG00000125733 | *TRIP10* | 0.000115467 | 0.579 | 0.788 |
| ENSG00000138834 | *MAPK8IP3* | 0.000116995 | 0.586 | 1.074 |
| ENSG00000134107 | *BHLHE40* | 0.000117052 | 0.587 | 1.179 |
| ENSG00000215305 | *VPS16* | 0.000118167 | 0.592 | 0.734 |
| ENSG00000187109 | *NAP1L1* | 0.000119748 | 0.600 | 1.156 |
| ENSG00000280071 | | 0.000120359 | 0.603 | 1.020 |
| ENSG00000162623 | *TYW3* | 0.000121446 | 0.609 | 0.896 |
| ENSG00000172661 | *FAM21C* | 0.000125868 | 0.631 | 0.906 |
| ENSG00000170873 | *MTSS1* | 0.000126813 | 0.636 | 1.054 |
| ENSG00000133612 | *AGAP3* | 0.00012721 | 0.638 | 0.795 |
| ENSG00000115568 | *ZNF142* | 0.000130816 | 0.656 | 1.593 |
| ENSG00000136026 | *CKAP4* | 0.000135342 | 0.678 | 0.784 |
| ENSG00000167721 | *TSR1* | 0.000135675 | 0.680 | 1.001 |
| ENSG00000154134 | *ROBO3* | 0.000136066 | 0.682 | 1.217 |
| ENSG00000131435 | *PDLIM4* | 0.000138415 | 0.694 | 0.820 |
| ENSG00000163874 | *ZC3H12A* | 0.000140939 | 0.707 | 0.914 |
| ENSG00000078177 | *N4BP2* | 0.000141725 | 0.710 | 0.659 |
| ENSG00000135829 | *DHX9* | 0.000143014 | 0.717 | 0.677 |
| ENSG00000126746 | *ZNF384* | 0.000143715 | 0.720 | 0.982 |
| ENSG00000175573 | *C11orf68* | 0.000146388 | 0.734 | 1.128 |
| ENSG00000179409 | *GEMIN4* | 0.000146849 | 0.736 | 1.241 |
| ENSG00000184009 | *ACTG1* | 0.000146948 | 0.737 | 1.062 |
| ENSG00000147454 | *SLC25A37* | 0.000149271 | 0.748 | 0.974 |
| ENSG00000177426 | *TGIF1* | 0.000152671 | 0.765 | 1.098 |
| ENSG00000105397 | *TYK2* | 0.000157287 | 0.788 | 0.814 |
| ENSG00000147677 | *EIF3H* | 0.000158103 | 0.793 | 0.999 |
| ENSG00000049246 | *PER3* | 0.000158961 | 0.797 | 1.755 |
| ENSG00000156990 | *RPUSD3* | 0.000159667 | 0.800 | 1.609 |
| ENSG00000135801 | *TAF5L* | 0.00016119 | 0.808 | 0.807 |
| ENSG00000130396 | *MLLT4* | 0.000162828 | 0.816 | 0.971 |
| ENSG00000133027 | *PEMT* | 0.00016379 | 0.821 | 0.951 |
| ENSG00000138095 | *LRPPRC* | 0.000166519 | 0.835 | 2.527 |
| ENSG00000106211 | *HSPB1* | 0.000170639 | 0.855 | 2.792 |
| ENSG00000108344 | *PSMD3* | 0.000173172 | 0.868 | 1.187 |
| ENSG00000113070 | *HBEGF* | 0.000175151 | 0.878 | 0.991 |
| ENSG00000169554 | *ZEB2* | 0.000175298 | 0.879 | 1.108 |
| ENSG00000127993 | *RBM48* | 0.000181705 | 0.911 | 1.020 |
| ENSG00000205250 | *E2F4* | 0.000182827 | 0.917 | 1.324 |
| ENSG00000148248 | *SURF4* | 0.000184866 | 0.927 | 0.630 |
| ENSG00000099797 | *TECR* | 0.000186644 | 0.936 | 0.810 |
| ENSG00000143157 | *POGK* | 0.000186797 | 0.936 | 0.918 |
| ENSG00000168675 | *LDLRAD4* | 0.000193667 | 0.971 | 1.028 |

**Supplementary Table 4:**
**Top 1000 differentially expressed genes (normalized to total counts per library)**

Full list supplied after Supplementary References, at end of Data Supplement

#### Supplementary Table 5: Gene Ontology terms enriched by clustering  top 100-1,000 Gene IDs meeting CV10

| | Number of top-ranked Ensembl gene IDs clustered | | | | | | | |
|---|---|---|---|---|---|---|---|---|
| **Significant Gene Ontology (GO) terms by function** | **100** | **200** | **300** | **400** | **500** | **600** | **750** | **1000** |
| **PROTEIN FOLDING** | | | | | | | | |
| GOTERM_BP_DIRECT GO:0006457~protein folding | 100 | 200 | 300 | 400 | 500 | 600 | 750 | 1000 |
| **EXTRACELLULAR MATRIX** | | | | | | | | |
| GOTERM_BP_DIRECT GO:0031012~extracellular matrix | 100 | 200 | 300 | 400 | 500 | 600 | 750 | 1000 |
| GOTERM_MF_DIRECT GO:0005201~extracellular matrix structural constituent |  | 200 | 300 | 400 | 500 | 600 | 750 | 1000 |
| GOTERM_BP_DIRECT GO:0030198~extracellular matrix organization | 100 |  |  |  |  |  |  |  |
| **ENERGY** | | | | | | | | |
| GOTERM_MF_DIRECT GO:0005524~ATP binding | 100 |  |  |  |  | 600 | 750 | 1000 |
| GOTERM_BP_DIRECT GO:0006099~tricarboxylic acid cycle |  | 200 | 300 | 400 | 500 | 600 | 750 | 1000 |
| GOTERM_BP_DIRECT GO:0006096~glycolytic process |  |  | 300 | 400 | 500 | 600 | 750 | 1000 |
| GOTERM_BP_DIRECT GO:0006102~isocitrate metabolic process |  |  |  |  |  |  | 750 | 1000 |
| GOTERM_MF_DIRECT GO:0051287~NAD binding |  |  |  |  |  |  | 750 | 1000 |
| GOTERM_MF_DIRECT GO:0004553~hydrolase activity, hydrolyzing O-glycosyl compounds |  |  |  |  |  |  |  | 1000 |
| **PROTEIN ENZYMATIC CHANGES (not glycoslylation)** | | | | | | | | |
| GOTERM_MF_DIRECT GO:0003756~protein disulfide isomerase activity |  | 200 | 300 | 400 | 500 | 600 | 750 | 1000 |
| GOTERM_BP_DIRECT GO:0017185~peptidyl-lysine hydroxylation |  |  |  |  |  | 600 | 750 | 1000 |
| GOTERM_MF_DIRECT GO:0008475~procollagen-lysine 5-dioxygenase activity |  |  |  |  |  | 600 | 750 | 1000 |
| **MITOCHONDRIA** | | | | | | | | |
| GOTERM_CC_DIRECT GO:0005739~mitochondrion |  |  | 300 | 400 | 500 | 600 | 750 | 1000 |
| GOTERM_CC_DIRECT GO:0005759~mitochondrial matrix |  |  | 300 | 400 | 500 | 600 | 750 | 1000 |
| **PROTEIN BINDING** | | | | | | | | |
| GOTERM_MF_DIRECT GO:0048156~tau protein binding | 100 | 200 |  |  |  |  |  |  |
| GOTERM_MF_DIRECT GO:0031072~heat shock protein binding | 100 | 200 |  |  |  |  |  |  |
| GOTERM_MF_DIRECT GO:0031418~L-ascorbic acid binding |  |  |  |  |  | 600 | 750 | 1000 |
| GOTERM_MF_DIRECT GO:0005178~integrin binding | 100 |  |  |  | 500 |  |  |  |
| **CIRCADIAN RHYTHM** | | | | | | | | |
| GOTERM_BP_DIRECT GO:0042752~regulation of circadian rhythm |  |  |  | 400 |  | 600 | 750 | 1000 |
| GOTERM_BP_DIRECT GO:0032922~circadian regulation of gene expression |  |  |  | 400 | 500 | 600 | 750 | 1000 |
| **SIGNALING PATHWAYS REGULATING RAS, GTPASES, SRC KINASE** | | | | | | | | |
| GOTERM_MF_DIRECT GO:0017124~SH3 domain binding |  |  |  |  | 500 | 600 | 750 |  |
| GOTERM_MF_DIRECT GO:0005085~guanyl-nucleotide exchange factor activity |  |  |  | 400 | 500 | 600 | 750 | 1000 |
| GOTERM_BP_DIRECT GO:0051056~regulation of small GTPase mediated signal transduction |  |  |  |  |  |  | 750 |  |
| GOTERM_BP_DIRECT GO:0006468~protein phosphorylation |  |  |  |  |  |  | 750 | 1000 |
| GOTERM_MF_DIRECT GO:0004712~protein serine/threonine/tyrosine kinase activity |  |  |  |  |  |  |  | 1000 |
| GOTERM_MF_DIRECT GO:0004672~protein kinase activity |  |  |  |  |  |  |  | 1000 |
| GOTERM_BP_DIRECT GO:0007266~Rho protein signal transduction |  |  |  |  |  |  |  | 1000 |
| **GROWTH FACTOR/CELL DIVISION** | | | | | | | | |
| GOTERM_MF_DIRECT GO:0008083~growth factor activity |  | 200 |  | 400 |  |  | 750 | 1000 |
| GOTERM_BP_DIRECT GO:0051781~positive regulation of cell division |  |  |  | 400 | 500 |  | 750 | 1000 |
| GOTERM_BP_DIRECT GO:0009611~response to wounding |  |  |  | 400 |  |  |  |  |

| ……./ GO terms by function | Number of top-ranked Ensembl gene IDs clustered | | | | | | | |
|---|---|---|---|---|---|---|---|---|
| | 100 | 200 | 300 | 400 | 500 | 600 | 750 | 1000 |
| **CYTOSKELETON** | | | | | | | | |
| GOTERM_MF_DIRECT GO:0005200~structural constituent of cytoskeleton | | 200 | | | | | | |
| GOTERM_MF_DIRECT GO:0008092~cytoskeletal protein binding | | 200 | 300 | | | | 750 | 1000 |
| GOTERM_CC_DIRECT GO:0001725~stress fiber | | | | | | 600 | 750 | 1000 |
| GOTERM_BP_DIRECT GO:0031274~positive regulation of pseudopodium assembly | | | | | | | | 1000 |
| **AMINO ACIDS** | | | | | | | | |
| GOTERM_BP_DIRECT GO:0006541~glutamine metabolic process | | | 300 | | | | 750 | 1000 |
| **VIRAL ENTRY INTO HOST CELL** | | | | | | | | |
| GOTERM_BP_DIRECT GO:0046718~viral entry into host cell | | 200 | | | 500 | | 750 | |
| GOTERM_MF_DIRECT GO:0001618~virus receptor activity | | 200 | | 400 | 500 | | 750 | |
| **TRANSCRIPTION** | | | | | | | | |
| GOTERM_MF_DIRECT GO:0008134~transcription factor binding | | | | | 500 | 600 | | |
| GOTERM_MF_DIRECT GO:0061629~RNA pol. II sequence-specific DNA binding transcription factor binding | | | | | 500 | 600 | | |
| GOTERM_CC_DIRECT GO:0005675~holo TFIIH complex | | | | | | | 750 | |
| GOTERM_CC_DIRECT GO:0090575~RNA polymerase II transcription factor complex | | | | | | | | 1000 |
| **GLYCOSYLATION** | | | | | | | | |
| GOTERM_BP_DIRECT GO:0006013~mannose metabolic process | | | | | | 600 | 750 | 1000 |
| GOTERM_BP_DIRECT GO:0004571~mannosyl-oligosaccharide 1,2-alpha-mannosidase activity | | | | | | | 750 | 1000 |
| GOTERM_BP_DIRECT GO:0006491~N-glycan processing | | | | | | 600 | 750 | 1000 |
| GOTERM_BP_DIRECT GO:0016266~O-glycan processing | | | | | | 600 | | |
| GOTERM_BP_DIRECT GO:0006486~protein glycosylation | | | | | | 600 | | 1000 |
| **CELLULAR RESPONSE TO STRESS** | | | | | | | | |
| GOTERM_BP_DIRECT GO:0034976~response to endoplasmic reticulum stress | | | | | 500 | 600 | | |
| GOTERM_BP_DIRECT GO:0042149~cellular response to glucose starvation | | | | 400 | | | | |
| **CCYTOKINES/ NFKB** | | | | | | | | |
| GOTERM_BP_DIRECT GO:0070555~response to interleukin-1 | | | 300 | 400 | | | | |
| GOTERM_BP_DIRECT GO:1901224~positive regulation of NIK/NF-kappaB signaling | | | | 400 | | | | |
| **IRON** | | | | | | | | |
| GOTERM_BP_DIRECT GO:0055072~iron ion homeostasis | | | | | | | 750 | 1000 |
| **tRNAs** | | | | | | | | |
| GOTERM_MF_DIRECT GO:0002161~aminoacyl-tRNA editing activity | | | | | | 600 | 750 | |
| GOTERM_MF_DIRECT GO:0000049~tRNA binding | | | | | | | 750 | |
| **LIGAND-DEPENDENT RECEPTOR NUCLEAR BINDING** | | | | | | | | |
| GOTERM_MF_DIRECT GO:0016922~ligand-dependent nuclear receptor binding | | | | | | | | 1000 |
| **TGF-β** | | | | | | | | |
| GOTERM_MF_DIRECT GO:0050431~transforming growth factor beta binding | | | | | | | | 1000 |
| **BASEMENT MEMBRANE** | | | | | | | | |
| GOTERM_CC_DIRECT GO:0005604~basement membrane | | | | | | | | 1000 |
| **ENDOPLASMIC RETICULUM LUMEN** | | | | | | | | |
| GOTERM_CC_DIRECT GO:0033116~endoplasmic reticulum-Golgi intermediate compartment membrane | | | | | | | 750 | |
| GOTERM_CC_DIRECT GO:0005793~endoplasmic reticulum-Golgi intermediate compartment | | | | | | | 750 | |

| **……./ GO terms by function** | **Number of top-ranked Ensembl gene IDs clustered** | | | | | | | |
|---|---|---|---|---|---|---|---|---|
| | **100** | **200** | **300** | **400** | **500** | **600** | **750** | **1000** |
| **FOCAL ADHESION** | | | | | | | | |
| GOTERM_CC_DIRECT GO:0005925~focal adhesion | 100 | | | | | | | |
| GOTERM_CC_DIRECT GO:0005911~cell-cell junction | 100 | | | | | | | |
| **MISCELLANEOUS** | | | | | | | | |
| GOTERM_CC_DIRECT GO:0098685~Schaffer collateral | 100 | | | | | | | |
| GOTERM_BP_DIRECT GO:0043525~positive regulation of neuron apoptotic process | 100 | | | | | | | |
| GOTERM_BP_DIRECT GO:0030593~neutrophil chemotaxis | 100 | | | | | | | |
| GOTERM_BP_DIRECT GO:0009410~response to xenobiotic stimulus | 100 | | | | | | | |
| GOTERM_BP_DIRECT GO:0098974~postsynaptic actin cytoskeleton organization | | 200 | | | | | | |

#### Supplementary Table 6: Gene Ontology terms enriched clustering 10 sets of 1000 random genes

10 randomly selected datasets of 1,000 Ensembl[35] Gene IDs from the current BOEC dataset of 16,807 were clustered using DAVID[29] to indicate gene ontology (GO) terms[27,28] that should be interpreted with caution if clustered in differential gene alignment studies. Supplementary Table 6A indicates the number of clusters generated by each random selection of 1,000 Ensembl Gene IDs with alignments in the BOEC datasets. Supplementary Table 6B list details the individual GO terms that were generated across the random datasets, where "significant" was defined by GO term p<0.05.

##### Supplementary Table 6A: Overview

| | 1000 #1 | 1000 #2 | 1000 #3 | 1000 #4 | 1000 #5 | 1000 #6 | 1000 #7 | 1000 #8 | 1000 #9 | 1000 #10 |
|---|---|---|---|---|---|---|---|---|---|---|
| Number of GO terms | | | | | | | | | | |
| Clustered | 11 | 11 | 19 | 6 | 12 | 8 | 11 | 16 | 37 | 13 |
| Significant | 4 | 4 | 6 | 4 | 8 | 6 | 8 | 6 | 11 | 6 |

##### Supplementary Table 6B: Details

| GO terms clustered in one or more of 10 random datasets of 1,000/16,807 Ensemble Gene IDs with alignments in current study BOECs | | Number of the 10 random sets where stated term was | |
|---|---|---|---|
| | | Clustered | Clustered |
| GOTERM_BP_DIRECT | GO:0000398~mRNA splicing, via spliceosome | 1 | 1 |
| GOTERM_BP_DIRECT | GO:0000722~telomere maintenance via recombination | 1 | 0 |
| GOTERM_BP_DIRECT | GO:0006024~glycosaminoglycan biosynthetic process | 1 | 0 |
| GOTERM_BP_DIRECT | GO:0006139~nucleobase-containing compound metabolic process | 1 | 0 |
| GOTERM_BP_DIRECT | GO:0006260~DNA replication | 1 | 0 |
| GOTERM_BP_DIRECT | GO:0006281~DNA repair | 1 | 1 |
| GOTERM_BP_DIRECT | GO:0006351~transcription, DNA-templated | 5 | 2 |
| GOTERM_BP_DIRECT | GO:0006355~regulation of transcription, DNA-templated | 5 | 3 |
| GOTERM_BP_DIRECT | GO:0006364~rRNA processing 20 | 1 | 1 |
| GOTERM_BP_DIRECT | GO:0006367~transcription initiation from RNA polymerase II promoter | 1 | 1 |
| GOTERM_BP_DIRECT | GO:0006368~transcription elongation from RNA polymerase II promoter | 1 | 1 |
| GOTERM_BP_DIRECT | GO:0006370~7-methylguanosine mRNA capping | 1 | 1 |
| GOTERM_BP_DIRECT | GO:0006397~mRNA processing | 1 | 1 |
| GOTERM_BP_DIRECT | GO:0006412~translation | 1 | 1 |
| GOTERM_BP_DIRECT | GO:0006413~translational initiation | 1 | 1 |
| GOTERM_BP_DIRECT | GO:0006418~tRNA aminoacylation for protein translation | 1 | 1 |
| GOTERM_BP_DIRECT | GO:0006468~protein phosphorylation | 4 | 2 |
| GOTERM_BP_DIRECT | GO:0006470~protein dephosphorylation | 1 | 1 |
| GOTERM_BP_DIRECT | GO:0006614~SRP-dependent cotranslational protein targeting to membrane | 1 | 0 |
| GOTERM_BP_DIRECT | GO:0006783~heme biosynthetic process | 1 | 0 |
| GOTERM_BP_DIRECT | GO:0006814~sodium ion transport | 1 | 1 |
| GOTERM_BP_DIRECT | GO:0007062~sister chromatid cohesion | 1 | 1 |
| GOTERM_BP_DIRECT | GO:0007067~mitotic nuclear division | 2 | 2 |
| GOTERM_BP_DIRECT | GO:0007156~homophilic cell adhesion via plasma membrane adhesion molecules | 1 | 0 |
| GOTERM_BP_DIRECT | GO:0007623~circadian rhythm | 1 | 0 |
| GOTERM_BP_DIRECT | GO:0008380~RNA splicing | 1 | 0 |
| GOTERM_BP_DIRECT | GO:0008654~phospholipid biosynthetic process | 1 | 0 |
| GOTERM_BP_DIRECT | GO:0009083~branched-chain amino acid catabolic process | 1 | 1 |
| GOTERM_BP_DIRECT | GO:0015012~heparan sulfate proteoglycan biosynthetic process | 1 | 1 |
| GOTERM_BP_DIRECT | GO:0015949~nucleobase-containing small molecule interconversion | 1 | 0 |
| GOTERM_BP_DIRECT | GO:0016569~covalent chromatin modification | 1 | 0 |
| GOTERM_BP_DIRECT | GO:0018105~peptidyl-serine phosphorylation | 2 | 0 |
| GOTERM_BP_DIRECT | GO:0019083~viral transcription | 1 | 1 |
| GOTERM_BP_DIRECT | GO:0030206~chondroitin sulfate biosynthetic process | 1 | 0 |
| GOTERM_BP_DIRECT | GO:0030518~intracellular steroid hormone receptor signaling pathway | 1 | 0 |
| GOTERM_BP_DIRECT | GO:0030521~androgen receptor signaling pathway | 2 | 0 |
| GOTERM_BP_DIRECT | GO:0032057~negative regulation of translational initiation in response to stress | 1 | 0 |
| GOTERM_BP_DIRECT | GO:0034198~cellular response to amino acid starvation | 1 | 0 |
| GOTERM_BP_DIRECT | GO:0034644~cellular response to UV | 1 | 1 |
| GOTERM_BP_DIRECT | GO:0035335~peptidyl-tyrosine dephosphorylation | 1 | 0 |
| GOTERM_BP_DIRECT | GO:0035556~intracellular signal transduction | 1 | 0 …../ |

| ……./ GO terms clustered in one or more of 10 random datasets of 1,000/16,807 Ensemble Gene IDs with alignments in current study BOECs | | Number of the 10 random sets where stated term was | |
|---|---|---|---|
| | | Clustered | Clustered, p<0.05 |
| GOTERM_BP_DIRECT | GO:0035725~sodium ion transmembrane transport | 1 | 0 |
| GOTERM_BP_DIRECT | GO:0040008~regulation of growth | 1 | 1 |
| GOTERM_BP_DIRECT | GO:0042795~snRNA transcription from RNA polymerase II promoter | 1 | 0 |
| GOTERM_BP_DIRECT | GO:0046426~negative regulation of JAK-STAT cascade | 1 | 0 |
| GOTERM_BP_DIRECT | GO:0050434~positive regulation of viral transcription | 1 | 0 |
| GOTERM_BP_DIRECT | GO:0051056~regulation of small GTPase mediated signal transduction | 1 | 1 |
| GOTERM_BP_DIRECT | GO:0051092~positive regulation of NF-kappaB transcription factor activity | 1 | 0 |
| GOTERM_BP_DIRECT | GO:0051301~cell division | 2 | 1 |
| GOTERM_BP_DIRECT | GO:0098609~cell-cell adhesion | 3 | 2 |
| GOTERM_CC_DIRECT | GO:0000151~ubiquitin ligase complex | 1 | 0 |
| GOTERM_CC_DIRECT | GO:0000776~kinetochore | 1 | 0 |
| GOTERM_CC_DIRECT | GO:0000777~condensed chromosome kinetochore | 1 | 0 |
| GOTERM_CC_DIRECT | GO:0000790~nuclear chromatin | 1 | 1 |
| GOTERM_CC_DIRECT | GO:0005634~nucleus | 1 | 1 |
| GOTERM_CC_DIRECT | GO:0005669~transcription factor TFIID complex | 1 | 1 |
| GOTERM_CC_DIRECT | GO:0005681~spliceosomal complex | 1 | 0 |
| GOTERM_CC_DIRECT | GO:0005759~mitochondrial matrix | 1 | 1 |
| GOTERM_CC_DIRECT | GO:0005840~ribosome | 1 | 0 |
| GOTERM_CC_DIRECT | GO:0005913~cell-cell adherens junction | 3 | 3 |
| GOTERM_CC_DIRECT | GO:0016592~mediator complex | 2 | 1 |
| GOTERM_CC_DIRECT | GO:0016607~nuclear speck | 1 | 0 |
| GOTERM_CC_DIRECT | GO:0022625~cytosolic large ribosomal subunit | 1 | 0 |
| GOTERM_CC_DIRECT | GO:0030054~cell junction | 1 | 0 |
| GOTERM_CC_DIRECT | GO:0071013~catalytic step 2 spliceosome | 2 | 1 |
| GOTERM_MF_DIRECT | GO:0000049~tRNA binding | 1 | 0 |
| GOTERM_MF_DIRECT | GO:0000166~nucleotide binding | 1 | 1 |
| GOTERM_MF_DIRECT | GO:0001104~RNA polymerase II transcription cofactor activity | 2 | 0 |
| GOTERM_MF_DIRECT | GO:0003676~nucleic acid binding | 3 | 0 |
| GOTERM_MF_DIRECT | GO:0003677~DNA binding | 3 | 2 |
| GOTERM_MF_DIRECT | GO:0003700~transcription factor activity, sequence-specific DNA binding | 2 | 1 |
| GOTERM_MF_DIRECT | GO:0003712~transcription cofactor activity | 1 | 0 |
| GOTERM_MF_DIRECT | GO:0003735~structural constituent of ribosome | 1 | 1 |
| GOTERM_MF_DIRECT | GO:0004402~histone acetyltransferase activity | 1 | 0 |
| GOTERM_MF_DIRECT | GO:0004550~nucleoside diphosphate kinase activity | 1 | 1 |
| GOTERM_MF_DIRECT | GO:0004672~protein kinase activity | 4 | 1 |
| GOTERM_MF_DIRECT | GO:0004674~protein serine/threonine kinase activity | 4 | 0 |
| GOTERM_MF_DIRECT | GO:0004697~protein kinase C activity | 1 | 1 |
| GOTERM_MF_DIRECT | GO:0004725~protein tyrosine phosphatase activity | 1 | 0 |
| GOTERM_MF_DIRECT | GO:0004872~receptor activity | 1 | 0 |
| GOTERM_MF_DIRECT | GO:0005096~GTPase activator activity | 1 | 0 |
| GOTERM_MF_DIRECT | GO:0005524~ATP binding | 5 | 5 |
| GOTERM_MF_DIRECT | GO:0008270~zinc ion binding | 1 | 0 |
| GOTERM_MF_DIRECT | GO:0008375~acetylglucosaminyltransferase activity | 1 | 1 |
| GOTERM_MF_DIRECT | GO:0016922~ligand-dependent nuclear receptor binding | 1 | 0 |
| GOTERM_MF_DIRECT | GO:0019205~nucleobase-containing compound kinase activity | 1 | 1 |
| GOTERM_MF_DIRECT | GO:0030374~ligand-dependent nuclear receptor transcription coactivator activity | 3 | 2 |
| GOTERM_MF_DIRECT | GO:0031404~chloride ion binding | 1 | 0 |
| GOTERM_MF_DIRECT | GO:0035257~nuclear hormone receptor binding | 1 | 1 |
| GOTERM_MF_DIRECT | GO:0042809~vitamin D receptor binding | 1 | 1 |
| GOTERM_MF_DIRECT | GO:0046872~metal ion binding | 5 | 4 |
| GOTERM_MF_DIRECT | GO:0046966~thyroid hormone receptor binding | 1 | 1 |
| GOTERM_MF_DIRECT | GO:0050660~flavin adenine dinucleotide binding | 1 | 0 |
| GOTERM_MF_DIRECT | GO:0050681~androgen receptor binding | 1 | 0 |
| GOTERM_MF_DIRECT | GO:0061630~ubiquitin protein ligase activity | 1 | 0 |
| GOTERM_MF_DIRECT | GO:0098641~cadherin binding involved in cell-cell adhesion | 3 | 3 |

#### Supplementary Table 7: PTC sequence contexts

| | *ACVRL1*$^{+/PTC391}$ | *ENG*$^{+/PTC93}$ | *ENG*$^{+/PTC436}$ | *SMAD4*$^{+/PTC368}$ |
|---|---|---|---|---|
| **i) DNA Variant Site** | | | | |
| *GRCh38/hg38* | 12:51,916,158 | 9:127,829,769 | 9:127,819,627 | 18:51,065,563 |
| *GRCh37/hg19* | 12:52,309,942 | 9:130,592,049 | 9:130,581,906 | 18:48,593,411 |
| Ensembl transcript | ENSG00000139567 | ENSG00000106991 | ENSG00000106991 | ENSG00000141646 |
| NCBI Reference Sequence | NM_000020.3 | NM_001114753.3 | NM_001114753.3 | NM_005359.6 |
| Exon context | Exon 8 of 10 | Exon 3 of 15 | Exon 10 of 15 | Exon 10 of 12 |
| Single nucleotide variant (SNV) | c.1171G>T | c.277C>T | c.1306C>T | c.1096C>T |
| Wildtype amino acid (αα) | Glu(E)391 | Arg(R)93 | Gln(Q)436 | Gln(Q)368 |
| Potential SNV missense αα | 6: A,G,Q,N,V | 4: Q,E,L,P | 6:E,H,L,K,P,R | 6:E,H,L,K,P,R |
| HHT pathogenic missense variant | - | - | - | - |
| VUS/benign missense variant | - | rs532649202 (Q)* | - | - |
| Synonymous variant | rs750157791 (E)* | - | - | - |
| **ii) PTC (Stop codon) read-through features** | | | | |
| PTC sequence | UAG | UGA | UAG | UAA |
| PTC context | U**<u>UAG</u>**U | C**<u>UGA</u>**G | A**<u>UAG</u>**C | U**<u>UAA</u>**C |
| No (%) AU in 3' 15 nucleotides | 8 (53%) | 6 (40%) | 7 (47%) | 8 (53%) |
| Nucleotides to 3' exon-exon boundary | 75 | 83 | 5 | 43 |
| Nucleotides from polyA site | 539 | 2004 | 975 | 1102 |
| Most likely readthrough αα | Y Tyrosine | W Tryptophan | Y Tyrosine | Y Tyrosine |
| Grantham distance from wildtype | 122 | 101 | 99 | 99 |
| **iii) Ribosomal reinitiation considerations** | | | | |
| Interval to next 3' AUG | 70 nts | 208 nts | 24 nts | 7 nts |
| Reading frame for next 3' AUG | +1 | +1 | In-frame | +1 |
| 3' protein homodimerization | No | Yes | Yes | No |
| RNASeq for potential modifiers | | | | |
| *ABCE1* (% of control BOECs | 75% | 99% | | 88% |
| *EIF3J* (% of control BOECs) | 78% | 69% | - | 72% |
| *EIF2D* (% of control BOECs) | 80% | 61% | - | 75% |
| *DENR* (% of control BOECs) | 98% | 90% | - | 112% |

**i)** DNA variant sites for the 4 PTCs examined by site, wildtype aminoacid, potential natural missense substitutions, and reported aminoacid substitutions (missense or synonymous variants at codon).[36-39] * allele frequency <$10^{-5}$.[36,39]
**ii)** PTC readthrough-relevant[16,17,40,41] comparative characteristics. αα: aminoacid; Green and red boxes indicate strongest and weakest per row.
**iii)** Ribosomal readthrough-relevant[16,17,40,41] comparative characteristics of the PTCs, with strongest and weakest readthrough enhancers by row indicated by green and red boxes respectively. RNASeq expression for well-known ribosome splitting/recycling factors,[42,43] shown as % of control.

**Supplementary Fig. 1. Blood outgrowth endothelial cell (BOEC) derivation indices**

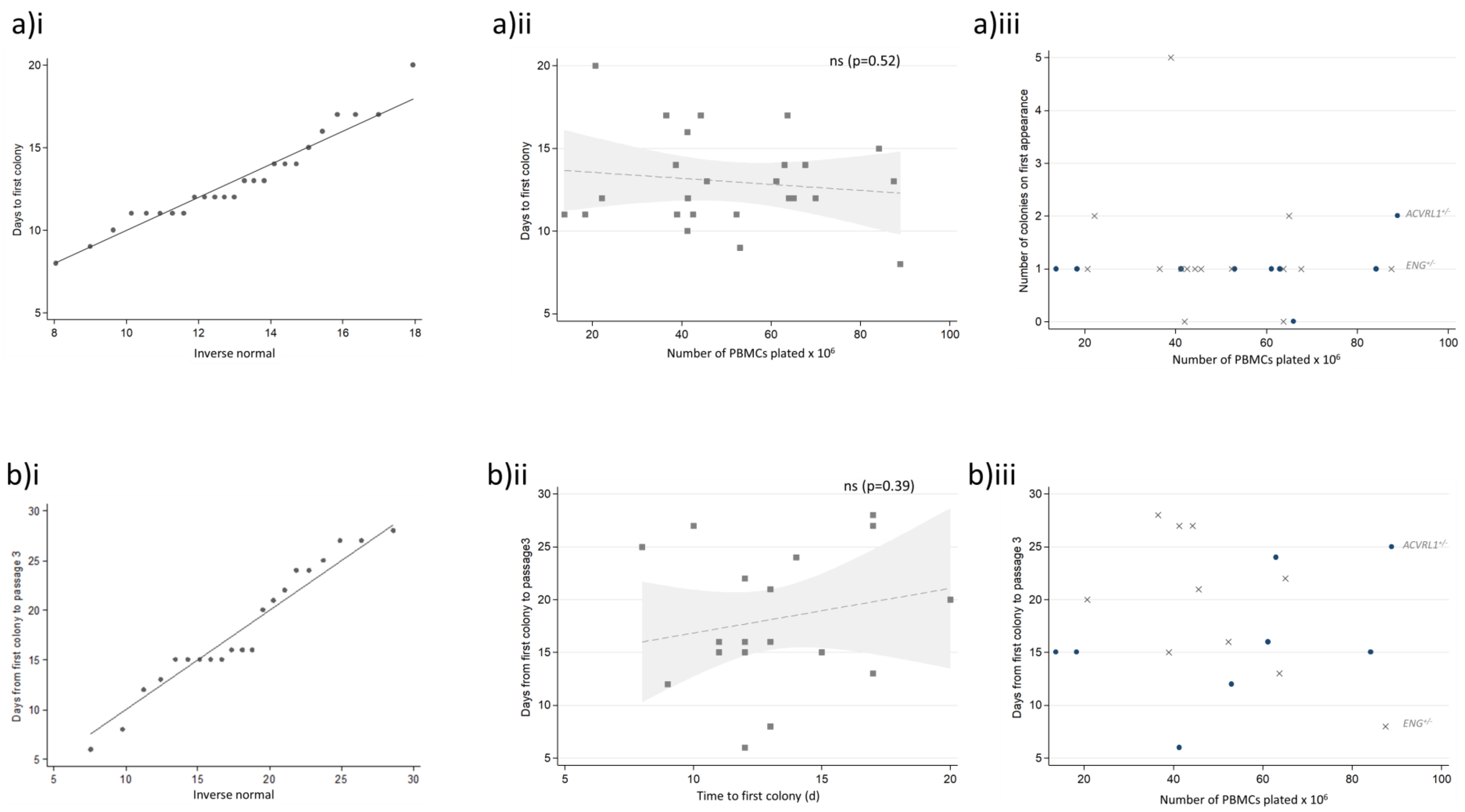

Derivation of control, *ACVRL1*$^{+/-}$, *ENG*$^{+/-}$ and *SMAD4*$^{+/-}$ BOECs. **A)** First colony appearance in the first 25 successful establishments. **(i)** Normal quantile plot indicating that the time to first colony/ies approximated to a normal distribution. (**ii**) Two-way scatter plot of number of PBMCs plated and time to first colony across all genotypes (control and HHT), p value calculated by linear regression. (**iii**) Two-way scatter plot of number of colonies emerging versus number of PBMCs plated, categorized by HHT genotype (*ACVRL1*$^{+/-}$ navy circles*;* *ENG*$^{+/-}$ grey crosses). Note the broad variability and no relationship with HHT genotype. **B)** Time from first colony appearance to passage 3, representing a rate of proliferation index. **(i)** Normal quantile plot indicating that the time approximated to a normal distribution. (**ii**) Two-way scatter plot of number of time to first colony, and time from first colony appearance to passage 3 across all genotypes (control and HHT), p value calculated by linear regression. (**iii**) Two-way scatter plot of time from first colony appearance to passage 3 versus number of PBMCs plated, categorized by HHT genotype (*ACVRL1*$^{+/-}$ navy circles*;* *ENG*$^{+/-}$ grey crosses). Note the broad variability and no relationship with HHT genotype.

#### Supplementary Fig. 2: Endothelial and RNASeq validations of blood outgrowth endothelial cells

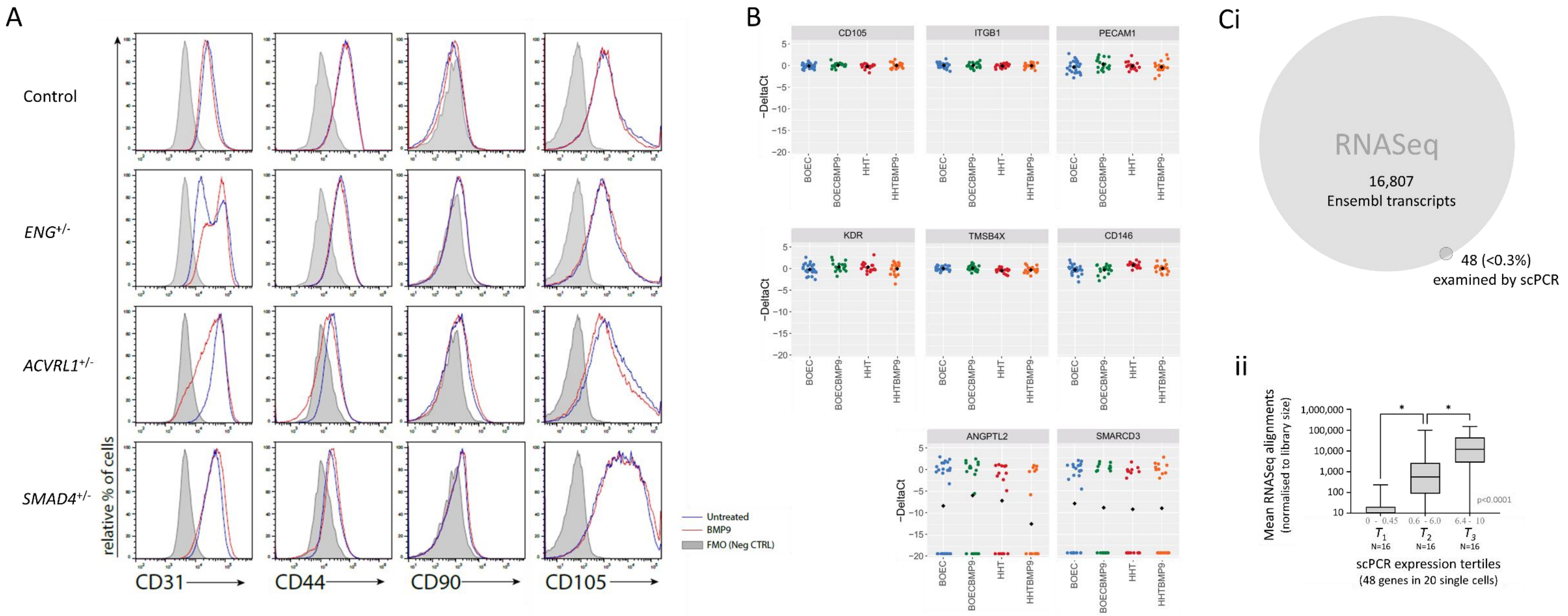

**A) Bulk flow cytometric evaluations of the control *ENG*$^{+/-}$, *ACVRL1*$^{+/-}$ and *SMAD4*$^{+/-}$ BOECs** sent for RNASeq, simultaneously examining CD31 (FITC-conjugated), CD44 (APC conjugated), CD90 (BV510-conjugated), and CD105 (PE-conjugated). Full antibody details are available on request. There is no evidence that CD31 (PECAM1), the most abundant endothelial surface receptor, displays altered expression in HHT, and it was used as the control for ENG expression in earlier studies.[30,44,45] CD44 is not referred to in the HHT literature but is a core endothelial cell surface marker which regulates endothelial functions through a CD31/PECAM1 dependent mechanism. CD90 (Thy1) is a mesenchymal stem cell marker used as a negative control. CD105 (endoglin) displays varying expression levels and there is evidence in the HHT literature[46] that it is reduced on BOECs with *ACVRL1* as well as *ENG* pathogenic variants (*ENG* c.511C>T, Arg171*; *ACVRL1* c.1120C>T, Arg374Trp, and *ACVRL1* c.436delG).

**B) Dot plots illustration of genes for 6 endothelial markers,** contrasted to two of the genes expressed only on a proportion of BOECs. Each box represents the results of 4 x20 viable BOECs, colour coded per donor (blue/green control; red/orange HHT (*SMAD4*$^{+/-}$), with mean values for each culture indicated by a black circle.

**C) Overall comparison of transcript numbers evaluated by RNASeq and single cell (sc) qRT-PCR i)** Comparison of numbers: RNASeq N=16,807, sc qRT-PCR N=48, with circle areas proportional. **ii)** RNASeq alignments (mean in replicate, untreated control BOECs) of all 48 transcripts ranked by single cell qRT-PCR expression for the 48 genes, categorized by tertiles of expression. The single cell qRT-PCR expression scale was 0-10, p value ($p<0.0001$) calculated by Kruskal Wallis, with Dunn's post test $p<0.05$ for each pairwise comparison.

**Supplementary Fig. 3: Quality scores generated across the 150bp reads for the 4 pairs of untreated BOECs**

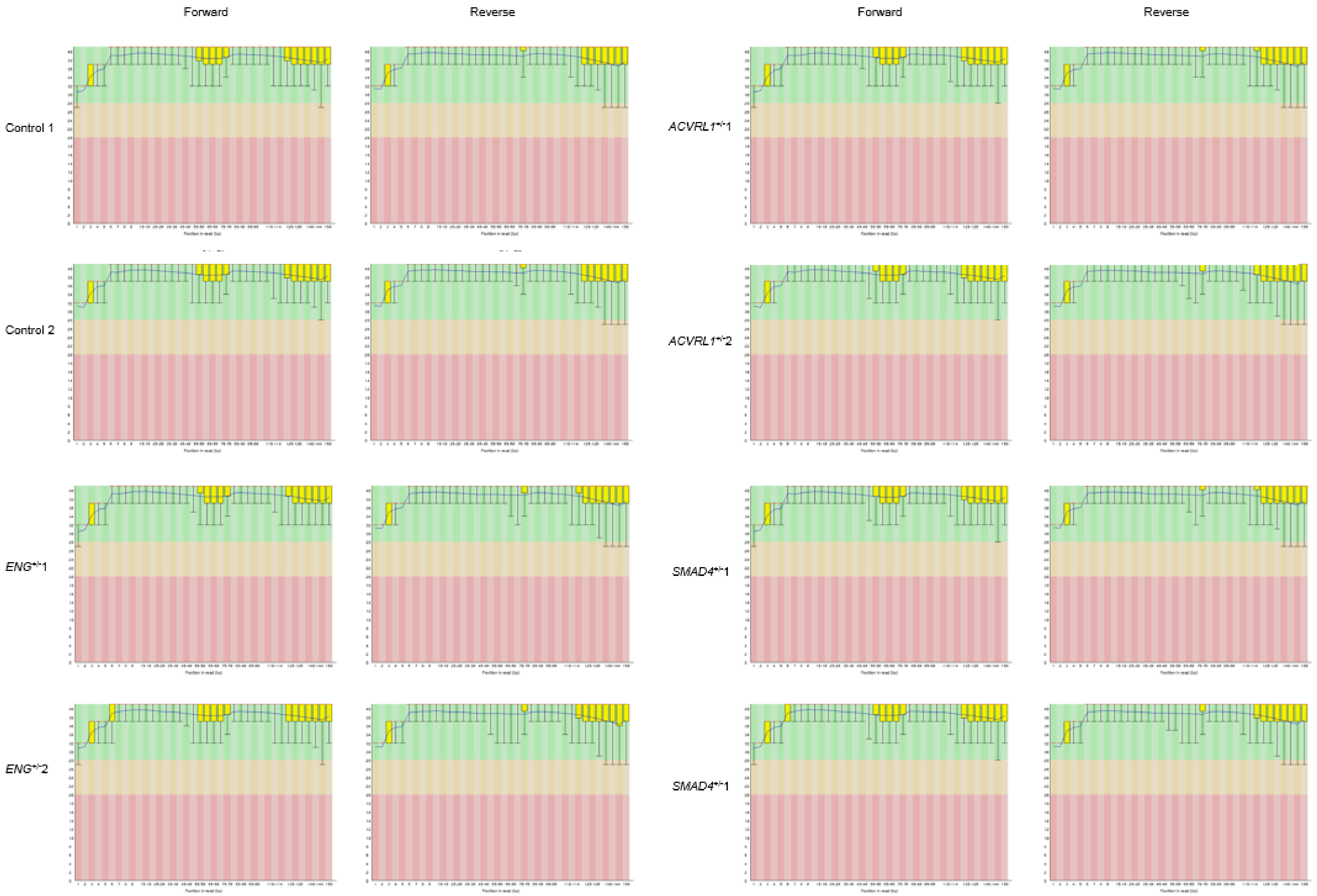

Coded data plots supplied by Genewiz

**Supplementary Fig. 4: Low Gini-coefficient reference genes for RNASeq normalizations**

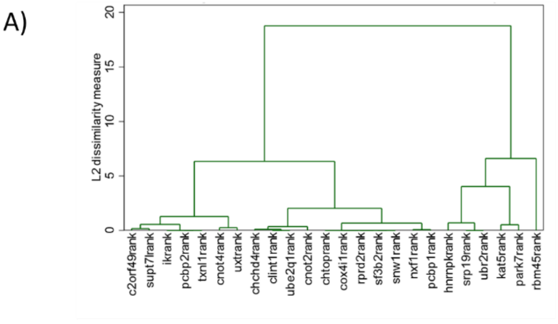

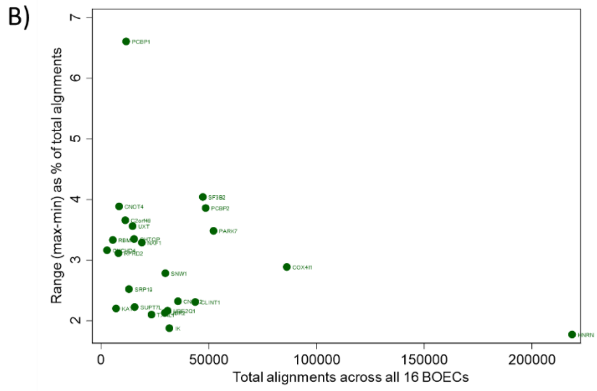

E)

| GC gene | minimum | maximum | mean |
|---|---|---|---|
| *UBR2* | 0.818112 | 1.159077 | 1.00 |
| *TXNL1* | 0.832152 | 1.168488 | 1.00 |
| *SNW1* | 0.804873 | 1.25038 | 1.00 |
| *IK* | 0.815884 | 1.116428 | 1.00 |
| *COX4I1* | 0.763953 | 1.226015 | 1.00 |
| *UBE2Q1* | 0.842423 | 1.189104 | 1.00 |
| *HNRNPK* | 0.86083 | 1.144654 | 1.00 |
| *KAT5* | 0.806968 | 1.159446 | 1.00 |

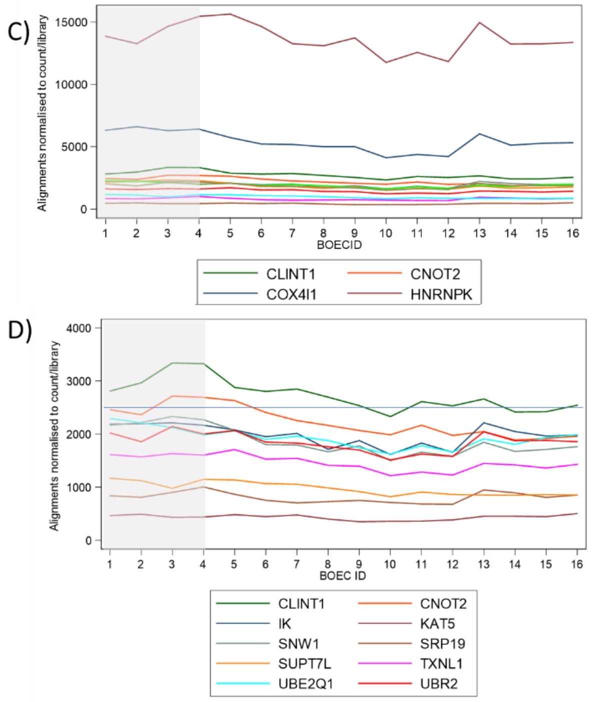

F)

| BOEC | Size factor |
|---|---|
| 1 | 1.103745 |
| 2 | 1.120637 |
| 3 | 1.11646 |
| 4 | 1.095045 |
| 5 | 1.11162 |
| 6 | 0.992159 |
| 7 | 1.002511 |
| 8 | 0.939606 |
| 9 | 0.937617 |
| 10 | 0.822878 |
| 11 | 0.884344 |
| 12 | 0.850947 |
| 13 | 1.070299 |
| 14 | 0.969812 |
| 15 | 0.98491 |
| 16 | 0.99357 |

Alignments to 24 of the 25 lowest GINI coefficient genes (GC <0.15) were identified in all 16 BOECs (See Supplementary Table 2 for details). **A)** Ward's unsupervised hierarchical linkage dendrograms for 4-way comparisons of control vs *ACVRL1*$^{+/PTC}$ vs *ENG*$^{+/PTC}$ vs *SMAD4*$^{+/PTC}$ BOEC RNASeq datasets. Note the unexpected major division of these 'most invariant' genes. **B)** Variability in alignments between BOECs expressed as the range/total alignments, by total alignments per gene. **C/D)** Alignment number per BOEC library for **C)** more highly expressed genes, and **D)** less highly expressed genes. Note these followed similar patterns with a rising pattern across the 4 control BOECs (shaded), trend down through BOECs 5-12, with 8 demonstrating a peak in BOEC13 (belonging to the 13-14 untreated replicate pair). **E)** The 8 genes (*HNRNPK, COX4I1, IK, UBR2, UBE2Q1, SNW1, TXNL1, and KAT5)* used for DeSeq2 normalization indicating the minimum and maximum ratios across the BOECs. The ratio of each gene count to the geometric mean of all read counts for that gene across all samples was used to derive the "size factor" to scale each BOEC sample as detailed in (**F**).

**Supplementary Fig. 5: Examples of raw sequence reads for nonsense variant detection**

# A

BOECs: ENSG00000139567 (*ACVRL1*) c.[1171G>T];[c.1171=]
gDNA: Chr12:51,916,157

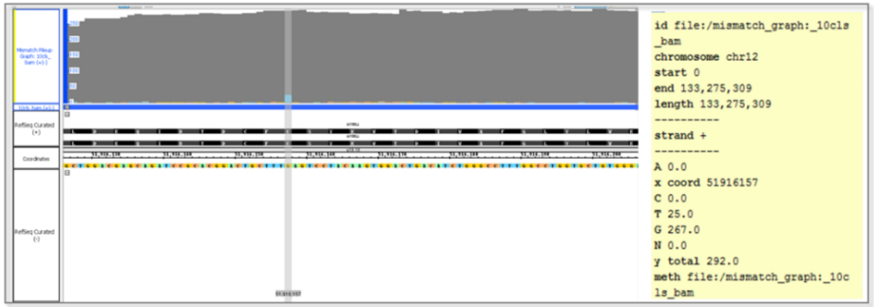

# B

BOECs: ENSG00000106991 (*ENG)* c.[277C>T];[277=]
gDNA: Chr9:127,829,769 (note *ENG* on – strand)

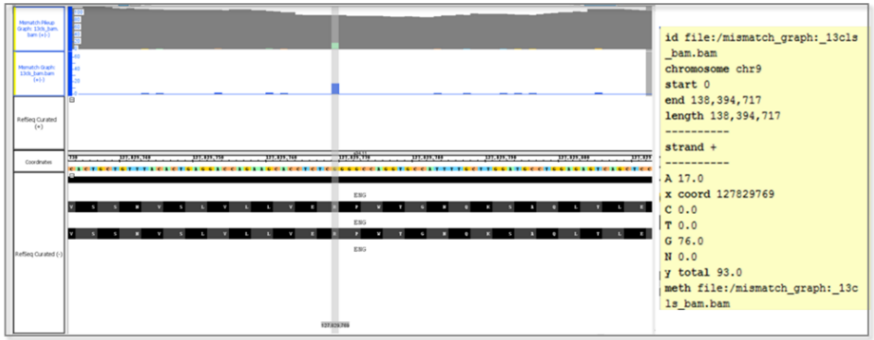

# C

BOECs: ENSG00000141646 (*SMAD4)* c.[1096C>T];[1096=]
gDNA: Chr18 :127,829,769

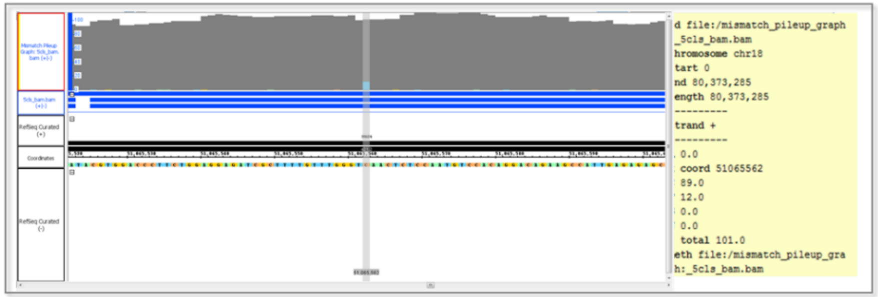

Integrated Genome Browser (IGB) 9.1.8[14] visualisations of RNASeq bam file alignments from 3 representative BOEC for **A** *ENG*$^{+/-}$; **B** *ENG*$^{+/-}$ and **C)** *SMAD4*$^{+/-}$ donors at the genomic loci corresponding to the site of their pathogenic nonsense variants. Alleles are described using Human Genome Variation Society version 20.05[47] recommendations if both alleles are described in a heterozygote, i.e. *ACVRL1* c.[1171G>T];[1171=]; *ENG* c.[277C>T];[277=]; *SMAD4* c.[1096C>T];[1096=].

**Supplementary Fig. 6: Alignments of reads to spliced transcript**

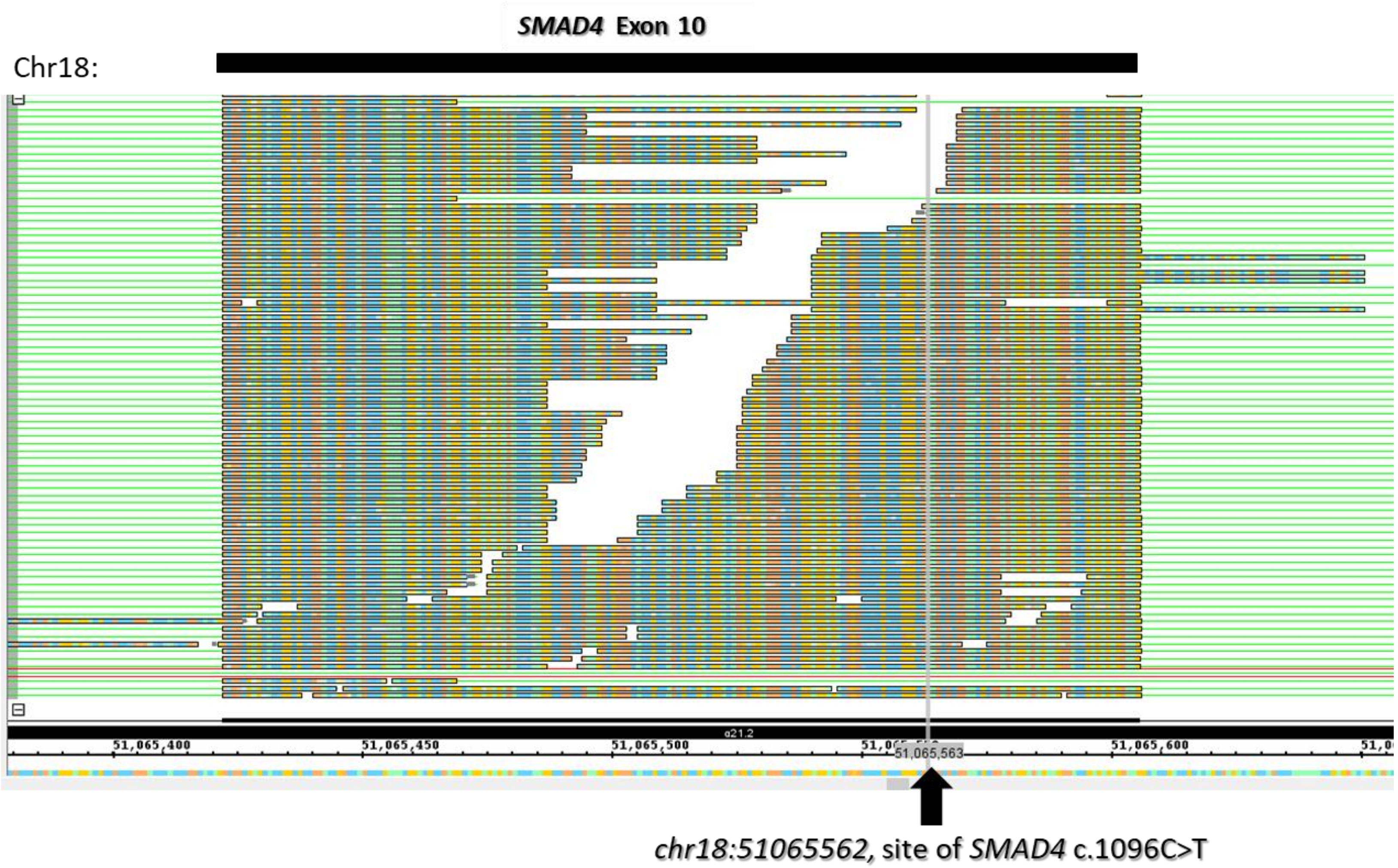

Integrated Genome Browser (IGB) 9.1.8[14] visualisations of RNASeq bam file alignments from *SMAD4*$^{+/-}$ donor BOECs at genomic loci spanning site of the pathogenic nonsense variant chr18:51065562 on *Homo sapiens* GRCh38.[9] Note that alignments sharply delineate the exon intron boundaries. The very occasional reads that also aligned to intronic sequence only aligned to wildtype sequence.

**Supplementary Fig. 7: Detailed examination of common single nucleotide variants (SNVs) in the 3' UTR of *ACVRL1***

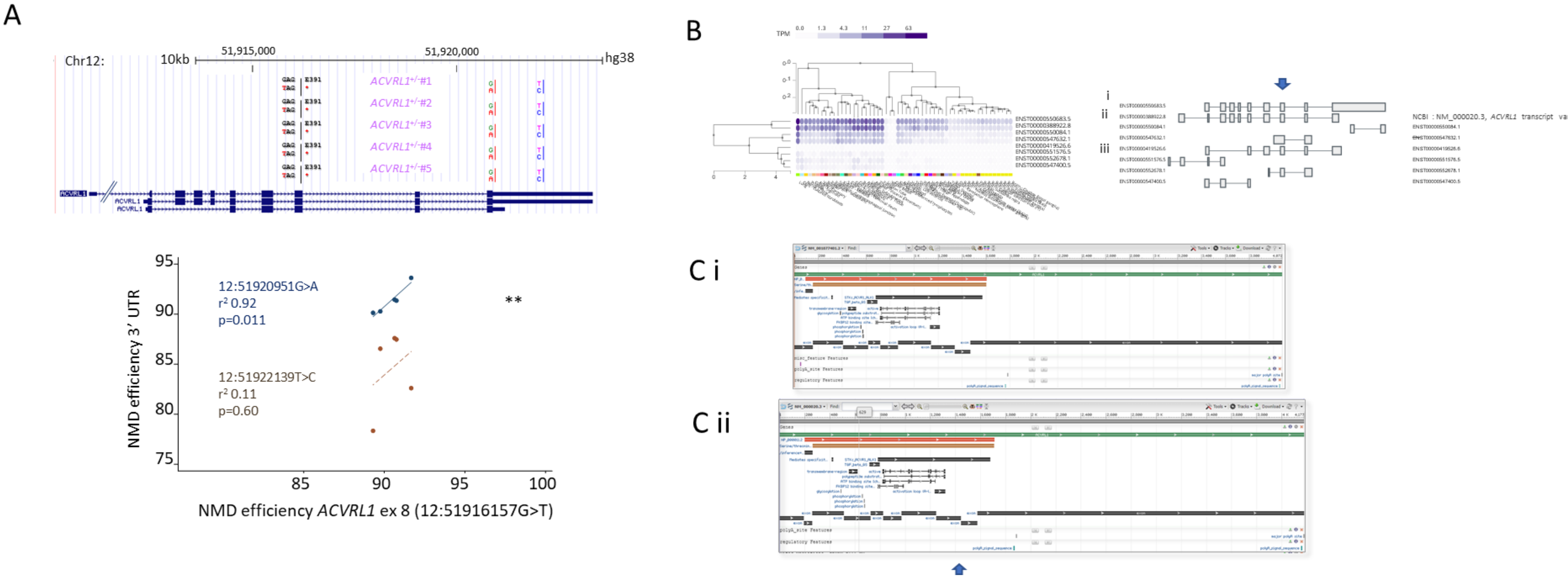

**A)** Relative genomic (GRCh38)[9] positions (custom tracks, UCSC Genome Browser[12,13]) and allele loss for two *ACVRL1* 3' UTR common variants in the five BOECs heterozygous for the *ACVRL1* exon 8 nonsense variant (chr12:51916157G>T, p.Glu391X). The proximal SNV, chr12:51920951G>A, is present in all known 3' UTRs (see **B**), whereas the more distal, chr12:51922139T>C, is sited in a region only present in transcripts with longer 3'UTRs.
**B)** *ACVRL1* splice isoform expression in GTEx[32]: **i** is the transcript corresponding to NCBI Reference Sequence: NM_001077401.2 [*ACVRL1* transcript variant 2], also shown in **Ci**. **ii** is the transcript corresponding to NCBI Reference Sequence: NM_000020.3 *ACVRL1* transcript variant 1 (shown in **Cii**, though note the differing 3'UTR lengths). Both transcripts contain all exons in which HHT-causal pathogenic variants have been identified as listed on the HHT Mutation Database, which references NM_000020.3 [*ACVRL1* transcript variant 1].[38] Note also **iii** which contains both the nonsense-containing exon 8 (arrowed) and the proximal but not distal 3' UTR variant, and not other exons which contain HHT-causal pathogenic variants.
**C)** Aligned annotations on the National Center for Biotechnology Information (NCBI)[37] for **i)** NM_001077401.2 [*ACVRL1* transcript variant 2], and **ii)** NM_000020.3 [*ACVRL1* transcript variant 1]. Again, the nonsense-containing exon 8 is arrowed.

#### Supplementary Table 4: Top 1000 differentially expressed genes normalized to total counts per library

| Rank | Ensembl Gene ID | Official gene symbol | Mean control counts* | Standard deviation control counts* | Mean HHT/ mean control | Ranking score |
|---|---|---|---|---|---|---|
| 1 | ENSG00000133110 | *POSTN* | 26654.8 | 376.5418 | 0.1722 | 10.43 |
| 2 | ENSG00000163931 | *TKT* | 14117.4 | 1012.437 | 0.3927 | 6.47 |
| 3 | ENSG00000164659 | *KIAA1324L* | 370.2549 | 22.13188 | 6.9333 | 6.00 |
| 4 | ENSG00000233913 | | 2908.972 | 57.9507 | 0.2441 | 5.72 |
| 5 | ENSG00000184009 | *ACTG1* | 148022.3 | 6295.94 | 0.5332 | 5.50 |
| 6 | ENSG00000213949 | *ITGA1* | 184.1164 | 14.8619 | 7.3185 | 5.37 |
| 7 | ENSG00000104884 | *ERCC2* | 3342.999 | 241.0485 | 0.3928 | 5.12 |
| 8 | ENSG00000168906 | *MAT2A* | 4843.216 | 427.9939 | 0.4661 | 4.63 |
| 9 | ENSG00000173597 | *SULT1B1* | 3005.8 | 144.3793 | 0.4046 | 4.50 |
| 10 | ENSG00000115602 | *IL1RL1* | 2830.229 | 235.5999 | 0.4430 | 4.45 |
| 11 | ENSG00000087086 | *FTL* | 75241.15 | 2881.234 | 0.5800 | 4.34 |
| 12 | ENSG00000001617 | *SEMA3F* | 3298.536 | 110.8267 | 0.4003 | 4.31 |
| 13 | ENSG00000138166 | *DUSP5* | 3579.96 | 161.7948 | 0.4286 | 4.31 |
| 14 | ENSG00000134668 | *SPOCD1* | 3598.399 | 245.8867 | 0.4574 | 4.31 |
| 15 | ENSG00000049449 | *RCN1* | 16609.43 | 992.8232 | 0.5470 | 4.16 |
| 16 | ENSG00000144810 | *COL8A1* | 16002.88 | 688.5164 | 0.5317 | 4.13 |
| 17 | ENSG00000161638 | *ITGA5* | 37474.91 | 1080.997 | 0.5684 | 3.95 |
| 18 | ENSG00000184470 | *TXNRD2* | 3766.465 | 107.0786 | 0.4328 | 3.91 |
| 19 | ENSG00000162772 | *ATF3* | 2230.2 | 92.72086 | 0.4246 | 3.88 |
| 20 | ENSG00000163739 | *CXCL1* | 3707.577 | 221.2235 | 0.4916 | 3.83 |
| 21 | ENSG00000125148 | *MT2A* | 15971.25 | 1438.538 | 0.5949 | 3.78 |
| 22 | ENSG00000104341 | *LAPTM4B* | 2639.014 | 210.1877 | 0.4943 | 3.77 |
| 23 | ENSG00000125810 | *CD93* | 39461.36 | 2643.33 | 0.6211 | 3.75 |
| 24 | ENSG00000167601 | *AXL* | 7852.663 | 267.4762 | 0.5127 | 3.73 |
| 25 | ENSG00000120708 | *TGFBI* | 708.2304 | 50.38221 | 2.5889 | 3.73 |
| 26 | ENSG00000279086 | | 273.3106 | 14.72698 | 0.2533 | 3.69 |
| 27 | ENSG00000142192 | *APP* | 66301.63 | 3400.013 | 0.6431 | 3.59 |
| 28 | ENSG00000173269 | *MMRN2* | 11681.27 | 390.0654 | 0.5486 | 3.58 |
| 29 | ENSG00000114698 | *PLSCR4* | 4473.404 | 139.8934 | 2.0421 | 3.53 |
| 30 | ENSG00000103257 | *SLC7A5* | 1800.436 | 118.0204 | 0.4816 | 3.49 |
| 31 | ENSG00000072110 | *ACTN1* | 17081.63 | 588.7491 | 0.5860 | 3.41 |
| 32 | ENSG00000065923 | *SLC9A7* | 1603.062 | 99.72222 | 0.4843 | 3.34 |
| 33 | ENSG00000070669 | *ASNS* | 731.8263 | 47.27482 | 0.4256 | 3.29 |
| 34 | ENSG00000163399 | *ATP1A1* | 10335.6 | 773.7389 | 0.6116 | 3.27 |
| 35 | ENSG00000173334 | *TRIB1* | 1544.601 | 43.47824 | 0.4212 | 3.26 |
| 36 | ENSG00000140873 | *ADAMTS18* | 7838.943 | 437.9732 | 0.5906 | 3.20 |
| 37 | ENSG00000141101 | *NOB1* | 1066.489 | 88.44758 | 0.4911 | 3.19 |
| 38 | ENSG00000143384 | *MCL1* | 23108.16 | 890.1595 | 0.6268 | 3.17 |
| 39 | ENSG00000239264 | *TXNDC5* | 9823.581 | 425.9681 | 0.5928 | 3.17 |
| 40 | ENSG00000135069 | *PSAT1* | 904.5049 | 22.11096 | 0.3610 | 3.15 |
| 41 | ENSG00000096384 | *HSP90AB1* | 30079.94 | 1956.04 | 0.6605 | 3.14 |
| 42 | ENSG00000166197 | *NOLC1* | 2504.548 | 187.4885 | 0.5515 | 3.11 |
| 43 | ENSG00000160145 | *KALRN* | 757.8326 | 19.76579 | 2.7363 | 3.00 |
| 44 | ENSG00000134333 | *LDHA* | 9153.639 | 700.1649 | 0.6323 | 3.00 |
| 45 | ENSG00000106688 | *SLC1A1* | 1049.711 | 37.6954 | 0.4391 | 2.99 |
| 46 | ENSG00000186716 | *BCR* | 6474.071 | 558.3758 | 0.6251 | 2.97 |
| 47 | ENSG00000184226 | *PCDH9* | 151.2016 | 10.64231 | 3.4780 | 2.95 |
| 48 | ENSG00000173705 | *SUSD5* | 1699.266 | 133.1351 | 0.5484 | 2.94 |
| 49 | ENSG00000153162 | *BMP6* | 8042.333 | 245.738 | 0.5871 | 2.93 |
| 50 | ENSG00000112208 | *BAG2* | 919.2485 | 66.46931 | 0.4974 | 2.93 |
| 51 | ENSG00000065911 | *MTHFD2* | 2953.434 | 130.3002 | 0.5496 | 2.92 |
| 52 | ENSG00000137507 | *LRRC32* | 4956.705 | 167.7034 | 0.5670 | 2.91 |
| 53 | ENSG00000186594 | *MIR22HG* | 3209.286 | 81.41171 | 0.5176 | 2.90 |
| 54 | ENSG00000102265 | *TIMP1* | 3392.385 | 148.1483 | 0.5616 | 2.88 |
| 55 | ENSG00000112837 | *TBX18* | 1348.941 | 30.5303 | 0.4303 | 2.88 |

| Rank | Ensembl Gene ID | Official gene symbol | Mean control counts* | Standard deviation control counts* | Mean HHT/ mean control | Ranking score |
|---|---|---|---|---|---|---|
| 56 | ENSG00000117298 | *ECE1* | 14080.63 | 950.6048 | 0.6618 | 2.83 |
| 57 | ENSG00000171793 | *CTPS1* | 1376.305 | 53.79674 | 0.4924 | 2.82 |
| 58 | ENSG00000198431 | *TXNRD1* | 41566.23 | 1358.97 | 0.6763 | 2.82 |
| 59 | ENSG00000197019 | *SERTAD1* | 4028.914 | 154.0008 | 0.5730 | 2.80 |
| 60 | ENSG00000075624 | *ACTB* | 134746.1 | 2804.552 | 0.7026 | 2.80 |
| 61 | ENSG00000141582 | *CBX4* | 1022.776 | 68.77336 | 0.5182 | 2.78 |
| 62 | ENSG00000177606 | *JUN* | 6662.5 | 283.3173 | 0.6114 | 2.78 |
| 63 | ENSG00000011422 | *PLAUR* | 1824.779 | 117.0301 | 0.5583 | 2.78 |
| 64 | ENSG00000149925 | *ALDOA* | 15210.22 | 205.9188 | 0.5941 | 2.77 |
| 65 | ENSG00000139514 | *SLC7A1* | 2238.689 | 139.7995 | 0.5727 | 2.75 |
| 66 | ENSG00000152952 | *PLOD2* | 14923.68 | 746.9146 | 0.6598 | 2.75 |
| 67 | ENSG00000149591 | *TAGLN* | 2131.969 | 194.9124 | 0.5937 | 2.75 |
| 68 | ENSG00000176978 | *DPP7* | 2208.846 | 145.1164 | 0.5758 | 2.75 |
| 69 | ENSG00000166337 | *TAF10* | 1850.606 | 75.12823 | 0.5295 | 2.75 |
| 70 | ENSG00000146411 | *SLC2A12* | 51.24069 | 3.294562 | 9.8981 | 2.73 |
| 71 | ENSG00000171992 | *SYNPO* | 4105.535 | 131.223 | 0.5721 | 2.72 |
| 72 | ENSG00000092969 | *TGFB2* | 105.6191 | 6.266095 | 4.4017 | 2.72 |
| 73 | ENSG00000175334 | *BANF1* | 1064.613 | 102.6296 | 0.5561 | 2.72 |
| 74 | ENSG00000049323 | *LTBP1* | 4731.561 | 49.82499 | 0.5009 | 2.70 |
| 75 | ENSG00000125505 | *MBOAT7* | 2866.489 | 181.7086 | 0.5951 | 2.70 |
| 76 | ENSG00000163874 | *ZC3H12A* | 962.3531 | 26.55453 | 0.4404 | 2.69 |
| 77 | ENSG00000134874 | *DZIP1* | 1354.368 | 33.34578 | 0.4652 | 2.68 |
| 78 | ENSG00000100504 | *PYGL* | 3780.109 | 112.8407 | 0.5670 | 2.68 |
| 79 | ENSG00000104368 | *PLAT* | 2622.031 | 172.6531 | 0.5953 | 2.67 |
| 80 | ENSG00000102802 | *MEDAG* | 294.6419 | 8.425163 | 3.4846 | 2.66 |
| 81 | ENSG00000048052 | *HDAC9* | 489.8862 | 30.60664 | 2.1740 | 2.66 |
| 82 | ENSG00000127481 | *UBR4* | 8293.803 | 617.7647 | 0.6615 | 2.66 |
| 83 | ENSG00000171877 | *FRMD5* | 895.563 | 33.31559 | 0.4699 | 2.65 |
| 84 | ENSG00000124875 | *CXCL6* | 214.4949 | 15.1303 | 2.6246 | 2.62 |
| 85 | ENSG00000073008 | *PVR* | 7823.824 | 566.2089 | 0.6616 | 2.62 |
| 86 | ENSG00000139278 | *GLIPR1* | 4246.061 | 57.68877 | 0.5245 | 2.62 |
| 87 | ENSG00000128872 | *TMOD2* | 179.4502 | 0.1549022 | 4.0613 | 2.61 |
| 88 | ENSG00000142657 | *PGD* | 7018.457 | 396.6919 | 0.6465 | 2.61 |
| 89 | ENSG00000179222 | *MAGED1* | 5017.459 | 302.626 | 0.6361 | 2.58 |
| 90 | ENSG00000104774 | *MAN2B1* | 2162.283 | 178.6989 | 0.6076 | 2.58 |
| 91 | ENSG00000065268 | *WDR18* | 728.0197 | 69.95076 | 0.5448 | 2.58 |
| 92 | ENSG00000103202 | *NME4* | 2343.062 | 144.7065 | 0.5955 | 2.58 |
| 93 | ENSG00000179431 | *FJX1* | 2812.162 | 238.897 | 0.6250 | 2.57 |
| 94 | ENSG00000198242 | *RPL23A* | 23795.12 | 537.3467 | 0.6646 | 2.57 |
| 95 | ENSG00000120742 | *SERP1* | 6532.447 | 498.2672 | 0.6622 | 2.56 |
| 96 | ENSG00000184465 | *WDR27* | 780.239 | 63.83123 | 0.5419 | 2.55 |
| 97 | ENSG00000004478 | *FKBP4* | 3016.582 | 97.11448 | 0.5766 | 2.52 |
| 98 | ENSG00000158615 | *PPP1R15B* | 4762.588 | 177.0677 | 0.6147 | 2.52 |
| 99 | ENSG00000104763 | *ASAH1* | 1137.552 | 50.95977 | 1.8959 | 2.51 |
| 100 | ENSG00000070087 | *PFN2* | 1498.242 | 84.54174 | 0.5674 | 2.51 |
| 101 | ENSG00000038382 | *TRIO* | 11621.45 | 618.2462 | 0.6771 | 2.51 |
| 102 | ENSG00000165072 | *MAMDC2* | 388.3026 | 25.83875 | 2.1578 | 2.50 |
| 103 | ENSG00000175592 | *FOSL1* | 1544.329 | 38.27771 | 0.5045 | 2.49 |
| 104 | ENSG00000162733 | *DDR2* | 721.9843 | 16.52068 | 2.4217 | 2.48 |
| 105 | ENSG00000166147 | *FBN1* | 12772.14 | 645.931 | 0.6816 | 2.48 |
| 106 | ENSG00000154134 | *ROBO3* | 508.4276 | 24.55999 | 0.4618 | 2.47 |
| 107 | ENSG00000165629 | *ATP5C1* | 4972.212 | 281.8607 | 0.6457 | 2.47 |
| 108 | ENSG00000141551 | *CSNK1D* | 2079.217 | 179.4311 | 0.6228 | 2.46 |
| 109 | ENSG00000176170 | *SPHK1* | 6391.061 | 141.242 | 0.6094 | 2.45 |
| 110 | ENSG00000079257 | *LXN* | 3331.902 | 24.73179 | 0.4679 | 2.44 |
| 111 | ENSG00000152767 | *FARP1* | 1505.523 | 110.2712 | 0.5957 | 2.44 |
| 112 | ENSG00000103966 | *EHD4* | 9949.068 | 444.9171 | 0.6715 | 2.43 |
| 113 | ENSG00000158710 | *TAGLN2* | 7922.88 | 410.2708 | 0.6681 | 2.43 |

| Rank | Ensembl Gene ID | Official gene symbol | Mean control counts* | Standard deviation control counts* | Mean HHT/ mean control | Ranking score | Rank | Ensembl Gene ID | Official gene symbol | Mean control counts* | Standard deviation control counts* | Mean HHT/ mean control | Ranking score |
|---|---|---|---|---|---|---|---|---|---|---|---|---|---|
| 114 | ENSG00000176788 | *BASP1* | 2250.865 | 204.72 | 0.6346 | 2.42 | 143 | ENSG00000107185 | *RGP1* | 1648.134 | 117.3512 | 0.6201 | 2.28 |
| 115 | ENSG00000253200 | | 149.7943 | 0.0058224 | 0.6251 | 2.42 | 144 | ENSG00000143321 | *HDGF* | 6402.871 | 168.0316 | 0.6425 | 2.27 |
| 116 | ENSG00000113140 | *SPARC* | 58218.69 | 3613.068 | 0.7454 | 2.41 | 145 | ENSG00000106853 | *PTGR1* | 2996.71 | 139.9418 | 0.6322 | 2.27 |
| 117 | ENSG00000172889 | *EGFL7* | 10293.81 | 665.8668 | 0.6912 | 2.40 | 146 | ENSG00000157514 | *TSC22D3* | 744.2245 | 23.57796 | 0.4884 | 2.26 |
| 118 | ENSG00000104823 | *ECH1* | 1066.327 | 71.38348 | 0.5700 | 2.40 | 147 | ENSG00000156976 | *EIF4A2* | 15299.16 | 799.5239 | 0.7127 | 2.26 |
| 119 | ENSG00000155660 | *PDIA4* | 10956.37 | 240.2563 | 0.6459 | 2.40 | 148 | ENSG00000090863 | *GLG1* | 7325.598 | 504.1657 | 0.6953 | 2.26 |
| 120 | ENSG00000073578 | *SDHA* | 4140.298 | 142.4122 | 0.6192 | 2.38 | 149 | ENSG00000184216 | *IRAK1* | 3653.106 | 187.4225 | 0.6498 | 2.26 |
| 121 | ENSG00000140406 | *MESDC1* | 1772.816 | 168.3049 | 0.6303 | 2.37 | 150 | ENSG00000091490 | *SEL1L3* | 7879.057 | 130.1147 | 0.6297 | 2.25 |
| 122 | ENSG00000171867 | *PRNP* | 5791.007 | 349.9803 | 0.6678 | 2.36 | 151 | ENSG00000169499 | *PLEKHA2* | 267.6008 | 19.28693 | 0.4676 | 2.25 |
| 123 | ENSG00000130479 | *MAP1S* | 921.2777 | 41.62237 | 0.5304 | 2.36 | 152 | ENSG00000167552 | *TUBA1A* | 10676.19 | 575.4505 | 1.4243 | 2.25 |
| 124 | ENSG00000090273 | *NUDC* | 1936.032 | 187.3818 | 0.6368 | 2.36 | 153 | ENSG00000071127 | *WDR1* | 20951.19 | 433.8622 | 0.6907 | 2.25 |
| 125 | ENSG00000065978 | *YBX1* | 19262.2 | 642.1424 | 0.6940 | 2.36 | 154 | ENSG00000106617 | *PRKAG2* | 2296.38 | 180.7015 | 0.6496 | 2.24 |
| 126 | ENSG00000065054 | *SLC9A3R2* | 1083.621 | 39.72164 | 0.5273 | 2.36 | 155 | ENSG00000162413 | *KLHL21* | 2451.953 | 111.3651 | 0.6218 | 2.24 |
| 127 | ENSG00000078269 | *SYNJ2* | 5890.674 | 364.6642 | 0.6713 | 2.35 | 156 | ENSG00000163171 | *CDC42EP3* | 1617.206 | 92.3455 | 0.6100 | 2.24 |
| 128 | ENSG00000142089 | *IFITM3* | 6350.348 | 230.486 | 0.6496 | 2.35 | 157 | ENSG00000111057 | *KRT18* | 5545.124 | 288.4522 | 0.6740 | 2.23 |
| 129 | ENSG00000160796 | *NBEAL2* | 1541.178 | 52.55493 | 0.5550 | 2.33 | 158 | ENSG00000187678 | *SPRY4* | 2228.122 | 85.57272 | 0.6053 | 2.23 |
| 130 | ENSG00000196562 | *SULF2* | 9955.438 | 409.2527 | 0.6787 | 2.33 | 159 | ENSG00000145990 | *GFOD1* | 1609.337 | 74.01178 | 0.5957 | 2.23 |
| 131 | ENSG00000225178 | *RPSAP58* | 171.4734 | 6.802753 | 3.3528 | 2.32 | 160 | ENSG00000010278 | *CD9* | 14536.31 | 235.5648 | 0.6664 | 2.22 |
| 132 | ENSG00000164687 | *FABP5* | 1286.651 | 18.27845 | 2.2130 | 2.31 | 161 | ENSG00000003249 | *DBNDD1* | 980.9619 | 25.49113 | 0.5048 | 2.21 |
| 133 | ENSG00000147065 | *MSN* | 41256.07 | 402.6525 | 0.6811 | 2.30 | 162 | ENSG00000136205 | *TNS3* | 697.9464 | 26.01758 | 1.9715 | 2.21 |
| 134 | ENSG00000150991 | *UBC* | 41535.11 | 1555.503 | 0.7310 | 2.30 | 163 | ENSG00000204946 | *ZNF783* | 318.4644 | 23.82387 | 0.4978 | 2.21 |
| 135 | ENSG00000162244 | *RPL29* | 26601.24 | 389.4792 | 0.6800 | 2.30 | 164 | ENSG00000148840 | *PPRC1* | 1131.405 | 96.07523 | 0.6181 | 2.20 |
| 136 | ENSG00000107815 | *C10orf2* | 477.8201 | 30.37806 | 0.5101 | 2.30 | 165 | ENSG00000087245 | *MMP2* | 6316.992 | 53.7856 | 0.5766 | 2.19 |
| 137 | ENSG00000128849 | *CGNL1* | 1943.064 | 27.56775 | 0.5007 | 2.29 | 166 | ENSG00000011243 | *AKAP8L* | 1281.962 | 70.4923 | 0.5975 | 2.19 |
| 138 | ENSG00000091136 | *LAMB1* | 35824.33 | 2266.75 | 0.7434 | 2.29 | 167 | ENSG00000181192 | *DHTKD1* | 1012.871 | 0.0059666 | 0.6522 | 2.19 |
| 139 | ENSG00000113070 | *HBEGF* | 2410.066 | 83.33243 | 0.5957 | 2.29 | 168 | ENSG00000148700 | *ADD3* | 1920.383 | 191.101 | 1.5164 | 2.19 |
| 140 | ENSG00000100219 | *XBP1* | 4186.955 | 132.5241 | 0.6261 | 2.29 | 169 | ENSG00000169991 | *IFFO2* | 3329.961 | 85.11951 | 0.6114 | 2.19 |
| 141 | ENSG00000255717 | *SNHG1* | 1927.301 | 114.707 | 0.6185 | 2.28 | 170 | ENSG00000185420 | *SMYD3* | 640.7399 | 36.3079 | 0.5444 | 2.18 |
| 142 | ENSG00000064932 | *SBNO2* | 2902.997 | 148.3167 | 0.6341 | 2.28 | 171 | ENSG00000158805 | *ZNF276* | 1230.849 | 71.94041 | 0.6002 | 2.18 |

| Rank | Ensembl Gene ID | Official gene symbol | Mean control counts* | Standard deviation control counts* | Mean HHT/ mean control | Ranking score | Rank | Ensembl Gene ID | Official gene symbol | Mean control counts* | Standard deviation control counts* | Mean HHT/ mean control | Ranking score |
|---|---|---|---|---|---|---|---|---|---|---|---|---|---|
| 172 | ENSG00000178952 | *TUFM* | 5183 | 160.7432 | 0.6519 | 2.17 | 201 | ENSG00000165689 | *SDCCAG3* | 952.5112 | 57.30867 | 0.5963 | 2.09 |
| 173 | ENSG00000111674 | *ENO2* | 286.1826 | 17.50631 | 0.4691 | 2.17 | 202 | ENSG00000152689 | *RASGRP3* | 1097.612 | 75.17531 | 1.6229 | 2.09 |
| 174 | ENSG00000125538 | *IL1B* | 72.82552 | 6.440045 | 0.3128 | 2.16 | 203 | ENSG00000196526 | *AFAP1* | 6356.872 | 123.2673 | 0.6497 | 2.08 |
| 175 | ENSG00000101265 | *RASSF2* | 2852.169 | 207.5714 | 0.6666 | 2.16 | 204 | ENSG00000232956 | *SNHG15* | 712.9213 | 14.92724 | 0.4645 | 2.07 |
| 176 | ENSG00000163110 | *PDLIM5* | 10618.52 | 454.9831 | 0.7026 | 2.16 | 205 | ENSG00000140743 | *CDR2* | 1768.296 | 97.47726 | 0.6361 | 2.07 |
| 177 | ENSG00000188735 | *TMEM120B* | 524.3518 | 45.51512 | 0.5679 | 2.16 | 206 | ENSG00000073536 | *NLE1* | 607.2594 | 48.92067 | 0.5897 | 2.05 |
| 178 | ENSG00000204351 | *SKIV2L* | 1213.83 | 94.79317 | 0.6222 | 2.16 | 207 | ENSG00000102755 | *FLT1* | 4541.166 | 362.5359 | 0.7060 | 2.05 |
| 179 | ENSG00000171490 | *RSL1D1* | 4892.903 | 154.5368 | 0.6518 | 2.16 | 208 | ENSG00000166557 | *TMED3* | 3811.165 | 57.89219 | 0.6038 | 2.05 |
| 180 | ENSG00000143761 | *ARF1* | 9069.467 | 304.7447 | 0.6859 | 2.16 | 209 | ENSG00000147955 | *SIGMAR1* | 893.0829 | 36.82299 | 0.5671 | 2.05 |
| 181 | ENSG00000198053 | *SIRPA* | 776.0149 | 42.42491 | 1.7739 | 2.15 | 210 | ENSG00000188522 | *FAM83G* | 2271.653 | 64.36019 | 0.6120 | 2.05 |
| 182 | ENSG00000235770 | *LINC00607* | 2566.239 | 13.78154 | 0.4412 | 2.15 | 211 | ENSG00000166886 | *NAB2* | 3285.287 | 279.6956 | 0.6958 | 2.04 |
| 183 | ENSG00000138326 | *RPS24* | 31638.07 | 2313.271 | 0.7583 | 2.14 | 212 | ENSG00000142534 | *RPS11* | 35681.52 | 1613.787 | 0.7583 | 2.04 |
| 184 | ENSG00000135048 | *TMEM2* | 3820.52 | 131.6461 | 0.6446 | 2.14 | 213 | ENSG00000163071 | *SPATA18* | 211.0362 | 8.797935 | 2.5491 | 2.03 |
| 185 | ENSG00000121966 | *CXCR4* | 91.54197 | 5.999419 | 3.3061 | 2.14 | 214 | ENSG00000165959 | *CLMN* | 636.7448 | 39.07572 | 0.5744 | 2.03 |
| 186 | ENSG00000116273 | *PHF13* | 812.4153 | 64.44077 | 0.5983 | 2.14 | 215 | ENSG00000214530 | *STARD10* | 691.2667 | 42.3531 | 0.5816 | 2.03 |
| 187 | ENSG00000111252 | *SH2B3* | 7390.744 | 366.1348 | 0.6964 | 2.14 | 216 | ENSG00000110799 | *VWF* | 83862.52 | 2198.159 | 0.7690 | 2.02 |
| 188 | ENSG00000110104 | *CCDC86* | 794.958 | 25.72273 | 0.5199 | 2.12 | 217 | ENSG00000142910 | *TINAGL1* | 11003.57 | 182.1477 | 0.6787 | 2.02 |
| 189 | ENSG00000134470 | *IL15RA* | 631.1135 | 28.30589 | 0.5298 | 2.12 | 218 | ENSG00000087460 | *GNAS* | 14953.11 | 245.5933 | 0.6944 | 2.01 |
| 190 | ENSG00000168209 | *DDIT4* | 4328.873 | 274.1436 | 0.6854 | 2.12 | 219 | ENSG00000124216 | *SNAI1* | 2782.251 | 52.7066 | 0.6028 | 2.01 |
| 191 | ENSG00000081665 | *ZNF506* | 666.598 | 46.37481 | 0.5755 | 2.12 | 220 | ENSG00000055483 | *USP36* | 964.5416 | 45.28546 | 0.5911 | 2.00 |
| 192 | ENSG00000177469 | *PTRF* | 35852.82 | 1358.892 | 0.7456 | 2.12 | 221 | ENSG00000160209 | *PDXK* | 1675.614 | 77.2387 | 0.6310 | 2.00 |
| 193 | ENSG00000048405 | *ZNF800* | 1718.21 | 148.724 | 0.6551 | 2.12 | 222 | ENSG00000247134 | | 327.1796 | 5.886854 | 0.3234 | 2.00 |
| 194 | ENSG00000114354 | *TFG* | 2602.996 | 232.717 | 0.6784 | 2.11 | 223 | ENSG00000149781 | *FERMT3* | 1417.065 | 111.439 | 0.6544 | 2.00 |
| 195 | ENSG00000182054 | *IDH2* | 2631.956 | 53.21767 | 0.5876 | 2.11 | 224 | ENSG00000067208 | *EVI5* | 1704.574 | 109.7768 | 0.6536 | 2.00 |
| 196 | ENSG00000146701 | *MDH2* | 3358.194 | 40.86913 | 0.5659 | 2.11 | 225 | ENSG00000116285 | *ERRFI1* | 3701.591 | 57.12444 | 0.6103 | 2.00 |
| 197 | ENSG00000182492 | *BGN* | 1715.771 | 140.9505 | 0.6531 | 2.11 | 226 | ENSG00000141696 | *P3H4* | 1428.976 | 63.92187 | 0.6193 | 1.99 |
| 198 | ENSG00000087365 | *SF3B2* | 4059 | 156.5113 | 0.6595 | 2.10 | 227 | ENSG00000070610 | *GBA2* | 3518.521 | 250.7714 | 0.6974 | 1.99 |
| 199 | ENSG00000185624 | *P4HB* | 18488.08 | 375.1164 | 0.7016 | 2.10 | 228 | ENSG00000186815 | *TPCN1* | 912.0665 | 24.38683 | 0.5368 | 1.99 |
| 200 | ENSG00000140403 | *DNAJA4* | 520.1459 | 41.13214 | 0.5686 | 2.10 | 229 | ENSG00000179262 | *RAD23A* | 1444.81 | 37.31947 | 0.5779 | 1.98 |

| Rank | Ensembl Gene ID | Official gene symbol | Mean control counts* | Standard deviation control counts* | Mean HHT/ mean control | Ranking score | Rank | Ensembl Gene ID | Official gene symbol | Mean control counts* | Standard deviation control counts* | Mean HHT/ mean control | Ranking score |
|---|---|---|---|---|---|---|---|---|---|---|---|---|---|
| 230 | ENSG00000090621 | *PABPC4* | 5035.919 | 30.42536 | 0.5593 | 1.98 | 259 | ENSG00000157020 | *SEC13* | 4882.846 | 101.0305 | 0.6625 | 1.90 |
| 231 | ENSG00000154237 | *LRRK1* | 1169.264 | 59.82685 | 0.6158 | 1.98 | 260 | ENSG00000149485 | *FADS1* | 1599.98 | 53.57362 | 0.6205 | 1.90 |
| 232 | ENSG00000124571 | *XPO5* | 1835.413 | 140.1058 | 0.6698 | 1.98 | 261 | ENSG00000167642 | *SPINT2* | 100.109 | 8.294341 | 2.4521 | 1.90 |
| 233 | ENSG00000119689 | *DLST* | 2230.012 | 72.81274 | 0.6302 | 1.98 | 262 | ENSG00000111817 | *DSE* | 909.5348 | 75.52199 | 0.6450 | 1.90 |
| 234 | ENSG00000174748 | *RPL15* | 30523.32 | 791.4928 | 0.7434 | 1.98 | 263 | ENSG00000147454 | *SLC25A37* | 1560.565 | 22.33084 | 0.5432 | 1.90 |
| 235 | ENSG00000179091 | *CYC1* | 2591.224 | 108.0159 | 0.6553 | 1.98 | 264 | ENSG00000048740 | *CELF2* | 904.8282 | 56.23916 | 1.5978 | 1.89 |
| 236 | ENSG00000073050 | *XRCC1* | 628.1912 | 38.2028 | 0.5812 | 1.98 | 265 | ENSG00000113790 | *EHHADH* | 283.7429 | 25.27973 | 0.5574 | 1.89 |
| 237 | ENSG00000102119 | *EMD* | 650.4202 | 49.99793 | 0.6035 | 1.98 | 266 | ENSG00000134352 | *IL6ST* | 39438.23 | 2485.243 | 1.2728 | 1.89 |
| 238 | ENSG00000106211 | *HSPB1* | 9887.668 | 237.0721 | 0.6970 | 1.97 | 267 | ENSG00000164176 | *EDIL3* | 339.0056 | 22.06136 | 1.8341 | 1.88 |
| 239 | ENSG00000115677 | *HDLBP* | 26797.1 | 371.7135 | 0.7171 | 1.97 | 268 | ENSG00000204381 | *LAYN* | 126.2254 | 5.94558 | 2.8635 | 1.88 |
| 240 | ENSG00000143641 | *GALNT2* | 2999.437 | 268.6622 | 0.7041 | 1.96 | 269 | ENSG00000154556 | *SORBS2* | 934.3066 | 15.71181 | 1.9745 | 1.87 |
| 241 | ENSG00000084774 | *CAD* | 1221.19 | 75.73892 | 0.6355 | 1.96 | 270 | ENSG00000168502 | *MTCL1* | 2601.488 | 73.45205 | 0.6467 | 1.87 |
| 242 | ENSG00000128340 | *RAC2* | 2683.61 | 199.4112 | 0.6922 | 1.95 | 271 | ENSG00000240694 | *PNMA2* | 378.8243 | 33.47883 | 0.5869 | 1.87 |
| 243 | ENSG00000154640 | *BTG3* | 742.4987 | 59.5619 | 0.6220 | 1.94 | 272 | ENSG00000134107 | *BHLHE40* | 6157.733 | 44.71768 | 0.6113 | 1.87 |
| 244 | ENSG00000165283 | *STOML2* | 3424.137 | 55.27172 | 0.6170 | 1.94 | 273 | ENSG00000117523 | *PRRC2C* | 5510.431 | 393.7156 | 0.7315 | 1.87 |
| 245 | ENSG00000149564 | *ESAM* | 4476.298 | 112.5226 | 0.6635 | 1.94 | 274 | ENSG00000143153 | *ATP1B1* | 2101.762 | 65.40706 | 0.6397 | 1.87 |
| 246 | ENSG00000089597 | *GANAB* | 6977.975 | 262.2803 | 0.7064 | 1.94 | 275 | ENSG00000205336 | *ADGRG1* | 1980.043 | 110.73 | 0.6731 | 1.86 |
| 247 | ENSG00000126216 | *TUBGCP3* | 863.8962 | 33.42657 | 0.5773 | 1.93 | 276 | ENSG00000117036 | *ETV3* | 2304.038 | 72.22549 | 0.6471 | 1.86 |
| 248 | ENSG00000163938 | *GNL3* | 3490.432 | 204.0323 | 0.6959 | 1.93 | 277 | ENSG00000006194 | *ZNF263* | 1687.893 | 46.90363 | 0.6165 | 1.86 |
| 249 | ENSG00000103495 | *MAZ* | 1184.65 | 67.12859 | 0.6324 | 1.93 | 278 | ENSG00000074800 | *ENO1* | 41006.45 | 637.8235 | 0.7501 | 1.86 |
| 250 | ENSG00000176485 | *PLA2G16* | 1098.712 | 65.2022 | 0.6312 | 1.92 | 279 | ENSG00000161940 | *BCL6B* | 5852.046 | 317.0873 | 0.7244 | 1.86 |
| 251 | ENSG00000165006 | *UBAP1* | 1315.387 | 96.71705 | 0.6580 | 1.91 | 280 | ENSG00000101384 | *JAG1* | 5472.49 | 215.5134 | 1.4123 | 1.85 |
| 252 | ENSG00000157617 | *C2CD2* | 3626.903 | 35.0804 | 0.5843 | 1.91 | 281 | ENSG00000126653 | *NSRP1* | 635.4453 | 61.08828 | 0.6375 | 1.85 |
| 253 | ENSG00000157557 | *ETS2* | 2743.877 | 18.39712 | 0.5189 | 1.91 | 282 | ENSG00000241878 | *PISD* | 1755.022 | 75.8995 | 0.6524 | 1.85 |
| 254 | ENSG00000135480 | *KRT7* | 3391.252 | 293.8566 | 0.7150 | 1.91 | 283 | ENSG00000086062 | *B4GALT1* | 1470.094 | 74.51696 | 0.6513 | 1.85 |
| 255 | ENSG00000110955 | *ATP5B* | 14744.49 | 339.8558 | 0.7211 | 1.91 | 284 | ENSG00000122862 | *SRGN* | 26145.82 | 70.9097 | 0.6482 | 1.85 |
| 256 | ENSG00000179094 | *PER1* | 385.8495 | 32.19016 | 0.5778 | 1.90 | 285 | ENSG00000099821 | *POLRMT* | 1089.427 | 68.97757 | 0.6465 | 1.85 |
| 257 | ENSG00000005108 | *THSD7A* | 1508.581 | 150.5625 | 1.4618 | 1.90 | 286 | ENSG00000146072 | *TNFRSF21* | 1781.455 | 52.26842 | 0.6271 | 1.85 |
| 258 | ENSG00000103145 | *HCFC1R1* | 661.2923 | 30.29935 | 0.5727 | 1.90 | 287 | ENSG00000171763 | *SPATA5L1* | 507.541 | 15.97677 | 0.5139 | 1.85 |

| Rank | Ensembl Gene ID | Official gene symbol | Mean control counts* | Standard deviation control counts* | Mean HHT/ mean control | Ranking score |
|---|---|---|---|---|---|---|
| 288 | ENSG00000006744 | *ELAC2* | 2479.127 | 157.1501 | 0.6944 | 1.84 |
| 289 | ENSG00000162714 | *ZNF496* | 2733.898 | 116.6871 | 0.6792 | 1.84 |
| 290 | ENSG00000179409 | *GEMIN4* | 1087.6 | 17.78398 | 0.5285 | 1.84 |
| 291 | ENSG00000104177 | *MYEF2* | 1114.953 | 57.37623 | 0.6356 | 1.84 |
| 292 | ENSG00000187840 | *EIF4EBP1* | 1737.468 | 26.08839 | 0.5706 | 1.83 |
| 293 | ENSG00000108932 | *SLC16A6* | 117.0815 | 4.179912 | 3.5825 | 1.83 |
| 294 | ENSG00000166780 | *C16orf45* | 114.0515 | 5.659065 | 2.8610 | 1.82 |
| 295 | ENSG00000115594 | *IL1R1* | 493.4503 | 17.66545 | 1.8824 | 1.82 |
| 296 | ENSG00000117394 | *SLC2A1* | 700.4534 | 25.3542 | 0.5702 | 1.82 |
| 297 | ENSG00000112759 | *SLC29A1* | 3926.972 | 90.53132 | 0.6683 | 1.82 |
| 298 | ENSG00000108262 | *GIT1* | 1686.36 | 61.88953 | 0.6445 | 1.81 |
| 299 | ENSG00000139679 | *LPAR6* | 687.0562 | 62.33741 | 1.5492 | 1.81 |
| 300 | ENSG00000006831 | *ADIPOR2* | 1725.166 | 108.0342 | 0.6797 | 1.81 |
| 301 | ENSG00000131653 | *TRAF7* | 4127.001 | 93.85075 | 0.6719 | 1.81 |
| 302 | ENSG00000178996 | *SNX18* | 1866.557 | 142.3593 | 0.6948 | 1.81 |
| 303 | ENSG00000266028 | *SRGAP2* | 2001.776 | 123.2266 | 0.6872 | 1.81 |
| 304 | ENSG00000138386 | *NAB1* | 2258.049 | 151.2931 | 0.6981 | 1.80 |
| 305 | ENSG00000071462 | *WBSCR22* | 1110.61 | 76.48763 | 0.6598 | 1.80 |
| 306 | ENSG00000158470 | *B4GALT5* | 4384.163 | 274.5179 | 0.7254 | 1.80 |
| 307 | ENSG00000164975 | *SNAPC3* | 1347.473 | 124.4686 | 0.6884 | 1.80 |
| 308 | ENSG00000102978 | *POLR2C* | 1987.596 | 100.2573 | 0.6770 | 1.80 |
| 309 | ENSG00000147677 | *EIF3H* | 6127.979 | 157.3185 | 0.7010 | 1.80 |
| 310 | ENSG00000166265 | *CYYR1* | 1590.715 | 76.49718 | 0.6614 | 1.79 |
| 311 | ENSG00000106397 | *PLOD3* | 2089.132 | 58.76992 | 0.6447 | 1.79 |
| 312 | ENSG00000065600 | *TMEM206* | 640.3605 | 56.82412 | 0.6435 | 1.78 |
| 313 | ENSG00000071967 | *CYBRD1* | 2416.742 | 220.0854 | 0.7188 | 1.78 |
| 314 | ENSG00000177156 | *TALDO1* | 5394.404 | 21.50163 | 0.5600 | 1.78 |
| 315 | ENSG00000108179 | *PPIF* | 972.2601 | 42.00614 | 0.6217 | 1.78 |

| Rank | Ensembl Gene ID | Official gene symbol | Mean control counts* | Standard deviation control counts* | Mean HHT/ mean control | Ranking score |
|---|---|---|---|---|---|---|
| 316 | ENSG00000147324 | *MFHAS1* | 1999.953 | 175.3761 | 0.7098 | 1.77 |
| 317 | ENSG00000107263 | *RAPGEF1* | 2988.522 | 87.69377 | 0.6731 | 1.77 |
| 318 | ENSG00000121797 | *CCRL2* | 262.0503 | 17.04915 | 0.5356 | 1.77 |
| 319 | ENSG00000117643 | *MAN1C1* | 92.02452 | 3.87593 | 3.6894 | 1.77 |
| 320 | ENSG00000196365 | *LONP1* | 2242.129 | 25.41357 | 0.5790 | 1.77 |
| 321 | ENSG00000112511 | *PHF1* | 580.1375 | 46.59095 | 0.6314 | 1.77 |
| 322 | ENSG00000092445 | *TYRO3* | 1121.718 | 77.56216 | 0.6667 | 1.76 |
| 323 | ENSG00000103507 | *BCKDK* | 1226.819 | 33.19296 | 0.6044 | 1.76 |
| 324 | ENSG00000176692 | *FOXC2* | 685.9587 | 18.99473 | 0.5495 | 1.76 |
| 325 | ENSG00000087111 | *PIGS* | 1564.693 | 64.19437 | 0.6549 | 1.76 |
| 326 | ENSG00000142546 | *NOSIP* | 1763.059 | 83.89547 | 0.6721 | 1.76 |
| 327 | ENSG00000182963 | *GJC1* | 754.7106 | 22.11678 | 0.5665 | 1.76 |
| 328 | ENSG00000100311 | *PDGFB* | 5279.76 | 78.41093 | 0.6684 | 1.76 |
| 329 | ENSG00000144028 | *SNRNP200* | 3396.661 | 121.9269 | 0.6940 | 1.75 |
| 330 | ENSG00000062716 | *VMP1* | 6293.524 | 122.0672 | 1.4408 | 1.75 |
| 331 | ENSG00000166986 | *MARS* | 3202.705 | 129.6267 | 0.6975 | 1.75 |
| 332 | ENSG00000169756 | *LIMS1* | 5816.772 | 326.286 | 1.3530 | 1.75 |
| 333 | ENSG00000174684 | *B4GAT1* | 915.4285 | 45.21538 | 0.6321 | 1.75 |
| 334 | ENSG00000164086 | *DUSP7* | 711.7518 | 55.47169 | 0.6473 | 1.75 |
| 335 | ENSG00000109046 | *WSB1* | 18438.36 | 946.6341 | 0.7751 | 1.75 |
| 336 | ENSG00000182670 | *TTC3* | 8423.824 | 249.1742 | 0.7292 | 1.74 |
| 337 | ENSG00000142864 | *SERBP1* | 12091.3 | 390.425 | 0.7468 | 1.74 |
| 338 | ENSG00000168268 | *NT5DC2* | 2130.502 | 31.43858 | 0.6043 | 1.74 |
| 339 | ENSG00000138095 | *LRPPRC* | 6454.995 | 81.05173 | 0.6736 | 1.74 |
| 340 | ENSG00000156515 | *HK1* | 3987.265 | 278.134 | 0.7346 | 1.74 |
| 341 | ENSG00000104964 | *AES* | 4210.712 | 118.5674 | 0.6955 | 1.73 |
| 342 | ENSG00000158769 | *F11R* | 4519.198 | 240.6812 | 0.7288 | 1.73 |
| 343 | ENSG00000109971 | *HSPA8* | 47998.06 | 1217.674 | 0.7834 | 1.73 |
| 344 | ENSG00000227372 | *TP73-AS1* | 546.0175 | 40.12855 | 0.6254 | 1.73 |

| Rank | Ensembl Gene ID | Official gene symbol | Mean control counts* | Standard deviation control counts* | Mean HHT/ mean control | Ranking score |
|---|---|---|---|---|---|---|
| 345 | ENSG00000104885 | *DOT1L* | 493.4527 | 35.65034 | 0.6162 | 1.73 |
| 346 | ENSG00000198911 | *SREBF2* | 1723.041 | 63.32728 | 0.6589 | 1.73 |
| 347 | ENSG00000196305 | *IARS* | 7366.594 | 156.1729 | 0.7102 | 1.73 |
| 348 | ENSG00000159176 | *CSRP1* | 13372.38 | 321.7745 | 0.7413 | 1.73 |
| 349 | ENSG00000128512 | *DOCK4* | 10203.58 | 49.69819 | 0.6425 | 1.73 |
| 350 | ENSG00000213923 | *CSNK1E* | 3472.68 | 131.1268 | 0.7017 | 1.73 |
| 351 | ENSG00000169957 | *ZNF768* | 795.9477 | 28.99602 | 0.5990 | 1.73 |
| 352 | ENSG00000161970 | *RPL26* | 25134.41 | 649.5969 | 0.7663 | 1.72 |
| 353 | ENSG00000162409 | *PRKAA2* | 364.6371 | 31.4797 | 0.6068 | 1.72 |
| 354 | ENSG00000073756 | *PTGS2* | 7624.36 | 310.78 | 1.3500 | 1.72 |
| 355 | ENSG00000112584 | *FAM120B* | 988.3799 | 67.77422 | 0.6648 | 1.72 |
| 356 | ENSG00000197971 | *MBP* | 604.5684 | 38.46618 | 0.6240 | 1.72 |
| 357 | ENSG00000140400 | *MAN2C1* | 1529.421 | 20.07732 | 0.5635 | 1.72 |
| 358 | ENSG00000204469 | *PRRC2A* | 2325.185 | 37.65012 | 0.6225 | 1.72 |
| 359 | ENSG00000145882 | *PCYOX1L* | 651.3449 | 41.48493 | 0.6303 | 1.72 |
| 360 | ENSG00000135272 | *MDFIC* | 670.6066 | 47.72062 | 1.5601 | 1.72 |
| 361 | ENSG00000124608 | *AARS2* | 597.155 | 51.83239 | 0.6471 | 1.72 |
| 362 | ENSG00000163882 | *POLR2H* | 1363.216 | 54.50447 | 0.6509 | 1.72 |
| 363 | ENSG00000167965 | *MLST8* | 709.9697 | 31.33565 | 0.6076 | 1.72 |
| 364 | ENSG00000156110 | *ADK* | 1235.814 | 27.67636 | 0.5969 | 1.71 |
| 365 | ENSG00000167291 | *TBC1D16* | 2259.589 | 201.0253 | 0.7245 | 1.71 |
| 366 | ENSG00000126458 | *RRAS* | 1777.065 | 121.1023 | 0.7004 | 1.71 |
| 367 | ENSG00000153246 | *PLA2R1* | 2527.501 | 95.7164 | 0.6878 | 1.71 |
| 368 | ENSG00000136026 | *CKAP4* | 8840.603 | 76.48302 | 0.6748 | 1.71 |
| 369 | ENSG00000050165 | *DKK3* | 9713.636 | 39.91227 | 0.6296 | 1.71 |
| 370 | ENSG00000170191 | *NANP* | 722.276 | 58.82671 | 0.6582 | 1.70 |
| 371 | ENSG00000105223 | *PLD3* | 4313.425 | 47.66657 | 0.6443 | 1.70 |
| 372 | ENSG00000155229 | *MMS19* | 2891.564 | 81.71088 | 0.6806 | 1.69 |
| 373 | ENSG00000172578 | *KLHL6* | 1093.043 | 26.92227 | 0.5980 | 1.69 |
| 374 | ENSG00000004487 | *KDM1A* | 3692.216 | 144.7399 | 0.7117 | 1.69 |
| 375 | ENSG00000185022 | *MAFF* | 6789.497 | 478.4692 | 0.7604 | 1.69 |
| 376 | ENSG00000081181 | *ARG2* | 416.1202 | 9.17811 | 0.4669 | 1.69 |
| 377 | ENSG00000197461 | *PDGFA* | 2899.368 | 53.83886 | 0.6549 | 1.69 |
| 378 | ENSG00000137831 | *UACA* | 4318.608 | 128.0181 | 1.4158 | 1.69 |
| 379 | ENSG00000104903 | *LYL1* | 1489.573 | 87.65459 | 0.6862 | 1.68 |
| 380 | ENSG00000137094 | *DNAJB5* | 527.2922 | 52.64155 | 0.6538 | 1.68 |
| 381 | ENSG00000170275 | *CRTAP* | 3343.306 | 98.03383 | 0.6930 | 1.68 |
| 382 | ENSG00000205937 | *RNPS1* | 2142.269 | 115.81 | 0.7023 | 1.68 |
| 383 | ENSG00000196428 | *TSC22D2* | 3091.773 | 40.08736 | 0.6348 | 1.68 |
| 384 | ENSG00000100714 | *MTHFD1* | 1562.57 | 136.2326 | 0.7118 | 1.67 |
| 385 | ENSG00000115380 | *EFEMP1* | 173366.2 | 1131.027 | 0.7885 | 1.67 |
| 386 | ENSG00000164828 | *SUN1* | 2698.865 | 164.8672 | 0.7210 | 1.67 |
| 387 | ENSG00000170873 | *MTSS1* | 1163.954 | 29.86943 | 0.6118 | 1.67 |
| 388 | ENSG00000134330 | *IAH1* | 789.4456 | 25.56481 | 0.5976 | 1.67 |
| 389 | ENSG00000107130 | *NCS1* | 2274.146 | 62.2748 | 0.6681 | 1.67 |
| 390 | ENSG00000130522 | *JUND* | 2411.924 | 38.90314 | 0.6350 | 1.66 |
| 391 | ENSG00000156011 | *PSD3* | 1546.571 | 39.25032 | 0.6359 | 1.66 |
| 392 | ENSG00000107443 | *CCNJ* | 1438.238 | 94.31409 | 0.6940 | 1.66 |
| 393 | ENSG00000103510 | *KAT8* | 748.6752 | 31.31329 | 0.6177 | 1.66 |
| 394 | ENSG00000150938 | *CRIM1* | 45473.65 | 212.4166 | 0.7337 | 1.66 |
| 395 | ENSG00000102178 | *UBL4A* | 1105.259 | 42.26419 | 0.6421 | 1.66 |
| 396 | ENSG00000120318 | *ARAP3* | 2380.973 | 89.89007 | 0.6921 | 1.66 |
| 397 | ENSG00000107554 | *DNMBP* | 2888.314 | 105.899 | 0.7013 | 1.65 |
| 398 | ENSG00000136240 | *KDELR2* | 5575.67 | 116.9988 | 0.7066 | 1.65 |
| 399 | ENSG00000115604 | *IL18R1* | 671.7737 | 30.63748 | 0.6168 | 1.65 |
| 400 | ENSG00000101901 | *ALG13* | 889.2495 | 56.65491 | 0.6640 | 1.65 |
| 401 | ENSG00000061987 | *MON2* | 2465.773 | 127.0403 | 0.7111 | 1.65 |
| 402 | ENSG00000182934 | *SRPR* | 1611.676 | 66.78609 | 0.6750 | 1.65 |

| Rank | Ensembl Gene ID | Official gene symbol | Mean control counts* | Standard deviation control counts* | Mean HHT/ mean control | Ranking score | Rank | Ensembl Gene ID | Official gene symbol | Mean control counts* | Standard deviation control counts* | Mean HHT/ mean control | Ranking score |
|---|---|---|---|---|---|---|---|---|---|---|---|---|---|
| 403 | ENSG00000091592 | *NLRP1* | 1779.605 | 35.24242 | 0.6292 | 1.65 | 432 | ENSG00000169908 | *TM4SF1* | 16973.72 | 614.6765 | 0.7786 | 1.61 |
| 404 | ENSG00000116337 | *AMPD2* | 1442.828 | 18.80224 | 0.5699 | 1.65 | 433 | ENSG00000109113 | *RAB34* | 1370.642 | 66.44868 | 0.6820 | 1.61 |
| 405 | ENSG00000048342 | *CC2D2A* | 739.8522 | 60.62637 | 0.6691 | 1.65 | 434 | ENSG00000116521 | *SCAMP3* | 1177.649 | 116.3117 | 0.7136 | 1.60 |
| 406 | ENSG00000140396 | *NCOA2* | 374.6811 | 20.60456 | 1.7241 | 1.65 | 435 | ENSG00000105339 | *DENND3* | 2858.91 | 26.43463 | 0.6127 | 1.60 |
| 407 | ENSG00000117318 | *ID3* | 13709.01 | 1346.879 | 0.7957 | 1.65 | 436 | ENSG00000184205 | *TSPYL2* | 1219.645 | 18.83954 | 0.5799 | 1.60 |
| 408 | ENSG00000105063 | *PPP6R1* | 1185.889 | 63.69617 | 0.6733 | 1.64 | 437 | ENSG00000187109 | *NAP1L1* | 20788.64 | 361.1171 | 0.7621 | 1.60 |
| 409 | ENSG00000114867 | *EIF4G1* | 11613.65 | 378.533 | 0.7582 | 1.64 | 438 | ENSG00000234616 | *JRK* | 798.4413 | 31.08141 | 0.6279 | 1.60 |
| 410 | ENSG00000131669 | *NINJ1* | 1900.051 | 55.70876 | 0.6646 | 1.64 | 439 | ENSG00000148337 | *CIZ1* | 1487.098 | 22.48438 | 0.5985 | 1.60 |
| 411 | ENSG00000130159 | *ECSIT* | 670.0702 | 51.28517 | 0.6598 | 1.64 | 440 | ENSG00000171853 | *TRAPPC12* | 954.7489 | 86.41222 | 0.6989 | 1.60 |
| 412 | ENSG00000170004 | *CHD3* | 5246.563 | 184.6954 | 0.7317 | 1.63 | 441 | ENSG00000076356 | *PLXNA2* | 7561.753 | 69.3177 | 0.6864 | 1.59 |
| 413 | ENSG00000159842 | *ABR* | 2155.994 | 112.6414 | 0.7086 | 1.63 | 442 | ENSG00000188177 | *ZC3H6* | 499.4026 | 26.08321 | 0.6137 | 1.59 |
| 414 | ENSG00000128578 | *STRIP2* | 851.6011 | 9.023854 | 0.4775 | 1.63 | 443 | ENSG00000141971 | *MVB12A* | 1008.095 | 79.71462 | 0.6953 | 1.59 |
| 415 | ENSG00000105655 | *ISYNA1* | 960.3556 | 25.17062 | 0.6042 | 1.63 | 444 | ENSG00000140553 | *UNC45A* | 3141.593 | 46.00728 | 0.6601 | 1.59 |
| 416 | ENSG00000162191 | *UBXN1* | 2640.088 | 115.4141 | 0.7101 | 1.63 | 445 | ENSG00000160226 | *C21orf2* | 235.0877 | 16.87283 | 0.5704 | 1.59 |
| 417 | ENSG00000148175 | *STOM* | 10451.63 | 508.6169 | 0.7709 | 1.62 | 446 | ENSG00000182903 | *ZNF721* | 552.9529 | 55.26029 | 0.6736 | 1.59 |
| 418 | ENSG00000130816 | *DNMT1* | 1692.965 | 157.8965 | 0.7259 | 1.62 | 447 | ENSG00000050405 | *LIMA1* | 6587.287 | 249.903 | 0.7504 | 1.59 |
| 419 | ENSG00000118579 | *MED28* | 1405.387 | 99.13287 | 0.7028 | 1.62 | 448 | ENSG00000108641 | *B9D1* | 333.0106 | 27.10269 | 0.6191 | 1.58 |
| 420 | ENSG00000157933 | *SKI* | 1460.454 | 83.94097 | 0.6938 | 1.62 | 449 | ENSG00000184916 | *JAG2* | 2117.522 | 75.04794 | 0.6933 | 1.58 |
| 421 | ENSG00000100412 | *ACO2* | 1445.392 | 77.14267 | 0.6890 | 1.62 | 450 | ENSG00000137177 | *KIF13A* | 4880.716 | 141.5101 | 0.7266 | 1.58 |
| 422 | ENSG00000142396 | *ERVK3-1* | 451.6879 | 22.01097 | 0.5924 | 1.62 | 451 | ENSG00000117676 | *RPS6KA1* | 447.0375 | 17.71002 | 0.5771 | 1.58 |
| 423 | ENSG00000173020 | *ADRBK1* | 1527.688 | 96.88482 | 0.7021 | 1.62 | 452 | ENSG00000152465 | *NMT2* | 6892.237 | 86.25171 | 0.7016 | 1.58 |
| 424 | ENSG00000138835 | *RGS3* | 10132.92 | 405.1796 | 0.7640 | 1.62 | 453 | ENSG00000136270 | *TBRG4* | 1143.361 | 31.61196 | 0.6332 | 1.58 |
| 425 | ENSG00000119139 | *TJP2* | 5391.763 | 283.4144 | 0.7512 | 1.62 | 454 | ENSG00000182158 | *CREB3L2* | 1794.288 | 29.01463 | 0.6262 | 1.58 |
| 426 | ENSG00000133800 | *LYVE1* | 1458.165 | 7.209668 | 2.2651 | 1.62 | 455 | ENSG00000162231 | *NXF1* | 1498.994 | 110.8584 | 0.7156 | 1.58 |
| 427 | ENSG00000124762 | *CDKN1A* | 14273.93 | 664.3991 | 0.7800 | 1.62 | 456 | ENSG00000174238 | *PITPNA* | 2353.369 | 94.34327 | 0.7072 | 1.58 |
| 428 | ENSG00000204713 | *TRIM27* | 2536.933 | 50.90284 | 0.6632 | 1.61 | 457 | ENSG00000105220 | *GPI* | 4831.582 | 110.3601 | 0.7159 | 1.57 |
| 429 | ENSG00000198925 | *ATG9A* | 1345.539 | 95.86649 | 0.7028 | 1.61 | 458 | ENSG00000138867 | *GUCD1* | 3201.614 | 304.0287 | 0.7597 | 1.57 |
| 430 | ENSG00000102606 | *ARHGEF7* | 3817.722 | 262.8373 | 0.7492 | 1.61 | 459 | ENSG00000111348 | *ARHGDIB* | 6112.515 | 329.0576 | 0.7627 | 1.57 |
| 431 | ENSG00000074695 | *LMAN1* | 7466.759 | 88.87469 | 0.6988 | 1.61 | 460 | ENSG00000102316 | *MAGED2* | 7770.226 | 316.2934 | 0.7613 | 1.57 |

| Rank | Ensembl Gene ID | Official gene symbol | Mean control counts* | Standard deviation control counts* | Mean HHT/ mean control | Ranking score |
|---|---|---|---|---|---|---|
| 461 | ENSG00000233817 | | 194.5328 | 0.1308221 | 0.4626 | 1.57 |
| 462 | ENSG00000154229 | *PRKCA* | 1777.933 | 53.58548 | 0.6746 | 1.57 |
| 463 | ENSG00000137801 | *THBS1* | 312144.5 | 673.0472 | 0.7861 | 1.57 |
| 464 | ENSG00000167004 | *PDIA3* | 13372.05 | 279.6207 | 0.7575 | 1.56 |
| 465 | ENSG00000132612 | *VPS4A* | 1354.622 | 59.64341 | 0.6821 | 1.56 |
| 466 | ENSG00000221968 | *FADS3* | 5640.128 | 76.27817 | 0.6971 | 1.56 |
| 467 | ENSG00000117419 | *ERI3* | 1063.934 | 75.10242 | 0.6962 | 1.56 |
| 468 | ENSG00000103148 | *NPRL3* | 214.3356 | 21.12138 | 0.5993 | 1.56 |
| 469 | ENSG00000184207 | *PGP* | 645.983 | 59.15527 | 0.6821 | 1.56 |
| 470 | ENSG00000213965 | *NUDT19* | 660.7131 | 30.84701 | 0.6344 | 1.56 |
| 471 | ENSG00000143799 | *PARP1* | 3840.642 | 135.3138 | 0.7277 | 1.56 |
| 472 | ENSG00000101298 | *SNPH* | 534.5012 | 35.37058 | 0.6462 | 1.56 |
| 473 | ENSG00000143811 | *PYCR2* | 764.4044 | 37.22883 | 0.6508 | 1.55 |
| 474 | ENSG00000101236 | *RNF24* | 685.6805 | 57.77467 | 0.6819 | 1.55 |
| 475 | ENSG00000221978 | *CCNL2* | 2895.046 | 126.6419 | 0.7256 | 1.55 |
| 476 | ENSG00000261371 | *PECAM1* | 62393.18 | 3044.904 | 0.8241 | 1.55 |
| 477 | ENSG00000167553 | *TUBA1C* | 4107.264 | 157.4256 | 0.7359 | 1.55 |
| 478 | ENSG00000125968 | *ID1* | 19487.92 | 1636.917 | 0.8109 | 1.55 |
| 479 | ENSG00000205726 | *ITSN1* | 2981.897 | 168.0963 | 0.7390 | 1.55 |
| 480 | ENSG00000196923 | *PDLIM7* | 2077.643 | 66.37191 | 0.6911 | 1.55 |
| 481 | ENSG00000133706 | *LARS* | 6187.719 | 169.2639 | 0.7396 | 1.55 |
| 482 | ENSG00000170043 | *TRAPPC1* | 1355.244 | 59.08475 | 0.6843 | 1.55 |
| 483 | ENSG00000071564 | *TCF3* | 2116.49 | 77.911 | 0.7011 | 1.55 |
| 484 | ENSG00000129351 | *ILF3* | 4997.54 | 375.469 | 0.7705 | 1.55 |
| 485 | ENSG00000131473 | *ACLY* | 7724.338 | 93.01312 | 0.7111 | 1.55 |
| 486 | ENSG00000177697 | *CD151* | 10114.76 | 571.1579 | 0.7842 | 1.54 |
| 487 | ENSG00000197261 | *C6orf141* | 475.4343 | 43.83952 | 0.6651 | 1.54 |
| 488 | ENSG00000151929 | *BAG3* | 1656.628 | 36.35907 | 0.6512 | 1.54 |
| 489 | ENSG00000101191 | *DIDO1* | 2495.067 | 141.8128 | 0.7327 | 1.54 |
| 490 | ENSG00000175309 | *PHYKPL* | 916.4744 | 40.93031 | 0.6603 | 1.54 |
| 491 | ENSG00000156990 | *RPUSD3* | 505.3572 | 18.81482 | 0.5918 | 1.54 |
| 492 | ENSG00000110697 | *PITPNM1* | 631.2054 | 15.05584 | 0.5673 | 1.54 |
| 493 | ENSG00000101220 | *C20orf27* | 790.7047 | 51.84339 | 0.6776 | 1.54 |
| 494 | ENSG00000149930 | *TAOK2* | 4014.004 | 189.1162 | 0.7460 | 1.54 |
| 495 | ENSG00000165724 | *ZMYND19* | 530.1313 | 19.1032 | 0.5941 | 1.54 |
| 496 | ENSG00000158863 | *FAM160B2* | 802.9189 | 35.97272 | 0.6514 | 1.54 |
| 497 | ENSG00000115419 | *GLS* | 7171.133 | 519.5792 | 0.7823 | 1.54 |
| 498 | ENSG00000197386 | *HTT* | 1833.36 | 120.747 | 0.7260 | 1.53 |
| 499 | ENSG00000142102 | *ATHL1* | 889.1013 | 41.01282 | 0.6617 | 1.53 |
| 500 | ENSG00000163597 | *SNHG16* | 6264.914 | 510.444 | 0.7820 | 1.53 |
| 501 | ENSG00000066629 | *EML1* | 1780.745 | 21.00328 | 0.6045 | 1.53 |
| 502 | ENSG00000116221 | *MRPL37* | 2175.427 | 64.70961 | 0.6925 | 1.53 |
| 503 | ENSG00000105976 | *MET* | 3559.515 | 144.4102 | 0.7351 | 1.53 |
| 504 | ENSG00000197043 | *ANXA6* | 8308.114 | 153.2652 | 0.7379 | 1.53 |
| 505 | ENSG00000196547 | *MAN2A2* | 1101.814 | 50.01856 | 0.6766 | 1.53 |
| 506 | ENSG00000135052 | *GOLM1* | 3299.553 | 58.95192 | 0.6874 | 1.53 |
| 507 | ENSG00000124588 | *NQO2* | 777.5141 | 11.57331 | 0.5358 | 1.53 |
| 508 | ENSG00000171469 | *ZNF561* | 950.9827 | 59.47025 | 0.6881 | 1.53 |
| 509 | ENSG00000125686 | *MED1* | 1566.457 | 122.0887 | 0.7280 | 1.53 |
| 510 | ENSG00000105135 | *ILVBL* | 996.5271 | 27.3289 | 0.6307 | 1.52 |
| 511 | ENSG00000031698 | *SARS* | 5175.287 | 228.9632 | 0.7554 | 1.52 |
| 512 | ENSG00000125818 | *PSMF1* | 1746.805 | 110.5768 | 0.7235 | 1.52 |
| 513 | ENSG00000141524 | *TMC6* | 626.2833 | 57.0908 | 0.6863 | 1.52 |
| 514 | ENSG00000015532 | *XYLT2* | 896.5685 | 33.33464 | 0.6478 | 1.52 |
| 515 | ENSG00000101224 | *CDC25B* | 951.2791 | 28.18607 | 0.6342 | 1.52 |
| 516 | ENSG00000118900 | *UBN1* | 2190.011 | 117.0382 | 0.7268 | 1.52 |
| 517 | ENSG00000114353 | *GNAI2* | 24424.07 | 1268.298 | 0.8085 | 1.52 |
| 518 | ENSG00000087269 | *NOP14* | 2166.77 | 29.67298 | 0.6389 | 1.52 |

| Rank | Ensembl Gene ID | Official gene symbol | Mean control counts* | Standard deviation control counts* | Mean HHT/ mean control | Ranking score | Rank | Ensembl Gene ID | Official gene symbol | Mean control counts* | Standard deviation control counts* | Mean HHT/ mean control | Ranking score |
|---|---|---|---|---|---|---|---|---|---|---|---|---|---|
| 519 | ENSG00000189306 | *RRP7A* | 1433.44 | 70.23519 | 0.6998 | 1.52 | 548 | ENSG00000089057 | *SLC23A2* | 1175.901 | 70.61572 | 0.7049 | 1.49 |
| 520 | ENSG00000197961 | *ZNF121* | 2149.618 | 82.97044 | 0.7095 | 1.52 | 549 | ENSG00000145014 | *TMEM44* | 1832.823 | 117.1824 | 0.7318 | 1.49 |
| 521 | ENSG00000151474 | *FRMD4A* | 2844.476 | 147.7211 | 0.7382 | 1.52 | 550 | ENSG00000088298 | *EDEM2* | 1811.633 | 50.27496 | 0.6842 | 1.49 |
| 522 | ENSG00000078596 | *ITM2A* | 1985.196 | 88.21785 | 0.7131 | 1.51 | 551 | ENSG00000178467 | *P4HTM* | 825.2557 | 12.80153 | 0.5582 | 1.49 |
| 523 | ENSG00000149577 | *SIDT2* | 815.6697 | 16.5375 | 0.5830 | 1.51 | 552 | ENSG00000151779 | *NBAS* | 4317.909 | 211.0832 | 0.7576 | 1.49 |
| 524 | ENSG00000112419 | *PHACTR2* | 5731.099 | 402.5511 | 1.2870 | 1.51 | 553 | ENSG00000083444 | *PLOD1* | 6111.345 | 258.6587 | 0.7654 | 1.49 |
| 525 | ENSG00000188785 | *ZNF548* | 514.8735 | 37.87503 | 0.6594 | 1.51 | 554 | ENSG00000125812 | *GZF1* | 1978.99 | 53.50246 | 0.6889 | 1.48 |
| 526 | ENSG00000112159 | *MDN1* | 3500.552 | 158.0171 | 0.7417 | 1.51 | 555 | ENSG00000071794 | *HLTF* | 1656.69 | 63.16246 | 0.6995 | 1.48 |
| 527 | ENSG00000117020 | *AKT3* | 4152.444 | 167.2999 | 0.7443 | 1.51 | 556 | ENSG00000140465 | *CYP1A1* | 476.195 | 35.05337 | 0.6595 | 1.48 |
| 528 | ENSG00000062822 | *POLD1* | 759.7628 | 28.25975 | 0.6364 | 1.51 | 557 | ENSG00000140945 | *CDH13* | 4758.811 | 98.232 | 0.7246 | 1.48 |
| 529 | ENSG00000131238 | *PPT1* | 3212.457 | 61.4144 | 0.6931 | 1.51 | 558 | ENSG00000188554 | *NBR1* | 4664.066 | 247.5918 | 0.7649 | 1.48 |
| 530 | ENSG00000134308 | *YWHAQ* | 12603.44 | 254.5905 | 0.7615 | 1.51 | 559 | ENSG00000099219 | *ERMP1* | 1776.645 | 23.70426 | 0.6276 | 1.47 |
| 531 | ENSG00000100813 | *ACIN1* | 3410.063 | 316.2501 | 0.7695 | 1.51 | 560 | ENSG00000153029 | *MR1* | 94.04902 | 5.336035 | 2.4052 | 1.47 |
| 532 | ENSG00000140521 | *POLG* | 1680.033 | 39.97221 | 0.6645 | 1.51 | 561 | ENSG00000130175 | *PRKCSH* | 3817.135 | 41.74363 | 0.6750 | 1.47 |
| 533 | ENSG00000139629 | *GALNT6* | 2900.542 | 123.1916 | 0.7313 | 1.51 | 562 | ENSG00000169871 | *TRIM56* | 1347.493 | 36.90076 | 0.6660 | 1.47 |
| 534 | ENSG00000132361 | *CLUH* | 1326.281 | 26.76909 | 0.6324 | 1.51 | 563 | ENSG00000173457 | *PPP1R14B* | 4105.937 | 141.8785 | 0.7439 | 1.47 |
| 535 | ENSG00000171016 | *PYGO1* | 306.2389 | 12.19924 | 0.5481 | 1.50 | 564 | ENSG00000169188 | *APEX2* | 448.0272 | 37.00873 | 0.6664 | 1.47 |
| 536 | ENSG00000127054 | *CPSF3L* | 1816.176 | 101.372 | 0.7221 | 1.50 | 565 | ENSG00000113732 | *ATP6V0E1* | 2916.219 | 79.11119 | 1.3984 | 1.47 |
| 537 | ENSG00000134759 | *ELP2* | 2687.537 | 128.9394 | 0.7341 | 1.50 | 566 | ENSG00000197226 | *TBC1D9B* | 4306.64 | 74.09753 | 0.7116 | 1.46 |
| 538 | ENSG00000100368 | *CSF2RB* | 998.5135 | 23.18097 | 0.6206 | 1.50 | 567 | ENSG00000128283 | *CDC42EP1* | 2125.434 | 35.79985 | 0.6642 | 1.46 |
| 539 | ENSG00000103994 | *ZNF106* | 4929.476 | 141.723 | 0.7394 | 1.50 | 568 | ENSG00000178057 | *NDUFAF3* | 1042.913 | 65.54838 | 0.7050 | 1.46 |
| 540 | ENSG00000115806 | *GORASP2* | 2192.76 | 92.10542 | 0.7185 | 1.50 | 569 | ENSG00000107566 | *ERLIN1* | 1575.035 | 72.48986 | 0.7108 | 1.46 |
| 541 | ENSG00000164251 | *F2RL1* | 501.7907 | 13.77112 | 1.7680 | 1.49 | 570 | ENSG00000153885 | *KCTD15* | 1882.308 | 177.6048 | 0.7543 | 1.46 |
| 542 | ENSG00000186470 | *BTN3A2* | 1107.093 | 29.49961 | 0.6431 | 1.49 | 571 | ENSG00000135372 | *NAT10* | 1064.316 | 71.34538 | 0.7102 | 1.46 |
| 543 | ENSG00000117632 | *STMN1* | 4135.625 | 32.55201 | 1.5347 | 1.49 | 572 | ENSG00000177302 | *TOP3A* | 783.4775 | 71.88484 | 0.7109 | 1.46 |
| 544 | ENSG00000204574 | *ABCF1* | 2764.334 | 230.7972 | 0.7603 | 1.49 | 573 | ENSG00000278540 | *ACACA* | 2330.094 | 78.26307 | 0.7156 | 1.46 |
| 545 | ENSG00000073849 | *ST6GAL1* | 4327.266 | 231.5214 | 0.7606 | 1.49 | 574 | ENSG00000182197 | *EXT1* | 4272.619 | 48.95049 | 0.6874 | 1.46 |
| 546 | ENSG00000122203 | *KIAA1191* | 3206.999 | 107.0139 | 0.7270 | 1.49 | 575 | ENSG00000160072 | *ATAD3B* | 916.6606 | 74.33797 | 0.7128 | 1.46 |
| 547 | ENSG00000161904 | *LEMD2* | 1824.045 | 119.7057 | 0.7325 | 1.49 | 576 | ENSG00000169727 | *GPS1* | 1866.937 | 49.22731 | 0.6878 | 1.46 |

| Rank | Ensembl Gene ID | Official gene symbol | Mean control counts* | Standard deviation control counts* | Mean HHT/ mean control | Ranking score |
|---|---|---|---|---|---|---|
| 577 | ENSG00000127663 | *KDM4B* | 936.1021 | 38.42585 | 0.6706 | 1.46 |
| 578 | ENSG00000135723 | *FHOD1* | 2065.157 | 31.25471 | 0.6550 | 1.46 |
| 579 | ENSG00000144029 | *MRPS5* | 1339.986 | 40.31292 | 0.6744 | 1.46 |
| 580 | ENSG00000164466 | *SFXN1* | 2009.062 | 62.88507 | 0.7036 | 1.46 |
| 581 | ENSG00000179832 | *MROH1* | 1139.094 | 47.37617 | 0.6860 | 1.45 |
| 582 | ENSG00000136802 | *LRRC8A* | 5434.46 | 91.11959 | 0.7246 | 1.45 |
| 583 | ENSG00000184428 | *TOP1MT* | 778.8025 | 18.30792 | 0.6067 | 1.45 |
| 584 | ENSG00000146112 | *PPP1R18* | 2752.278 | 139.7979 | 0.7453 | 1.45 |
| 585 | ENSG00000071994 | *PDCD2* | 2773.529 | 125.1778 | 0.7405 | 1.45 |
| 586 | ENSG00000167815 | *PRDX2* | 3364.945 | 95.49127 | 0.7276 | 1.45 |
| 587 | ENSG00000089159 | *PXN* | 4376.562 | 37.41291 | 0.6702 | 1.45 |
| 588 | ENSG00000085998 | *POMGNT1* | 1775.092 | 72.01444 | 0.7125 | 1.45 |
| 589 | ENSG00000170266 | *GLB1* | 3239.089 | 221.568 | 0.7646 | 1.45 |
| 590 | ENSG00000151726 | *ACSL1* | 720.2966 | 50.61079 | 0.6912 | 1.45 |
| 591 | ENSG00000140632 | *GLYR1* | 2149.522 | 172.2324 | 0.7548 | 1.45 |
| 592 | ENSG00000102125 | *TAZ* | 509.3365 | 38.6908 | 0.6730 | 1.45 |
| 593 | ENSG00000107959 | *PITRM1* | 4083.365 | 30.6985 | 0.6554 | 1.45 |
| 594 | ENSG00000168887 | *C2orf68* | 586.8013 | 32.95806 | 0.6614 | 1.44 |
| 595 | ENSG00000000971 | *CFH* | 2959.012 | 125.2184 | 0.7417 | 1.44 |
| 596 | ENSG00000072571 | *HMMR* | 191.651 | 14.29376 | 1.7195 | 1.44 |
| 597 | ENSG00000165458 | *INPPL1* | 2248.548 | 39.86547 | 0.6766 | 1.44 |
| 598 | ENSG00000135018 | *UBQLN1* | 4561.454 | 28.11972 | 0.6497 | 1.44 |
| 599 | ENSG00000152990 | *ADGRA3* | 845.2201 | 25.25314 | 0.6405 | 1.44 |
| 600 | ENSG00000138449 | *SLC40A1* | 845.8461 | 67.85983 | 1.4063 | 1.44 |
| 601 | ENSG00000065357 | *DGKA* | 919.6233 | 79.96886 | 0.7203 | 1.44 |
| 602 | ENSG00000148606 | *POLR3A* | 1063.389 | 26.5426 | 0.6453 | 1.44 |
| 603 | ENSG00000115042 | *FAHD2A* | 599.9291 | 26.29339 | 0.6447 | 1.44 |
| 604 | ENSG00000122126 | *OCRL* | 1177.16 | 44.33714 | 0.6850 | 1.43 |
| 605 | ENSG00000117280 | *RAB29* | 1645.466 | 133.7519 | 0.7460 | 1.43 |
| 606 | ENSG00000076604 | *TRAF4* | 1212.696 | 77.97131 | 0.7196 | 1.43 |
| 607 | ENSG00000124181 | *PLCG1* | 3746.054 | 73.09739 | 0.7162 | 1.43 |
| 608 | ENSG00000130695 | *CEP85* | 539.7308 | 13.94521 | 0.5808 | 1.43 |
| 609 | ENSG00000184281 | *TSSC4* | 1197.198 | 69.02351 | 0.7131 | 1.43 |
| 610 | ENSG00000237441 | *RGL2* | 1895.598 | 101.2892 | 0.7334 | 1.43 |
| 611 | ENSG00000188153 | *COL4A5* | 3816.218 | 210.204 | 0.7653 | 1.43 |
| 612 | ENSG00000104635 | *SLC39A14* | 3103.539 | 97.59982 | 0.7319 | 1.43 |
| 613 | ENSG00000174851 | *YIF1A* | 1446.536 | 120.4593 | 0.7421 | 1.43 |
| 614 | ENSG00000111481 | *COPZ1* | 3367.182 | 124.5948 | 0.7436 | 1.43 |
| 615 | ENSG00000131408 | *NR1H2* | 1547.02 | 125.6645 | 0.7440 | 1.43 |
| 616 | ENSG00000241973 | *PI4KA* | 1207.49 | 115.2885 | 0.7404 | 1.43 |
| 617 | ENSG00000079805 | *DNM2* | 3001.044 | 92.43269 | 0.7296 | 1.43 |
| 618 | ENSG00000214706 | *IFRD2* | 1526.847 | 26.52398 | 0.6475 | 1.42 |
| 619 | ENSG00000111885 | *MAN1A1* | 661.6671 | 16.7998 | 1.6563 | 1.42 |
| 620 | ENSG00000073792 | *IGF2BP2* | 2358.143 | 120.7361 | 0.7431 | 1.42 |
| 621 | ENSG00000116016 | *EPAS1* | 15703.07 | 234.2588 | 0.7704 | 1.42 |
| 622 | ENSG00000173210 | *ABLIM3* | 2066.744 | 83.22648 | 0.7249 | 1.42 |
| 623 | ENSG00000084207 | *GSTP1* | 8964.152 | 224.5532 | 0.7691 | 1.42 |
| 624 | ENSG00000090565 | *RAB11FIP3* | 1159.846 | 85.37038 | 0.7265 | 1.42 |
| 625 | ENSG00000116260 | *QSOX1* | 5937.504 | 373.371 | 0.7867 | 1.42 |
| 626 | ENSG00000186480 | *INSIG1* | 1965.394 | 109.2084 | 0.7390 | 1.42 |
| 627 | ENSG00000105197 | *TIMM50* | 1438.452 | 124.8777 | 0.7453 | 1.42 |
| 628 | ENSG00000100139 | *MICALL1* | 1833.499 | 76.87405 | 0.7214 | 1.42 |
| 629 | ENSG00000183688 | *FAM101B* | 2195.494 | 121.9105 | 0.7448 | 1.41 |
| 630 | ENSG00000198089 | *SFI1* | 490.4361 | 35.59319 | 0.6730 | 1.41 |
| 631 | ENSG00000149782 | *PLCB3* | 2279.08 | 52.41599 | 0.6997 | 1.41 |
| 632 | ENSG00000111897 | *SERINC1* | 4312.094 | 135.1296 | 1.3339 | 1.41 |
| 633 | ENSG00000099942 | *CRKL* | 2676.73 | 155.4466 | 0.7558 | 1.41 |
| 634 | ENSG00000141027 | *NCOR1* | 3814.371 | 280.5458 | 0.7783 | 1.41 |

| Rank | Ensembl Gene ID | Official gene symbol | Mean control counts* | Standard deviation control counts* | Mean HHT/ mean control | Ranking score | Rank | Ensembl Gene ID | Official gene symbol | Mean control counts* | Standard deviation control counts* | Mean HHT/ mean control | Ranking score |
|---|---|---|---|---|---|---|---|---|---|---|---|---|---|
| 635 | ENSG00000149658 | *YTHDF1* | 1738.312 | 43.13344 | 0.6873 | 1.41 | 664 | ENSG00000134480 | *CCNH* | 1739.342 | 93.58616 | 0.7368 | 1.39 |
| 636 | ENSG00000100650 | *SRSF5* | 8040.745 | 223.4105 | 0.7706 | 1.41 | 665 | ENSG00000146223 | *RPL7L1* | 5281.229 | 188.569 | 0.7675 | 1.39 |
| 637 | ENSG00000149798 | *CDC42EP2* | 607.3829 | 17.1932 | 0.6094 | 1.41 | 666 | ENSG00000067829 | *IDH3G* | 968.3593 | 29.28426 | 0.6634 | 1.39 |
| 638 | ENSG00000104131 | *EIF3J* | 1790.805 | 85.81877 | 0.7287 | 1.41 | 667 | ENSG00000125656 | *CLPP* | 1016.074 | 33.35649 | 0.6737 | 1.39 |
| 639 | ENSG00000128595 | *CALU* | 16729.92 | 243.9522 | 0.7741 | 1.41 | 668 | ENSG00000099956 | *SMARCB1* | 1636.924 | 131.9541 | 0.7530 | 1.39 |
| 640 | ENSG00000115107 | *STEAP3* | 639.4404 | 58.32045 | 0.7079 | 1.40 | 669 | ENSG00000106683 | *LIMK1* | 1363.418 | 33.17435 | 0.6734 | 1.38 |
| 641 | ENSG00000163161 | *ERCC3* | 1131.03 | 109.5748 | 0.7415 | 1.40 | 670 | ENSG00000227124 | *ZNF717* | 193.7024 | 18.59788 | 0.6229 | 1.38 |
| 642 | ENSG00000160049 | *DFFA* | 2332.338 | 109.3658 | 0.7416 | 1.40 | 671 | ENSG00000185104 | *FAF1* | 1368.003 | 26.68959 | 0.6562 | 1.38 |
| 643 | ENSG00000142327 | *RNPEPL1* | 803.0536 | 21.75264 | 0.6344 | 1.40 | 672 | ENSG00000186642 | *PDE2A* | 2556.096 | 154.8884 | 0.7609 | 1.38 |
| 644 | ENSG00000184575 | *XPOT* | 5566.512 | 213.0214 | 0.7700 | 1.40 | 673 | ENSG00000159692 | *CTBP1* | 3007.059 | 200.372 | 0.7711 | 1.38 |
| 645 | ENSG00000187601 | *MAGEH1* | 554.6384 | 32.1456 | 0.6678 | 1.40 | 674 | ENSG00000090615 | *GOLGA3* | 2651.952 | 94.31258 | 0.7387 | 1.38 |
| 646 | ENSG00000138073 | *PREB* | 815.077 | 79.10587 | 0.7258 | 1.40 | 675 | ENSG00000047056 | *WDR37* | 1331.05 | 107.0078 | 0.7449 | 1.38 |
| 647 | ENSG00000154930 | *ACSS1* | 472.3773 | 39.51635 | 0.6833 | 1.40 | 676 | ENSG00000125648 | *SLC25A23* | 565.6071 | 11.42566 | 0.5684 | 1.38 |
| 648 | ENSG00000142669 | *SH3BGRL3* | 2835.121 | 96.99767 | 0.7363 | 1.40 | 677 | ENSG00000167986 | *DDB1* | 10440.54 | 141.304 | 0.7575 | 1.37 |
| 649 | ENSG00000170515 | *PA2G4* | 3177.675 | 140.5265 | 0.7534 | 1.40 | 678 | ENSG00000068400 | *GRIPAP1* | 930.0872 | 69.51344 | 0.7232 | 1.37 |
| 650 | ENSG00000037474 | *NSUN2* | 2763.068 | 97.12842 | 0.7365 | 1.40 | 679 | ENSG00000012660 | *ELOVL5* | 5271.958 | 52.96717 | 0.7074 | 1.37 |
| 651 | ENSG00000105329 | *TGFB1* | 1080 | 28.98991 | 0.6601 | 1.40 | 680 | ENSG00000151835 | *SACS* | 12021.01 | 714.8978 | 0.8114 | 1.37 |
| 652 | ENSG00000183386 | *FHL3* | 355.1477 | 28.05417 | 0.6576 | 1.40 | 681 | ENSG00000136950 | *ARPC5L* | 1417.343 | 34.6696 | 0.6793 | 1.37 |
| 653 | ENSG00000122863 | *CHST3* | 2108.591 | 115.7371 | 0.7453 | 1.40 | 682 | ENSG00000188229 | *TUBB4B* | 5611.215 | 110.5192 | 0.7473 | 1.37 |
| 654 | ENSG00000086015 | *MAST2* | 899.1025 | 35.51527 | 0.6763 | 1.40 | 683 | ENSG00000167460 | *TPM4* | 33685.9 | 346.682 | 0.7912 | 1.37 |
| 655 | ENSG00000112659 | *CUL9* | 1012.9 | 56.15384 | 0.7073 | 1.39 | 684 | ENSG00000048828 | *FAM120A* | 4967.001 | 123.2402 | 0.7525 | 1.37 |
| 656 | ENSG00000168067 | *MAP4K2* | 1018.128 | 24.26341 | 0.6461 | 1.39 | 685 | ENSG00000108312 | *UBTF* | 3752.868 | 162.7845 | 0.7643 | 1.37 |
| 657 | ENSG00000173846 | *PLK3* | 1046.778 | 24.09529 | 0.6458 | 1.39 | 686 | ENSG00000165637 | *VDAC2* | 4605.912 | 266.7627 | 0.7831 | 1.37 |
| 658 | ENSG00000198380 | *GFPT1* | 3609.982 | 70.23389 | 0.7213 | 1.39 | 687 | ENSG00000144524 | *COPS7B* | 516.1912 | 48.16599 | 0.7031 | 1.36 |
| 659 | ENSG00000226950 | *DANCR* | 621.7518 | 57.88752 | 0.7102 | 1.39 | 688 | ENSG00000172059 | *KLF11* | 1062.581 | 58.7779 | 0.7153 | 1.36 |
| 660 | ENSG00000141867 | *BRD4* | 748.1792 | 30.61181 | 0.6665 | 1.39 | 689 | ENSG00000100201 | *DDX17* | 18238.9 | 1042.116 | 0.8218 | 1.36 |
| 661 | ENSG00000174738 | *NR1D2* | 2885.984 | 140.1085 | 0.7552 | 1.39 | 690 | ENSG00000068438 | *FTSJ1* | 945.8252 | 32.846 | 0.6768 | 1.36 |
| 662 | ENSG00000143543 | *JTB* | 2352.531 | 47.13703 | 0.6976 | 1.39 | 691 | ENSG00000183495 | *EP400* | 843.9072 | 45.84368 | 0.7003 | 1.36 |
| 663 | ENSG00000128606 | *LRRC17* | 329.4196 | 18.32538 | 0.6208 | 1.39 | 692 | ENSG00000110717 | *NDUFS8* | 1573.558 | 22.70052 | 0.6465 | 1.36 |

| Rank | Ensembl Gene ID | Official gene symbol | Mean control counts* | Standard deviation control counts* | Mean HHT/ mean control | Ranking score |
|---|---|---|---|---|---|---|
| 693 | ENSG00000166532 | *RIMKLB* | 1331.297 | 25.21993 | 1.5239 | 1.36 |
| 694 | ENSG00000091947 | *TMEM101* | 856.4828 | 46.79279 | 0.7022 | 1.36 |
| 695 | ENSG00000113119 | *TMCO6* | 263.5788 | 19.21073 | 0.6320 | 1.36 |
| 696 | ENSG00000185753 | *CXorf38* | 681.6807 | 46.08908 | 0.7019 | 1.36 |
| 697 | ENSG00000273749 | *CYFIP1* | 5406.542 | 92.46856 | 0.7412 | 1.36 |
| 698 | ENSG00000162236 | *STX5* | 978.9081 | 83.11104 | 0.7360 | 1.35 |
| 699 | ENSG00000133812 | *SBF2* | 3163.593 | 33.62827 | 0.6802 | 1.35 |
| 700 | ENSG00000116120 | *FARSB* | 1221.209 | 81.36442 | 0.7350 | 1.35 |
| 701 | ENSG00000166860 | *ZBTB39* | 755.6757 | 17.8698 | 0.6254 | 1.35 |
| 702 | ENSG00000186575 | *NF2* | 5033.358 | 35.45001 | 0.6844 | 1.35 |
| 703 | ENSG00000138448 | *ITGAV* | 18854.56 | 575.4916 | 1.2372 | 1.35 |
| 704 | ENSG00000103423 | *DNAJA3* | 1174.961 | 24.39093 | 0.6548 | 1.35 |
| 705 | ENSG00000127463 | *EMC1* | 2935.735 | 123.8583 | 0.7555 | 1.35 |
| 706 | ENSG00000157193 | *LRP8* | 523.7773 | 0.0299025 | 0.6805 | 1.35 |
| 707 | ENSG00000115970 | *THADA* | 1121.684 | 78.11916 | 0.7334 | 1.35 |
| 708 | ENSG00000069667 | *RORA* | 1578.301 | 60.3822 | 1.3898 | 1.35 |
| 709 | ENSG00000189091 | *SF3B3* | 5314.521 | 123.2485 | 0.7556 | 1.35 |
| 710 | ENSG00000233822 | *HIST1H2BN* | 910.2124 | 65.21873 | 0.7241 | 1.35 |
| 711 | ENSG00000197976 | *AKAP17A* | 1742.029 | 30.86814 | 0.6748 | 1.35 |
| 712 | ENSG00000113719 | *ERGIC1* | 4967.555 | 83.61442 | 0.7373 | 1.35 |
| 713 | ENSG00000172216 | *CEBPB* | 863.29 | 30.5638 | 0.6741 | 1.35 |
| 714 | ENSG00000156171 | *DRAM2* | 1236.092 | 49.09303 | 0.7074 | 1.35 |
| 715 | ENSG00000198816 | *ZNF358* | 840.859 | 63.68905 | 0.7230 | 1.35 |
| 716 | ENSG00000114019 | *AMOTL2* | 8189.596 | 194.6086 | 0.7745 | 1.35 |
| 717 | ENSG00000144724 | *PTPRG* | 1884.286 | 21.88533 | 0.6464 | 1.35 |
| 718 | ENSG00000074356 | *C17orf85* | 1981.785 | 82.3279 | 0.7371 | 1.35 |
| 719 | ENSG00000070081 | *NUCB2* | 2646.962 | 45.16758 | 0.7026 | 1.35 |
| 720 | ENSG00000145390 | *USP53* | 2260.722 | 28.26082 | 0.6688 | 1.34 |
| 721 | ENSG00000133028 | *SCO1* | 1372.734 | 57.87853 | 0.7181 | 1.34 |
| 722 | ENSG00000166326 | *TRIM44* | 4073.59 | 54.3426 | 0.7144 | 1.34 |
| 723 | ENSG00000166272 | *WBP1L* | 2197.427 | 77.08862 | 0.7340 | 1.34 |
| 724 | ENSG00000231925 | *TAPBP* | 2983.95 | 92.75652 | 0.7434 | 1.34 |
| 725 | ENSG00000124789 | *NUP153* | 2658.445 | 196.0909 | 0.7754 | 1.34 |
| 726 | ENSG00000171161 | *ZNF672* | 628.2808 | 58.47471 | 0.7191 | 1.34 |
| 727 | ENSG00000118777 | *ABCG2* | 298.408 | 18.5168 | 0.6317 | 1.34 |
| 728 | ENSG00000135862 | *LAMC1* | 30867.88 | 232.1623 | 0.7819 | 1.34 |
| 729 | ENSG00000197746 | *PSAP* | 40490.5 | 1244.023 | 0.8288 | 1.34 |
| 730 | ENSG00000105700 | *KXD1* | 1992.181 | 93.77254 | 0.7450 | 1.34 |
| 731 | ENSG00000173209 | *AHSA2* | 825.6733 | 31.28073 | 0.6783 | 1.34 |
| 732 | ENSG00000136238 | *RAC1* | 7169.825 | 73.88434 | 0.7331 | 1.34 |
| 733 | ENSG00000121671 | *CRY2* | 427.4748 | 31.00018 | 0.6778 | 1.34 |
| 734 | ENSG00000148248 | *SURF4* | 7919.232 | 78.22238 | 0.7362 | 1.34 |
| 735 | ENSG00000119630 | *PGF* | 7972.051 | 23.85413 | 0.6565 | 1.33 |
| 736 | ENSG00000179889 | *PDXDC1* | 3835.911 | 166.5027 | 0.7707 | 1.33 |
| 737 | ENSG00000138698 | *RAP1GDS1* | 2372.839 | 132.0575 | 0.7614 | 1.33 |
| 738 | ENSG00000177595 | *PIDD1* | 412.0577 | 13.52033 | 0.6001 | 1.33 |
| 739 | ENSG00000100292 | *HMOX1* | 8137.996 | 779.0256 | 0.8190 | 1.33 |
| 740 | ENSG00000142733 | *MAP3K6* | 404.4716 | 34.67557 | 0.6873 | 1.33 |
| 741 | ENSG00000166598 | *HSP90B1* | 24137.15 | 1502.24 | 0.8339 | 1.33 |
| 742 | ENSG00000106105 | *GARS* | 6015.256 | 197.1248 | 0.7777 | 1.33 |
| 743 | ENSG00000167972 | *ABCA3* | 1581.488 | 148.4705 | 0.7667 | 1.33 |
| 744 | ENSG00000179218 | *CALR* | 28280.57 | 859.1922 | 0.8217 | 1.33 |
| 745 | ENSG00000142507 | *PSMB6* | 2467.05 | 208.8153 | 0.7801 | 1.33 |
| 746 | ENSG00000104361 | *NIPAL2* | 452.6396 | 27.67996 | 1.4909 | 1.33 |
| 747 | ENSG00000102144 | *PGK1* | 10582.5 | 383.867 | 0.8004 | 1.32 |
| 748 | ENSG00000169594 | *BNC1* | 1160.403 | 79.56353 | 0.7389 | 1.32 |
| 749 | ENSG00000141858 | *SAMD1* | 771.7346 | 37.98523 | 0.6949 | 1.32 |
| 750 | ENSG00000157911 | *PEX10* | 563.6745 | 33.39617 | 0.6861 | 1.32 |

| Rank | Ensembl Gene ID | Official gene symbol | Mean control counts* | Standard deviation control counts* | Mean HHT/ mean control | Ranking score | Rank | Ensembl Gene ID | Official gene symbol | Mean control counts* | Standard deviation control counts* | Mean HHT/ mean control | Ranking score |
|---|---|---|---|---|---|---|---|---|---|---|---|---|---|
| 751 | ENSG00000185324 | *CDK10* | 799.3279 | 9.455352 | 0.5552 | 1.32 | 780 | ENSG00000145555 | *MYO10* | 7574.255 | 221.6821 | 0.7854 | 1.30 |
| 752 | ENSG00000141568 | *FOXK2* | 1468.875 | 43.97235 | 0.7052 | 1.32 | 781 | ENSG00000118508 | *RAB32* | 3047.791 | 145.5737 | 0.7695 | 1.30 |
| 753 | ENSG00000100592 | *DAAM1* | 1137.914 | 65.88132 | 0.7294 | 1.32 | 782 | ENSG00000167562 | *ZNF701* | 242.9725 | 19.53125 | 0.6452 | 1.30 |
| 754 | ENSG00000149499 | *EML3* | 1327.756 | 30.29554 | 0.6789 | 1.32 | 783 | ENSG00000131435 | *PDLIM4* | 1915.484 | 18.45082 | 0.6397 | 1.30 |
| 755 | ENSG00000165271 | *NOL6* | 1448.837 | 69.3883 | 0.7324 | 1.32 | 784 | ENSG00000221869 | *CEBPD* | 365.9793 | 4.089559 | 0.3967 | 1.30 |
| 756 | ENSG00000176225 | *RTTN* | 659.748 | 26.60003 | 0.6689 | 1.32 | 785 | ENSG00000125124 | *BBS2* | 1505.395 | 62.4742 | 0.7301 | 1.30 |
| 757 | ENSG00000070214 | *SLC44A1* | 3006.834 | 117.9146 | 0.7585 | 1.32 | 786 | ENSG00000131446 | *MGAT1* | 3485.925 | 165.7088 | 0.7753 | 1.30 |
| 758 | ENSG00000101363 | *MANBAL* | 351.166 | 32.244 | 0.6844 | 1.32 | 787 | ENSG00000198677 | *TTC37* | 8012.375 | 118.3477 | 0.7616 | 1.30 |
| 759 | ENSG00000167202 | *TBC1D2B* | 2242.834 | 112.4831 | 0.7566 | 1.32 | 788 | ENSG00000156860 | *FBRS* | 755.9 | 64.58762 | 0.7322 | 1.30 |
| 760 | ENSG00000131873 | *CHSY1* | 2844.159 | 44.93246 | 0.7077 | 1.32 | 789 | ENSG00000169398 | *PTK2* | 6085.823 | 58.41893 | 0.7266 | 1.30 |
| 761 | ENSG00000052841 | *TTC17* | 3232.327 | 33.3138 | 0.6872 | 1.32 | 790 | ENSG00000168994 | *PXDC1* | 5058.343 | 157.2475 | 0.7735 | 1.30 |
| 762 | ENSG00000064545 | *TMEM161A* | 540.6174 | 26.59155 | 0.6698 | 1.31 | 791 | ENSG00000180198 | *RCC1* | 1113.748 | 89.34284 | 0.7489 | 1.30 |
| 763 | ENSG00000189190 | *ZNF600* | 285.1501 | 14.64324 | 0.6128 | 1.31 | 792 | ENSG00000196295 | | 573.7993 | 27.22014 | 0.6750 | 1.30 |
| 764 | ENSG00000058091 | *CDK14* | 849.1075 | 11.10925 | 0.5794 | 1.31 | 793 | ENSG00000101337 | *TM9SF4* | 1759.836 | 106.5584 | 0.7573 | 1.30 |
| 765 | ENSG00000148296 | *SURF6* | 709.5434 | 29.79166 | 0.6791 | 1.31 | 794 | ENSG00000122729 | *ACO1* | 4133.712 | 234.4772 | 0.7885 | 1.30 |
| 766 | ENSG00000092820 | *EZR* | 3518.398 | 184.6576 | 0.7776 | 1.31 | 795 | ENSG00000095319 | *NUP188* | 1753.022 | 23.96763 | 0.6648 | 1.30 |
| 767 | ENSG00000197555 | *SIPA1L1* | 708.8073 | 58.21278 | 0.7240 | 1.31 | 796 | ENSG00000136878 | *USP20* | 479.2273 | 19.74157 | 0.6476 | 1.30 |
| 768 | ENSG00000164054 | *SHISA5* | 4183.121 | 112.6922 | 0.7575 | 1.31 | 797 | ENSG00000082212 | *ME2* | 1875.237 | 22.05676 | 0.6578 | 1.30 |
| 769 | ENSG00000148498 | *PARD3* | 1389.887 | 54.76489 | 0.7206 | 1.31 | 798 | ENSG00000112941 | *PAPD7* | 866.9127 | 33.48372 | 0.6915 | 1.30 |
| 770 | ENSG00000145741 | *BTF3* | 13673.31 | 286.2169 | 0.7932 | 1.31 | 799 | ENSG00000169032 | *MAP2K1* | 1831.228 | 107.9108 | 0.7583 | 1.30 |
| 771 | ENSG00000186866 | *POFUT2* | 1396.052 | 15.78348 | 0.6219 | 1.31 | 800 | ENSG00000197081 | *IGF2R* | 4408.729 | 143.232 | 0.7704 | 1.29 |
| 772 | ENSG00000081760 | *AACS* | 847.6599 | 17.47971 | 0.6326 | 1.31 | 801 | ENSG00000023191 | *RNH1* | 4979.042 | 144.5322 | 0.7708 | 1.29 |
| 773 | ENSG00000173757 | *STAT5B* | 1985.282 | 26.94577 | 0.6721 | 1.31 | 802 | ENSG00000132471 | *WBP2* | 1227.153 | 15.42738 | 0.6231 | 1.29 |
| 774 | ENSG00000278259 | *MYO19* | 1203.624 | 25.64481 | 0.6681 | 1.31 | 803 | ENSG00000005007 | *UPF1* | 1872.75 | 139.976 | 0.7696 | 1.29 |
| 775 | ENSG00000125826 | *RBCK1* | 1663.471 | 143.3631 | 0.7685 | 1.31 | 804 | ENSG00000244462 | *RBM12* | 3575.48 | 210.2193 | 0.7852 | 1.29 |
| 776 | ENSG00000263528 | *IKBKE* | 227.0975 | 11.33718 | 1.7131 | 1.31 | 805 | ENSG00000077782 | *FGFR1* | 2659.399 | 74.73497 | 0.7411 | 1.29 |
| 777 | ENSG00000257923 | *CUX1* | 1539.894 | 24.19542 | 0.6635 | 1.31 | 806 | ENSG00000276368 | *HIST1H2AJ* | 274.8256 | 13.98741 | 1.6318 | 1.29 |
| 778 | ENSG00000104231 | *ZFAND1* | 1054.55 | 57.50828 | 0.7245 | 1.31 | 807 | ENSG00000074696 | *HACD3* | 3422.068 | 167.6294 | 0.7771 | 1.29 |
| 779 | ENSG00000160410 | *SHKBP1* | 1057.419 | 73.09322 | 0.7378 | 1.31 | 808 | ENSG00000165097 | *KDM1B* | 490.288 | 19.9511 | 0.6496 | 1.29 |

| Rank | Ensembl Gene ID | Official gene symbol | Mean control counts* | Standard deviation control counts* | Mean HHT/ mean control | Ranking score |
|---|---|---|---|---|---|---|
| 809 | ENSG00000197905 | *TEAD4* | 827.9625 | 45.45057 | 0.7130 | 1.29 |
| 810 | ENSG00000162139 | *NEU3* | 442.1716 | 34.67887 | 0.6950 | 1.29 |
| 811 | ENSG00000126453 | *BCL2L12* | 311.776 | 13.01501 | 0.6048 | 1.29 |
| 812 | ENSG00000118894 | *EEF2KMT* | 289.291 | 25.79828 | 0.6724 | 1.29 |
| 813 | ENSG00000163444 | *TMEM183A* | 1900.475 | 170.1518 | 0.7785 | 1.29 |
| 814 | ENSG00000114796 | *KLHL24* | 2367.439 | 65.33741 | 1.3600 | 1.29 |
| 815 | ENSG00000087088 | *BAX* | 2289.386 | 27.00694 | 0.6775 | 1.28 |
| 816 | ENSG00000196628 | *TCF4* | 11520.42 | 813.9514 | 1.2109 | 1.28 |
| 817 | ENSG00000110888 | *CAPRIN2* | 1006.587 | 79.49316 | 0.7461 | 1.28 |
| 818 | ENSG00000081923 | *ATP8B1* | 5670.54 | 109.3954 | 0.7612 | 1.28 |
| 819 | ENSG00000111275 | *ALDH2* | 1622.851 | 80.15367 | 0.7466 | 1.28 |
| 820 | ENSG00000241685 | *ARPC1A* | 4614.659 | 179.1226 | 0.7813 | 1.28 |
| 821 | ENSG00000203879 | *GDI1* | 2213.425 | 19.89367 | 0.6518 | 1.28 |
| 822 | ENSG00000244879 | *GABPB1-AS1* | 385.8495 | 32.19016 | 1.4452 | 1.28 |
| 823 | ENSG00000172716 | *SLFN11* | 3369.328 | 110.3366 | 0.7621 | 1.28 |
| 824 | ENSG00000105497 | *ZNF175* | 949.9819 | 47.14227 | 0.7177 | 1.28 |
| 825 | ENSG00000246451 | | 42.72751 | 0.0985479 | 1.7338 | 1.28 |
| 826 | ENSG00000142208 | *AKT1* | 3050.724 | 207.3644 | 0.7874 | 1.27 |
| 827 | ENSG00000106635 | *BCL7B* | 1229.299 | 29.68556 | 0.6867 | 1.27 |
| 828 | ENSG00000086696 | *HSD17B2* | 199.0779 | 2.349552 | 4.4408 | 1.27 |
| 829 | ENSG00000165915 | *SLC39A13* | 1591.155 | 136.2025 | 0.7718 | 1.27 |
| 830 | ENSG00000097007 | *ABL1* | 3634.662 | 118.106 | 0.7659 | 1.27 |
| 831 | ENSG00000101407 | *TTI1* | 727.8669 | 52.3229 | 0.7250 | 1.27 |
| 832 | ENSG00000072682 | *P4HA2* | 2210.451 | 77.41833 | 0.7463 | 1.27 |
| 833 | ENSG00000068724 | *TTC7A* | 502.3137 | 15.91365 | 0.6315 | 1.27 |
| 834 | ENSG00000180530 | *NRIP1* | 6493.854 | 380.2819 | 0.8074 | 1.27 |
| 835 | ENSG00000066583 | *ISOC1* | 451.6744 | 23.43298 | 0.6684 | 1.27 |
| 836 | ENSG00000197448 | *GSTK1* | 2294.742 | 113.8481 | 0.7649 | 1.27 |
| 837 | ENSG00000067182 | *TNFRSF1A* | 2160.12 | 17.4621 | 0.6418 | 1.27 |
| 838 | ENSG00000180185 | *FAHD1* | 1328.503 | 110.6104 | 0.7640 | 1.27 |
| 839 | ENSG00000253368 | *TRNP1* | 1089.492 | 22.77183 | 0.6667 | 1.27 |
| 840 | ENSG00000130204 | *TOMM40* | 1072.234 | 47.93188 | 0.7209 | 1.27 |
| 841 | ENSG00000172500 | *FIBP* | 1687.578 | 92.43414 | 0.7560 | 1.27 |
| 842 | ENSG00000112658 | *SRF* | 1357.007 | 73.10487 | 0.7447 | 1.27 |
| 843 | ENSG00000155380 | *SLC16A1* | 4999.743 | 134.8987 | 0.7726 | 1.27 |
| 844 | ENSG00000178605 | *GTPBP6* | 1161.296 | 38.42506 | 0.7071 | 1.26 |
| 845 | ENSG00000187446 | *CHP1* | 3815.66 | 273.2026 | 0.7982 | 1.26 |
| 846 | ENSG00000152642 | *GPD1L* | 876.3752 | 28.89416 | 0.6867 | 1.26 |
| 847 | ENSG00000011485 | *PPP5C* | 1498.798 | 68.99706 | 0.7421 | 1.26 |
| 848 | ENSG00000047230 | *CTPS2* | 1151.993 | 12.29887 | 0.6051 | 1.26 |
| 849 | ENSG00000160445 | *ZER1* | 963.8032 | 20.39079 | 0.6583 | 1.26 |
| 850 | ENSG00000143418 | *CERS2* | 2878.719 | 111.1002 | 0.7653 | 1.26 |
| 851 | ENSG00000011405 | *PIK3C2A* | 8595.629 | 313.4528 | 0.8032 | 1.26 |
| 852 | ENSG00000135905 | *DOCK10* | 806.2341 | 48.68439 | 1.3828 | 1.26 |
| 853 | ENSG00000077092 | *RARB* | 933.53 | 8.443277 | 0.5543 | 1.26 |
| 854 | ENSG00000198931 | *APRT* | 2301.493 | 22.51234 | 0.6677 | 1.26 |
| 855 | ENSG00000198324 | *FAM109A* | 933.7836 | 17.85435 | 0.6466 | 1.26 |
| 856 | ENSG00000171444 | *MCC* | 2108.955 | 77.34286 | 0.7495 | 1.25 |
| 857 | ENSG00000172366 | *FAM195A* | 425.6793 | 8.286147 | 0.5527 | 1.25 |
| 858 | ENSG00000104859 | *CLASRP* | 439.1954 | 38.88775 | 0.7101 | 1.25 |
| 859 | ENSG00000189241 | *TSPYL1* | 2633.453 | 160.5317 | 0.7813 | 1.25 |
| 860 | ENSG00000182165 | *TP53TG1* | 457.5863 | 36.11677 | 0.7052 | 1.25 |
| 861 | ENSG00000142279 | *WTIP* | 1182.275 | 105.5609 | 0.7643 | 1.25 |
| 862 | ENSG00000001461 | *NIPAL3* | 750.435 | 48.33822 | 0.7241 | 1.25 |
| 863 | ENSG00000167930 | *ITFG3* | 704.0825 | 69.12363 | 0.7441 | 1.25 |
| 864 | ENSG00000072501 | *SMC1A* | 1656.455 | 71.18875 | 0.7458 | 1.25 |
| 865 | ENSG00000130856 | *ZNF236* | 611.9795 | 28.21583 | 0.6877 | 1.25 |
| 866 | ENSG00000164323 | *CFAP97* | 1500.284 | 124.4173 | 1.2958 | 1.25 |

| Rank | Ensembl Gene ID | Official gene symbol | Mean control counts* | Standard deviation control counts* | Mean HHT/ mean control | Ranking score |
|---|---|---|---|---|---|---|
| 867 | ENSG00000122378 | *FAM213A* | 9966.105 | 503.5972 | 0.8180 | 1.25 |
| 868 | ENSG00000163762 | *TM4SF18* | 3863.714 | 188.4089 | 1.2694 | 1.25 |
| 869 | ENSG00000168894 | *RNF181* | 2129.687 | 41.76627 | 0.7156 | 1.25 |
| 870 | ENSG00000128805 | *ARHGAP22* | 1383.997 | 36.33888 | 0.7066 | 1.25 |
| 871 | ENSG00000135597 | *REPS1* | 1202.293 | 61.81822 | 0.7390 | 1.25 |
| 872 | ENSG00000143624 | *INTS3* | 1216.941 | 104.8044 | 0.7648 | 1.25 |
| 873 | ENSG00000076108 | *BAZ2A* | 1547.485 | 82.6311 | 0.7539 | 1.25 |
| 874 | ENSG00000198604 | *BAZ1A* | 3863.506 | 267.2591 | 0.8002 | 1.25 |
| 875 | ENSG00000065882 | *TBC1D1* | 3694.779 | 86.39942 | 0.7565 | 1.24 |
| 876 | ENSG00000106261 | *ZKSCAN1* | 3337.248 | 47.67635 | 0.7252 | 1.24 |
| 877 | ENSG00000105711 | *SCN1B* | 479.8557 | 20.05561 | 0.6612 | 1.24 |
| 878 | ENSG00000103365 | *GGA2* | 1607.961 | 46.1003 | 0.7234 | 1.24 |
| 879 | ENSG00000135521 | *LTV1* | 672.1667 | 63.91655 | 0.7421 | 1.24 |
| 880 | ENSG00000145414 | *NAF1* | 268.539 | 12.19593 | 0.6091 | 1.24 |
| 881 | ENSG00000214078 | *CPNE1* | 2225.862 | 100.0089 | 0.7642 | 1.24 |
| 882 | ENSG00000105520 | | 434.4865 | 33.71086 | 0.7034 | 1.24 |
| 883 | ENSG00000167280 | *ENGASE* | 360.0944 | 19.61736 | 0.6601 | 1.24 |
| 884 | ENSG00000115486 | *GGCX* | 1417.085 | 45.66434 | 0.7238 | 1.24 |
| 885 | ENSG00000116809 | *ZBTB17* | 940.1019 | 65.4379 | 0.7444 | 1.23 |
| 886 | ENSG00000089916 | *GPATCH2L* | 2872.794 | 218.6343 | 0.7952 | 1.23 |
| 887 | ENSG00000179776 | *CDH5* | 28020.21 | 1101.361 | 0.8385 | 1.23 |
| 888 | ENSG00000143507 | *DUSP10* | 109.1586 | 8.465775 | 1.7812 | 1.23 |
| 889 | ENSG00000168495 | *POLR3D* | 1036.79 | 31.01484 | 0.6985 | 1.23 |
| 890 | ENSG00000149257 | *SERPINH1* | 8509.883 | 80.1702 | 0.7550 | 1.23 |
| 891 | ENSG00000100418 | *DESI1* | 1781.91 | 57.23592 | 0.7378 | 1.23 |
| 892 | ENSG00000158941 | *CCAR2* | 2122.927 | 35.13646 | 0.7078 | 1.23 |
| 893 | ENSG00000129103 | *SUMF2* | 2234.911 | 99.83745 | 0.7656 | 1.23 |
| 894 | ENSG00000215301 | *DDX3X* | 23806.01 | 81.32214 | 0.7563 | 1.23 |
| 895 | ENSG00000127511 | *SIN3B* | 1655.041 | 78.12212 | 0.7545 | 1.23 |
| 896 | ENSG00000175634 | *RPS6KB2* | 657.1332 | 37.31271 | 0.7123 | 1.23 |
| 897 | ENSG00000185787 | *MORF4L1* | 6331.712 | 226.5786 | 0.7976 | 1.23 |
| 898 | ENSG00000187498 | *COL4A1* | 36661.58 | 354.0307 | 1.2323 | 1.23 |
| 899 | ENSG00000110063 | *DCPS* | 317.2861 | 10.98676 | 0.5996 | 1.23 |
| 900 | ENSG00000150990 | *DHX37* | 685.5593 | 44.97659 | 0.7249 | 1.22 |
| 901 | ENSG00000130402 | *ACTN4* | 11115.58 | 198.1819 | 0.7934 | 1.22 |
| 902 | ENSG00000162129 | *CLPB* | 973.2679 | 11.3287 | 0.6040 | 1.22 |
| 903 | ENSG00000105810 | *CDK6* | 2196.592 | 165.2531 | 1.2708 | 1.22 |
| 904 | ENSG00000130669 | *PAK4* | 518.642 | 11.50121 | 0.6059 | 1.22 |
| 905 | ENSG00000139318 | *DUSP6* | 2032.155 | 93.23438 | 0.7636 | 1.22 |
| 906 | ENSG00000154553 | *PDLIM3* | 1119.186 | 26.42701 | 0.6883 | 1.22 |
| 907 | ENSG00000103512 | *NOMO1* | 2522.939 | 112.484 | 0.7719 | 1.22 |
| 908 | ENSG00000144063 | *MALL* | 271.0864 | 15.79857 | 0.6422 | 1.22 |
| 909 | ENSG00000170540 | *ARL6IP1* | 3728.87 | 149.0216 | 1.2765 | 1.22 |
| 910 | ENSG00000144642 | *RBMS3* | 1900.61 | 155.9318 | 0.7852 | 1.22 |
| 911 | ENSG00000175274 | *TP53I11* | 1122.041 | 43.43397 | 0.7234 | 1.22 |
| 912 | ENSG00000132275 | *RRP8* | 692.7009 | 50.14558 | 0.7321 | 1.22 |
| 913 | ENSG00000108395 | *TRIM37* | 2700.878 | 100.2516 | 0.7673 | 1.22 |
| 914 | ENSG00000169291 | *SHE* | 4949.657 | 381.0999 | 0.8144 | 1.22 |
| 915 | ENSG00000133612 | *AGAP3* | 2027.982 | 24.2003 | 0.6819 | 1.22 |
| 916 | ENSG00000160917 | *CPSF4* | 643.35 | 59.61099 | 0.7420 | 1.22 |
| 917 | ENSG00000168056 | *LTBP3* | 890.8699 | 15.45477 | 0.6405 | 1.22 |
| 918 | ENSG00000125817 | *CENPB* | 2315.617 | 176.505 | 0.7900 | 1.22 |
| 919 | ENSG00000090060 | *PAPOLA* | 3757.405 | 263.1165 | 0.8037 | 1.22 |
| 920 | ENSG00000155016 | *CYP2U1* | 813.3489 | 49.28179 | 0.7320 | 1.22 |
| 921 | ENSG00000159461 | *AMFR* | 3711.413 | 187.7397 | 0.7929 | 1.21 |
| 922 | ENSG00000143612 | *C1orf43* | 3824.156 | 66.17661 | 0.7486 | 1.21 |
| 923 | ENSG00000141385 | *AFG3L2* | 2051.601 | 82.69463 | 0.7596 | 1.21 |

| Rank | Ensembl Gene ID | Official gene symbol | Mean control counts* | Standard deviation control counts* | Mean HHT/ mean control | Ranking score |
|---|---|---|---|---|---|---|
| 924 | ENSG00000099917 | *MED15* | 956.7488 | 34.48034 | 0.7098 | 1.21 |
| 925 | ENSG00000143374 | *TARS2* | 544.2219 | 17.41452 | 0.6541 | 1.21 |
| 926 | ENSG00000025708 | *TYMP* | 145.2224 | 5.068569 | 2.1113 | 1.21 |
| 927 | ENSG00000197498 | *RPF2* | 872.369 | 25.76744 | 0.6886 | 1.21 |
| 928 | ENSG00000189159 | *HN1* | 4134.188 | 78.07533 | 0.7572 | 1.21 |
| 929 | ENSG00000169762 | *TAPT1* | 409.443 | 24.23301 | 1.4619 | 1.21 |
| 930 | ENSG00000215447 | | 301.961 | 14.89511 | 0.6388 | 1.21 |
| 931 | ENSG00000163902 | *RPN1* | 9606.687 | 398.7983 | 0.8171 | 1.21 |
| 932 | ENSG00000100360 | *IFT27* | 624.6224 | 20.15668 | 0.6685 | 1.21 |
| 933 | ENSG00000131389 | *SLC6A6* | 2297.004 | 34.27275 | 0.7103 | 1.21 |
| 934 | ENSG00000124145 | *SDC4* | 2304.839 | 146.8523 | 0.7848 | 1.21 |
| 935 | ENSG00000183207 | *RUVBL2* | 1942.94 | 38.54611 | 0.7185 | 1.21 |
| 936 | ENSG00000133226 | *SRRM1* | 1440.815 | 34.55165 | 0.7113 | 1.21 |
| 937 | ENSG00000168397 | *ATG4B* | 1442.346 | 20.92572 | 0.6725 | 1.21 |
| 938 | ENSG00000132591 | *ERAL1* | 616.0419 | 23.87361 | 0.6837 | 1.21 |
| 939 | ENSG00000101294 | *HM13* | 2360.58 | 59.64693 | 0.7445 | 1.21 |
| 940 | ENSG00000164741 | *DLC1* | 8499.729 | 36.60592 | 0.7153 | 1.21 |
| 941 | ENSG00000100968 | *NFATC4* | 1453.916 | 21.85578 | 0.6764 | 1.21 |
| 942 | ENSG00000104983 | *CCDC61* | 202.7116 | 14.50329 | 0.6370 | 1.21 |
| 943 | ENSG00000181789 | *COPG1* | 6453.585 | 38.37244 | 0.7186 | 1.21 |
| 944 | ENSG00000240972 | *MIF* | 1100.059 | 76.99863 | 0.7578 | 1.20 |
| 945 | ENSG00000170545 | *SMAGP* | 527.0743 | 23.42637 | 0.6828 | 1.20 |
| 946 | ENSG00000109107 | *ALDOC* | 312.6872 | 3.079999 | 0.3431 | 1.20 |
| 947 | ENSG00000135269 | *TES* | 1255.318 | 35.93304 | 1.3991 | 1.20 |
| 948 | ENSG00000112559 | *MDFI* | 1011.033 | 34.87374 | 0.7128 | 1.20 |
| 949 | ENSG00000136869 | *TLR4* | 3784.097 | 368.3471 | 1.2254 | 1.20 |
| 950 | ENSG00000072134 | *EPN2* | 1073.444 | 26.73308 | 0.6939 | 1.20 |
| 951 | ENSG00000081189 | *MEF2C* | 1402.444 | 52.33958 | 1.3544 | 1.20 |
| 952 | ENSG00000197622 | *CDC42SE1* | 4097.655 | 60.99548 | 0.7467 | 1.20 |

| Rank | Ensembl Gene ID | Official gene symbol | Mean control counts* | Standard deviation control counts* | Mean HHT/ mean control | Ranking score |
|---|---|---|---|---|---|---|
| 953 | ENSG00000198722 | *UNC13B* | 3569.414 | 54.65226 | 0.7409 | 1.20 |
| 954 | ENSG00000205339 | *IPO7* | 9315.042 | 68.40971 | 0.7529 | 1.20 |
| 955 | ENSG00000100241 | *SBF1* | 2261.394 | 10.4762 | 0.6005 | 1.20 |
| 956 | ENSG00000134954 | *ETS1* | 5501.796 | 65.83798 | 0.7512 | 1.20 |
| 957 | ENSG00000123992 | *DNPEP* | 1019.6 | 32.57882 | 0.7094 | 1.20 |
| 958 | ENSG00000211584 | *SLC48A1* | 672.0004 | 59.77912 | 0.7465 | 1.20 |
| 959 | ENSG00000132128 | *LRRC41* | 2271.828 | 82.84631 | 0.7629 | 1.20 |
| 960 | ENSG00000183726 | *TMEM50A* | 2591.948 | 84.54326 | 1.3092 | 1.20 |
| 961 | ENSG00000170089 | | 94.04902 | 5.336035 | 2.0418 | 1.20 |
| 962 | ENSG00000036257 | *CUL3* | 2596.814 | 101.5121 | 0.7721 | 1.20 |
| 963 | ENSG00000105971 | *CAV2* | 10537.39 | 688.9767 | 0.8329 | 1.20 |
| 964 | ENSG00000174749 | *C4orf32* | 1796.95 | 114.532 | 1.2865 | 1.19 |
| 965 | ENSG00000164182 | *NDUFAF2* | 590.0491 | 44.58901 | 0.7302 | 1.19 |
| 966 | ENSG00000115520 | *COQ10B* | 2147.458 | 98.65063 | 0.7710 | 1.19 |
| 967 | ENSG00000179632 | *MAF1* | 2732.624 | 145.1463 | 0.7867 | 1.19 |
| 968 | ENSG00000196363 | *WDR5* | 1047.665 | 64.63205 | 0.7510 | 1.19 |
| 969 | ENSG00000162664 | *ZNF326* | 399.8569 | 27.96899 | 1.4309 | 1.19 |
| 970 | ENSG00000139620 | *KANSL2* | 869.4443 | 17.65143 | 0.6599 | 1.19 |
| 971 | ENSG00000122034 | *GTF3A* | 2091.619 | 103.2628 | 0.7731 | 1.19 |
| 972 | ENSG00000099899 | *TRMT2A* | 667.1884 | 37.12223 | 0.7189 | 1.19 |
| 973 | ENSG00000136383 | *ALPK3* | 3358.742 | 311.0126 | 0.8124 | 1.19 |
| 974 | ENSG00000064666 | *CNN2* | 5278.673 | 405.4347 | 0.8201 | 1.19 |
| 975 | ENSG00000119231 | *SENP5* | 2494.614 | 118.3766 | 0.7793 | 1.19 |
| 976 | ENSG00000162437 | *RAVER2* | 1166.672 | 54.67339 | 0.7427 | 1.19 |
| 977 | ENSG00000114480 | *GBE1* | 5087.51 | 321.725 | 0.8140 | 1.19 |

| Rank | Ensembl Gene ID | Official gene symbol | Mean control counts* | Standard deviation control counts* | Mean HHT/ mean control | Ranking score |
|---|---|---|---|---|---|---|
| 978 | ENSG00000205250 | *E2F4* | 2815.592 | 17.08351 | 0.6580 | 1.19 |
| 979 | ENSG00000120705 | *ETF1* | 7121.312 | 260.3422 | 0.8079 | 1.19 |
| 980 | ENSG00000225697 | *SLC26A6* | 294.2536 | 20.50271 | 0.6752 | 1.19 |
| 981 | ENSG00000182957 | *SPATA13* | 344.7019 | 13.37455 | 0.6330 | 1.19 |
| 982 | ENSG00000137764 | *MAP2K5* | 211.8151 | 20.36276 | 0.6747 | 1.19 |
| 983 | ENSG00000101361 | *NOP56* | 2273.823 | 28.1466 | 0.7012 | 1.18 |
| 984 | ENSG00000153904 | *DDAH1* | 3471.729 | 125.4578 | 0.7827 | 1.18 |
| 985 | ENSG00000146757 | *ZNF92* | 814.4487 | 39.30869 | 0.7244 | 1.18 |
| 986 | ENSG00000162458 | *FBLIM1* | 9617.956 | 217.3113 | 0.8025 | 1.18 |
| 987 | ENSG00000154957 | *ZNF18* | 235.0877 | 16.87283 | 0.6578 | 1.18 |
| 988 | ENSG00000154059 | *IMPACT* | 1616.093 | 103.7406 | 0.7750 | 1.18 |
| 989 | ENSG00000113712 | *CSNK1A1* | 8209.812 | 236.1674 | 0.8053 | 1.18 |
| 990 | ENSG00000164597 | *COG5* | 2163.814 | 123.2215 | 0.7822 | 1.18 |
| 991 | ENSG00000121680 | *PEX16* | 524.9691 | 33.41853 | 0.7139 | 1.18 |
| 992 | ENSG00000168924 | *LETM1* | 2062.917 | 55.46695 | 0.7449 | 1.18 |
| 993 | ENSG00000115275 | *MOGS* | 1919.425 | 26.90667 | 0.6982 | 1.18 |
| 994 | ENSG00000168795 | *ZBTB5* | 768.5565 | 55.10647 | 0.7446 | 1.18 |
| 995 | ENSG00000089154 | *GCN1L1* | 3833.92 | 327.834 | 0.8154 | 1.18 |
| 996 | ENSG00000100034 | *PPM1F* | 6128.27 | 236.5016 | 0.8056 | 1.18 |
| 997 | ENSG00000128159 | *TUBGCP6* | 910.0847 | 80.50854 | 0.7642 | 1.18 |
| 998 | ENSG00000088038 | *CNOT3* | 320.37 | 18.15395 | 0.6657 | 1.18 |
| 999 | ENSG00000139579 | *NABP2* | 880.6104 | 59.38162 | 0.7492 | 1.18 |
| 1000 | ENSG00000150551 | *LYPD1* | 903.3109 | 2.183795 | 0.2211 | 1.18 |

The top 1000-rankeed genes based on normalization by total library read counts only (to provide a complementary analysis to those performed using genes differentially expressed genes following DeSeq2 normalization (as listed in Supplementary Table 3). Counts *, normalized to total library read counts. Control and HHT refer to the untreated samples, and as described in the Supplementary Methods, the ranking score was calculated as the square root of the square of

$$Ln\left(\frac{Mean\ \ ENG,ACVRL\ and\ SMAD4\ untreated\ BOEC\ \ alignment}{Mean\ untreated\ control\ BOEC\ alignment}\right)*\left(-\ln\left(\frac{1}{Standard\ deviation}\right)\right)$$

For gene ontology analyses, the top 100, 200, 300 400, 500, 750 and 1000 (all) of these genes were used in separate clustering analyse
